# Spatial multi-omics reveals targetable immunosuppressive macrophage T-cell interactions in human AML bone marrow

**DOI:** 10.64898/2026.08.07.743431

**Authors:** Merel van der Meulen, Emma S. Pool, Alicia Perzolli, Joost B. Koedijk, Eva Argiro, Li-Ting Chen, Wim J. de Jonge, Elizabeth Schweighart, Marijn A. Vermeulen, Stefan Nierkens, Jana Ihlow, David Horst, Andrej Lissat, H. Josef Vormoor, Mirjam E. Belderbos, Hendrik Veelken, Livius Penter, Bianca F. Goemans, Erik van den Akker, Thanasis Margaritis, C. Michel Zwaan, Marieke Griffioen, Jennifer M.L. Tjon, Olaf Heidenreich

## Abstract

The immunosuppressive bone marrow microenvironment is an important contributor to the limited success of immunotherapy in acute myeloid leukemia (AML), but the cellular interactions underlying AML immune evasion are incompletely understood. We therefore generated a single-cell spatial transcriptomic and proteomic atlas using 148 bone biopsies from 113 individuals comprising pediatric and adult AML at diagnosis and non-leukemic controls. We observed an expansion of regulatory T cells (Tregs) in AML, with stronger colocalization between Tregs and macrophages compared to non-leukemic bone marrow. Distinct cellular neighborhoods were enriched for myeloid progenitor-like cells together with macrophages and T cells, which correlated with higher macrophage and T cell immune checkpoint expression. Moreover, these neighborhoods were associated with specific AML subtypes, especially *KMT2A*-rearranged and *RUNX1*::*RUNX1T1* AML. These spatial patterns were validated by identification of malignant cells via *in situ* fusion detection in *RUNX1*::*RUNX1T1* cases. Functional experiments revealed that macrophages and AML cells not only actively recruit Tregs, but also promote naïve T cell differentiation into Tregs. Spatially informed ligand-receptor analysis predicted the involvement of the Galectin-9 – CD44/TIM-3 axis in this immunosuppressive crosstalk, which was supported by *in vitro* inhibition of CD44 and/or TIM-3 preventing macrophage- and AML-induced Treg differentiation. Collectively, this comprehensive spatial map of the AML bone marrow identified tripartite crosstalk between AML, macrophages, and T cells mediated by the Galectin-9 – CD44/TIM-3 axis as a key component of the immunosuppressive microenvironment. Targeting Galectin-9 – CD44/TIM-3 interactions may be a promising strategy to overcome immune evasion and enhance immunotherapeutic success in AML.

**Highlights:**

- Spatial transcriptomic and proteomic atlas of pediatric and adult acute myeloid leukemia (AML) bone marrow
- Increased colocalization of macrophages and regulatory T cells (Tregs) in AML
- Macrophages and AML cells induce differentiation of naïve T cells to Tregs *in vitro*, which can be prevented by inhibition of CD44 or TIM-3

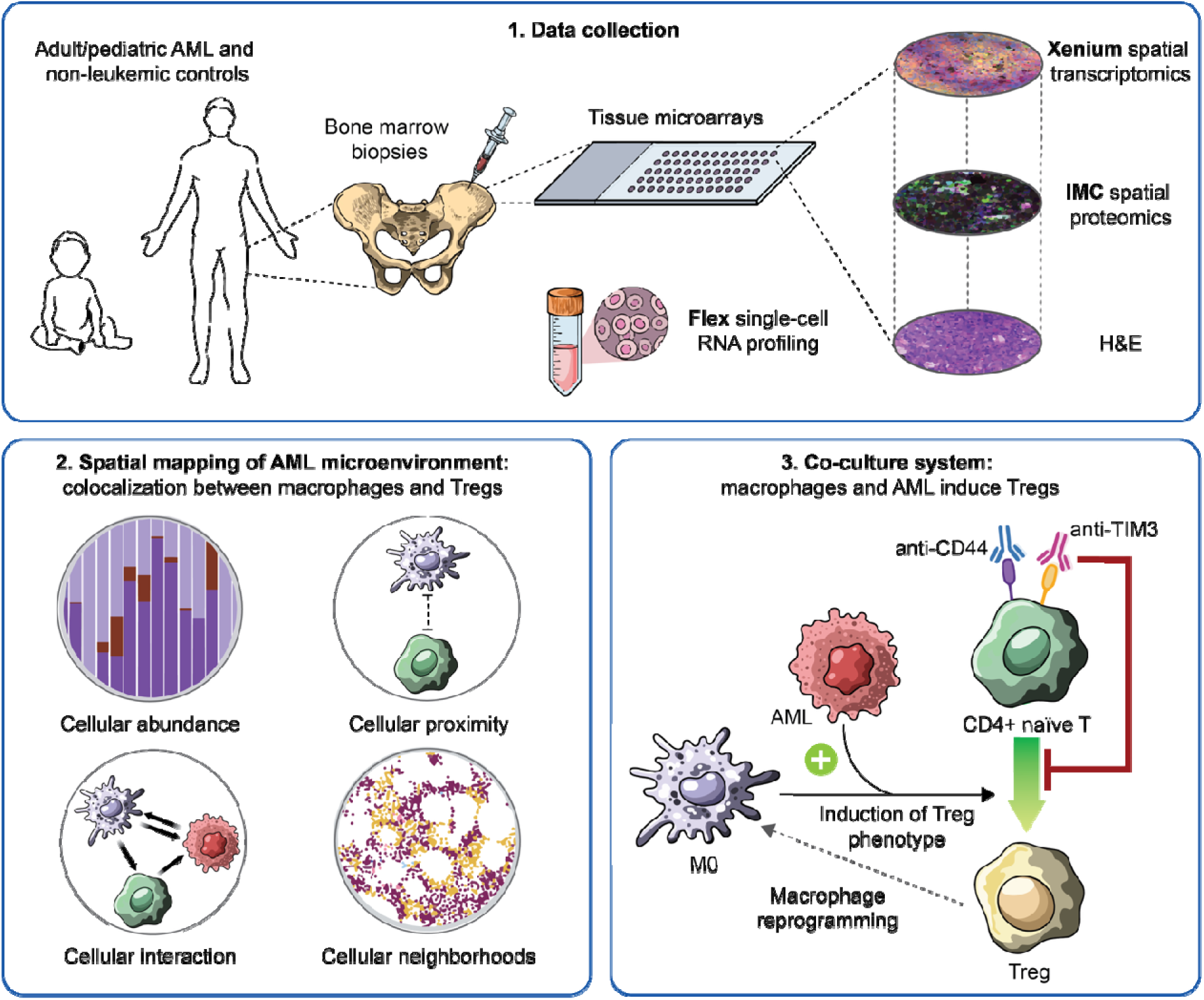

## Introduction

Acute myeloid leukemia (AML) is a heterogeneous and aggressive hematological malignancy with a poor prognosis in both adult and pediatric patients, especially in relapsed disease^1–3^. The poor survival and significant toxicities associated with conventional chemotherapy necessitate the development of novel therapeutic strategies. Based on the graft-versus-leukemia effect of allogeneic hematopoietic stem cell transplantation^4^, targeted immunotherapy is considered a promising strategy^5,6^. However, neither T and natural killer (NK) cell engagers nor AML-directed chimeric antigen receptor (CAR) T/NK therapy have so far yielded significant clinical benefit^6–8^.

The limited efficacy of immunotherapy in AML has been attributed to several factors, including a low mutational burden, scarcity of AML-specific antigens, and, importantly, an immunosuppressive bone marrow (BM) microenvironment^6,8^. Beyond AML-intrinsic mechanisms such as reduced antigen presentation^9,10^ and increased inhibitory signaling^10,11^, several innate and adaptive immune and stromal cells have been implicated in this immunosuppressive state^12–16^. Notably, the AML BM is enriched with immunosuppressive cell types such as regulatory T cells (Tregs)^17–19^ and anti-inflammatory macrophages^12,20^. However, immune profiling has generally relied on BM aspirates, which do not preserve spatial information and incompletely capture various cell types such as neutrophils, macrophages, stromal and endothelial cells^21–23^. Consequently, the spatial organization and cellular interactions underpinning immune evasion within the AML BM remain insufficiently defined. Insight into these interactions is crucial to improve immunotherapeutic strategies.

To address this gap, we performed single-cell spatial transcriptomic and proteomic profiling of 148 pediatric and adult AML bone marrow biopsies. This spatial atlas reveals distinct neighborhoods where AML cells, macrophages, and T cells colocalize. We demonstrate that crosstalk between these three cell types promotes Treg differentiation and immunosuppressive macrophage polarization. Mechanistically, we identify the Galectin-9–CD44/TIM-3 axis as a primary driver of these interactions, showing that inhibition of CD44 or TIM-3 prevents Treg induction. Collectively, our findings map the AML microenvironment at unprecedented resolution and highlight the CD44/TIM-3 axis as a therapeutic target to overcome immune evasion and enhance immunotherapy.

## Methods

BM biopsies were arranged in three tissue microarrays (TMAs). Spatial transcriptomics was performed on all three TMAs (n=148 biopsies) using 10x Genomics Xenium with a 477-gene panel (Figure 1A, Table S1). Spatial proteomics was performed on serial sections of two of the three TMAs (n=104 biopsies) using 37-plex imaging mass cytometry (IMC)^24^. Near whole-transcriptome profiling was performed on 4 formalin-fixed paraffin-embedded (FFPE) BM biopsies using 10x Flex single-cell RNA profiling capturing ∼18,000 protein-coding genes.

**Figure 1.**
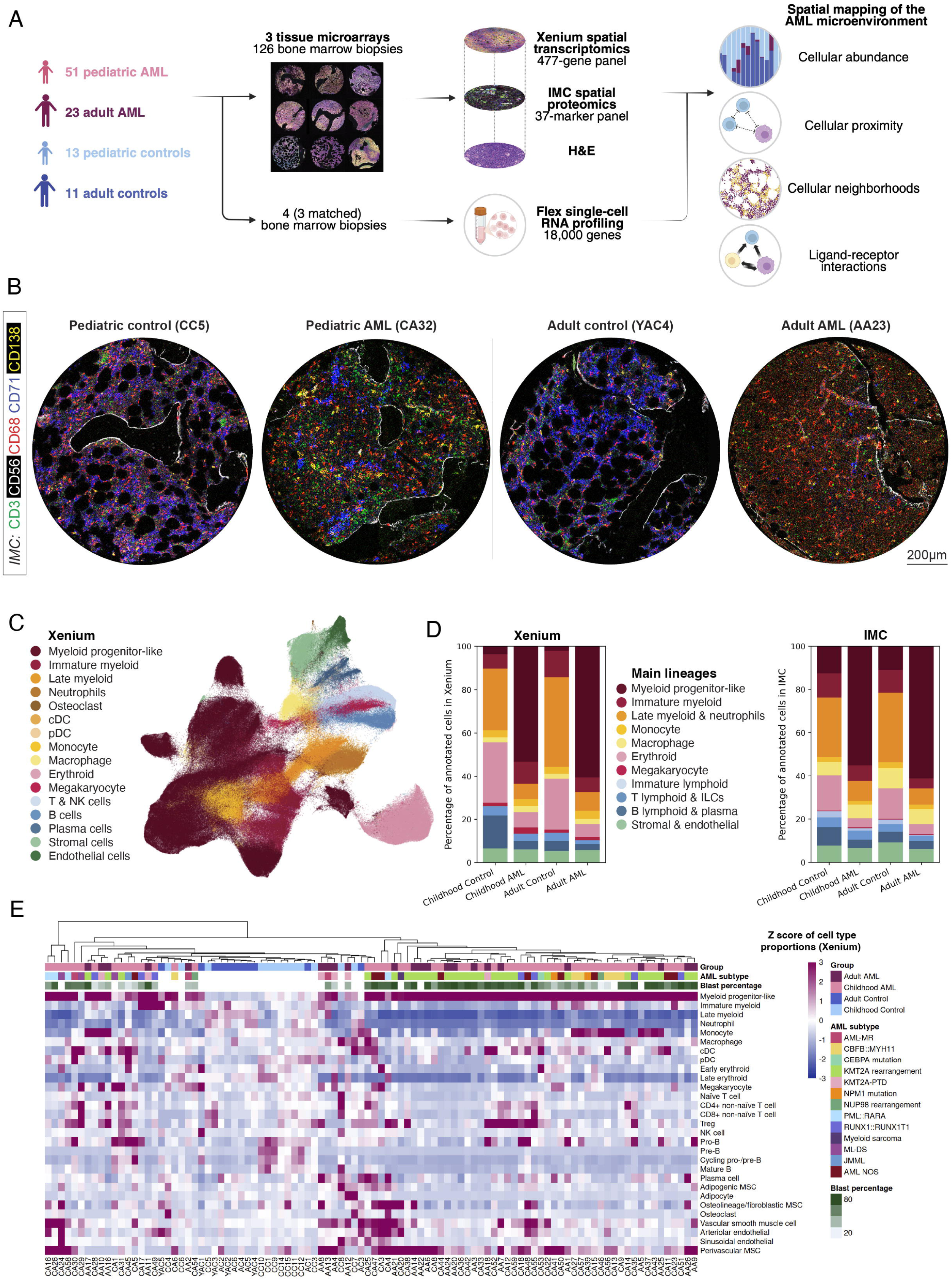
Spatial transcriptomics and proteomics identify all major hematopoietic and non-hematopoietic lineages in the AML bone marrow 1A. Study set-up showing the number of samples that were included after quality control. 1B. Representative IMC images of several key markers, for each of the age and condition groups. 1C. UMAP of annotated cell types in the Xenium data, showing the broad annotation of the main lineages (level 1 annotation); a UMAP with the fine annotation (level 2) can be found in Figure S1I. 1D. Comparison of proportions of main cell type lineages for Xenium and IMC. Immature lymphoid cells without clear B or T lineage bias were only identified as a separate category in the IMC data, because all immature B or T cells in the Xenium data were annotated as part of the B or T lymphoid cells. Late myeloid & neutrophils include late myeloid cells, neutrophils, conventional dendritic cells (cDC), plasmacytoid DCs (pDC), and osteoclasts. ILCs, innate lymphoid cells including NK cells. Statistical differences between the groups are presented in Supplemental tables S4-5. 1E. Proportions of cell types per individual (Xenium), presented as Z scores (compared to the mean of the controls) for the percentages of the detailed (level 2) cell type annotation (*i.e.* normalized per row).

All methods are described in detail in the supplements.

## Results

### Spatial transcriptomics and proteomics identify the major hematopoietic and non-hematopoietic lineages in the AML BM

To better understand the spatial organization and cellular interactions underlying immune evasion in AML, we generated a single-cell spatial transcriptomic and proteomic atlas of the pediatric and adult AML BM at diagnosis. We collected BM biopsies from 113 individuals, including pediatric (n=59, ages 0-18 years, 37% female) and adult (n=26, 20-79 years, 54% female) AML patients, alongside pediatric (n=14, 0-15 years, 43% female) and adult (n=14, 28-59 years, 43% female) non-leukemic controls without BM abnormalities (Figure S1A; Table S2). After quality control, 126 biopsies from 98 individuals were included for Xenium spatial transcriptomics (n=51 pediatric AML, n=23 adult AML, n=13 pediatric controls, n=11 adult controls; 675,235 cells) and 76 biopsies for IMC spatial proteomics (n=35 pediatric AML, n=20 adult AML, n=13 pediatric controls, n=8 adult controls; 634,012 cells) (Figures 1A, S1A; quality control reporting per sample in Supplemental table S3).

Both Xenium and IMC identified the major hematopoietic and non-hematopoietic lineages, with comparable cell frequencies between the modalities (Figures 1B-D, S1B). Immature myeloid clusters specific to AML or enriched for AML-associated markers were annotated as ‘myeloid progenitor-like cells’, while immature myeloid clusters detectable in both AML patients and controls were classified as ‘immature myeloid cells’ (Figure S1C-D). The AML BM was dominated by myeloid progenitor-like cells, which correlated positively with known blast percentages (Figure S1E-F); pediatric controls harbored more B cells; and adult controls more (mature) myeloid cells (Figure 1D, Supplemental tables S4-5)^25^. Notably, both Xenium and IMC identified macrophages (median 2.0% and 6.6%, respectively) and stromal cells (median 3.7% and 4.9%) at substantially higher frequencies than in typical BM aspirate-based analyses^26–28^, highlighting the advantage of biopsy-based spatial profiling.

Regarding detailed cell type annotation, IMC enabled distinction between major T lymphoid differentiation stages based on subtype-defining marker proteins (e.g., CD45RA, CD45RO, CD27) (S1G-H), while Xenium provided information on important T cell function markers and was able to identify several stromal and endothelial subtypes that could not be distinguished by IMC (Figures S1I-L). This showcases the complementarity of the two modalities in cell type identification. In addition, detailed cell type proportions were highly concordant between duplicate samples from the same individual (Figure S1M) and correlated with their morphological (FAB) classification (Figure S1N), emphasizing the robustness of the annotation.

Hierarchical clustering separated most patients from controls but showed no clear correlation with age (Figures 1E (Xenium), S1G-H (IMC)). Instead, it revealed substantial microenvironmental variation, ranging from blast-dominated samples with virtual absence of microenvironmental cells to samples enriched for T cells, often together with macrophages, and/or stromal cells.

In summary, this spatial atlas enables robust identification of the major hematopoietic lineages, with transcriptomics and proteomics providing both cross-validation and complementarity for cellular subtype identification.

### Macrophages show a predominantly immunosuppressive phenotype in both AML and non-leukemic BM

Immune profiling studies have shown a predominance of anti-inflammatory macrophages in the AML BM associated with poorer prognosis^12,20,29^. However, macrophage heterogeneity and interactions in the human AML BM have so far been incompletely characterized due to their poor detection in BM aspirate-based RNA-seq studies^12,23^. We therefore leveraged our spatial, BM biopsy-based dataset to deeply characterize AML-associated macrophages.

Spatial transcriptomics revealed that macrophage proportions did not differ significantly between AML or controls, age groups, or AML subtypes (Figures 2A, S2A-B). In both AML and controls, macrophages expressed both pro- and anti-inflammatory markers, with a tendency towards an anti-inflammatory phenotype. Virtually all macrophages identified by IMC expressed both the pan-macrophage marker CD68 and the anti-inflammatory marker CD163 (Figures 2B-C). On a transcriptomic level, anti-inflammatory markers such as *CD163*, *MRC1* and *CXCL12* were highly expressed, with lower expression of pro-inflammatory markers such as *CD80* and *CD86* (Figure 2D, S2C). Using unsupervised clustering, no clear pro-inflammatory or AML-specific macrophage population could be identified. Macrophages were generally distributed throughout the BM in both patients and controls, suggesting a potential for widespread interaction within the microenvironment (example in Figure 2B), which was subsequently quantified in further detail in the neighborhood analysis (Figure 3). In summary, macrophages in both AML and non-leukemic BM have a mixed phenotype, tending towards immunosuppression.

**Figure 2:**
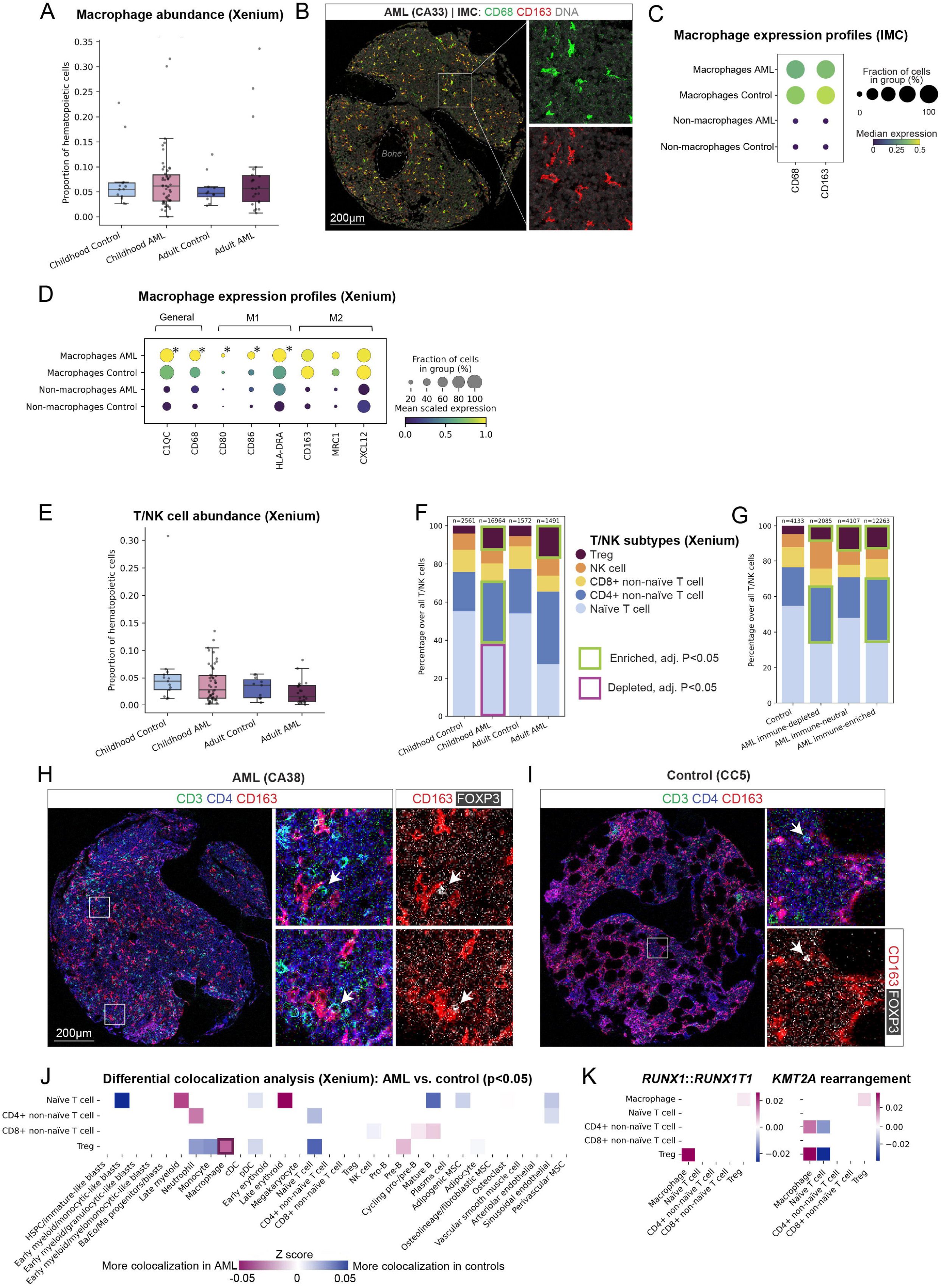
AML is characterized by immunosuppressive macrophages and an enrichment of regulatory T cells with increased macrophage colocalization. 2A. Macrophage abundance between age and condition groups (Xenium); no significant differences between the groups. 2B. Representative IMC image of patient CA33, showing the expression of the macrophage markers CD68 and CD163. 2C. Quantification of protein expression of macrophage markers CD68 and CD163 in macrophages and all other cells excluding myeloid progenitor-like cells (‘non-macrophages’) as a reference (IMC). 2D. Gene expression in macrophages and all other cells (‘non-macrophages’) as a reference, showing a selection of general macrophage markers (‘General’), in addition to markers associated with a pro-inflammatory (M1) or immunosuppressive (M2) macrophage phenotype (Xenium). Asterisks indicate those genes that are upregulated in macrophages in either AML or controls (adjusted P<0.05, log2 fold change >0.1). The gene expression of all macrophage-associated genes used for annotation is shown in Figure S2C. 2E. T cell abundance (level 1 annotation) as percentage of all hematopoietic cells (Xenium); no significant differences between the age and condition groups. 2F. T cell subtype abundance (level 2 annotation) over the T/NK cell compartment per age and condition group, showing Treg enrichment in AML. The boxes surrounding T/NK subtypes indicate significant (adjusted P value <0.05) enrichment or depletion compared to controls of the same age (Xenium). 2G. T cell subtype abundance (level 2 annotation) over the T/NK cell compartment for the different AML immune abundance categories. The boxes surrounding T/NK subtypes indicate significant (adjusted P value <0.05) enrichment or depletion compared to controls (Xenium). 2H. Representative IMC image of an AML sample showing colocalization of macrophages (CD163) and T cells (CD3, CD4, FOXP3). Arrows indicate FOXP3+ regulatory T cells. 2I. Representative IMC image of a control sample showing macrophages (CD163) and T cells (CD3, CD4, FOXP3) without clear colocalization. 2J. Heatmap showing differential colocalization analysis between AML and controls for the level 2 annotation; only significant differences (P<0.05) are shown. The dark purple box highlights the increased Treg-macrophage colocalization. 2K. Heatmap showing differential colocalization analysis as in Figure 2J, for *RUNX1*::*RUNX1T1* vs. controls and *KMT2A*r AML vs. controls; only significant differences (P<0.05) are shown.

**Figure 3:**
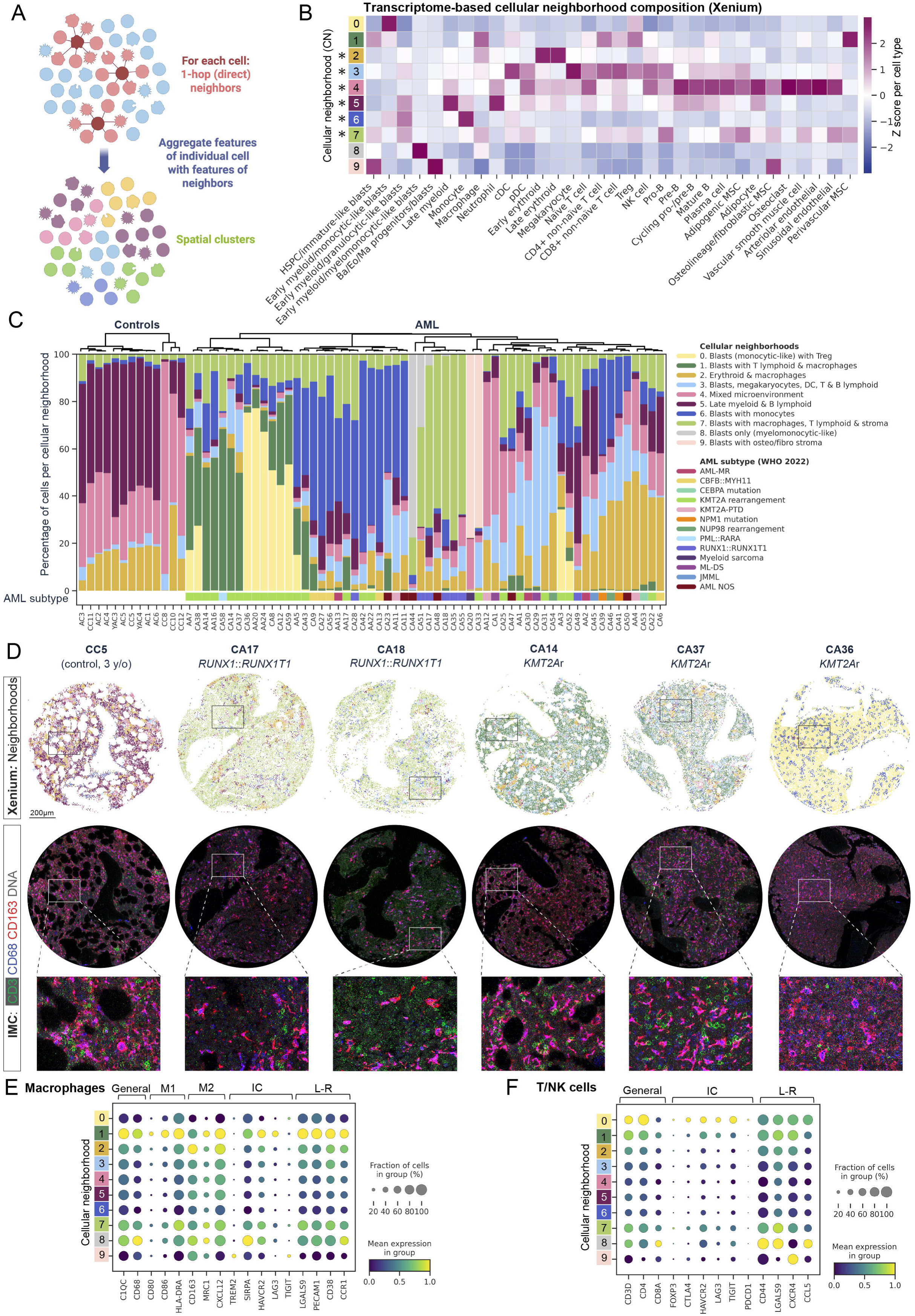
Cellular neighborhoods correlate with molecular subtypes and macrophage and T cell expression patterns. 3A. Schematic depiction of the method of neighborhood detection by CellCharter: for each cell, the transcriptomes of the 1-hop (direct) neighbors were considered their direct neighborhood; the transcriptomes of these neighbors were aggregated across all cells and all biopsies and clustered into spatial clusters (cellular neighborhoods). 3B. Cell type composition for the 10 cellular neighborhoods (CNs) identified by CellCharter (Xenium). Asterisks indicate the 6 CNs shared between patients and controls. 3C. Proportions of CellCharter CNs per individual in relation to condition (AML vs. control) and to AML subtypes, with hierarchical clustering based on similarity of CN proportions. 3D. Spatial plots showing examples of CN architecture in a control sample and 5 AML samples. 3E. Macrophage gene expression per CN. M1: pro-inflammatory macrophage markers. M2: anti-inflammatory macrophage markers. IC: immune checkpoint. LR: ligands and receptors. 3F. T/NK cell gene expression per CN.

### AML is characterized by enrichment of Tregs with increased macrophage colocalization

Besides macrophages, the lymphoid compartment plays an important role in the immunosuppressive AML BM microenvironment, with reports of an increased Treg frequency^17–19^ and T cell exhaustion. However, the underlying interactions remain insufficiently characterized. In our spatial data, overall T and NK cell proportions did not differ significantly between conditions and age groups (Figures 2D, S2D). Subclustering of the T/NK cells in the transcriptomics data identified naïve T cells, non-naïve CD4^+^ T cells, non-naïve CD8^+^ T cells, Tregs, and NK cells. In accordance with previous reports ^14,17–19,30^ and supporting an immunoregulatory microenvironment, both adult and pediatric AML showed significant enrichment of Tregs and a relative decrease in naïve T cells within the T/NK cell compartment compared to controls, suggesting an expansion of the memory compartment, which was also observed in our IMC data (Figures 2E, S2E-F).

Macrophages and T cells showed coordinated enrichment or depletion in the same individuals (Figure 1E). Indeed, T/NK cell and macrophage abundances were significantly correlated in AML (Spearman =0.54, P=7.7e-7), but not in controls (Spearman =0.37, P=0.074) (Figure S2G). Based on the abundance of T/NK cells and macrophages in controls, we defined three immune categories in AML: immune-neutral (combined macrophage and T/NK cell proportion within the interquartile range of controls), immune-depleted (below the first quartile), and immune-enriched (above the third quartile). These immune categories were related to myeloid progenitor-like cell abundance (Figure S2H), with lower percentages of myeloid progenitor-like cells in immune-enriched samples, but did not differ between adult and pediatric AML (Figure S2I), and the three categories were detected across the various AML subtypes (Figure S2J). Notably, Tregs were enriched across all immune categories compared to controls (Figure 2F), indicating that immunosuppression is a general feature of AML, independent of overall immune cell abundance.

Given the Treg enrichment in AML, we wondered whether Tregs also show spatial differences between AML and controls. In the IMC images (examples in Figures 2G-H), we observed colocalization of FoxP3+ T cells with macrophages in AML, while only few FoxP3+ T cells were identified in controls without a clear colocalization pattern. Comparative Xenium-based analysis revealed that Tregs indeed colocalized significantly more with macrophages in AML vs. controls (Figures 2I, S2K). This was consistent across various molecular AML subtypes (*KMT2A*r, *RUNX1*::*RUNX1T1*, *CBFB*::*MYH11*, and not otherwise specified AML (AML NOS)) (Figures 2J, S2L). Increased Treg-macrophage colocalization was also seen across all three immune categories (Figure S2M), suggesting that even in samples with low macrophage and T/NK cell abundance, these cells have a higher likelihood of interacting compared to the non-leukemic BM. Together, these findings point towards an immunoregulatory T cell compartment in AML, with potential immunosuppressive crosstalk between Tregs and macrophages.

### Cellular neighborhoods enriched for macrophages and T cells correlate with molecular AML subtypes and macrophage/T cell immune checkpoint expression

Cells in the BM typically function in a larger context of multiple cell types, so-called cellular neighborhoods (CN). To assess cell compositions, we performed CN analysis in two complementary ways: based on transcriptome and based on cell types annotated in the proteomics data. Using the transcriptome-based method CellCharter^31^, we identified 10 CNs: 6 shared between AML and controls, and 4 AML-specific CNs (Figures 3A, S3A-B). The three major shared CNs included erythroid (CN2), late myeloid and B lymphoid (CN5) and mixed microenvironmental/stromal CNs (CN4) (Figure 3B). In most controls, the late myeloid CN5 predominated, interspersed by erythroid islands (CN2) (example in Figure 3D). The AML-specific CNs were enriched for myeloid progenitor-like cells with varying involvement of immune or stromal cells, reflecting interpatient variability in microenvironmental abundance (Figure 1E). Two AML-specific CNs (CN1 and CN7) were enriched for AML with a varying extent of macrophages and T cells, consistent with the findings of the colocalization analysis.

Next, we asked whether CN composition was related to specific AML subtypes, and indeed observed several correlations (Figure 3C). CN7 (AML with T lymphoid, macrophages and stromal cells) was strongly enriched in cases with *RUNX1*::*RUNX1T1* AML, whereas CN0 (a mostely immune-depleted AML with some Tregs) and CN1 (AML with T cells and macrophages) were largely restricted to *KMT2A*r AML. The IMC-based CN analysis confirmed the presence of AML-macrophage-T cell CNs across most patients, and the correlation between KMT2Ar AML with an AML-only CN (Figures S3C-E).

While macrophage subtypes were not distinguishable by gene or protein expression alone, we hypothesized that their phenotype may be affected by their spatial colocalization, especially with T cells. Indeed, macrophages in CN1 and CN7 (colocalizing with AML and T cells) had increased gene expression of the immune checkpoints *HAVCR2* (encoding TIM-3) and *LAG3*, and the ‘don’t eat me’ receptor *SIRP* compared to most other CNs (Figure 3E). Similarly, T cells in these CNs and CN0 (mainly AML with some Tregs; Figure S3F), exhibited the highest expression of *CTLA4*, *HAVCR2*, *LAG3* and *TIGIT* (Figure 3F), which are associated with T cell inhibition or exhaustion^32^. These transcriptional differences in both macrophages and T cells may reflect immunosuppressive interactions in these CNs.

In summary, we identified distinct AML-associated CNs enriched for macrophages and T cells, that correlate with molecular subtypes and immune checkpoint expression, suggesting spatially organized, subtype-specific immunosuppressive interactions in the BM microenvironment.

### Ligand-receptor interaction analysis reveals immunosuppressive pathways involved in macrophage-T cell interactions

Given the spatial proximity between macrophages, myeloid progenitor-like cells and T cells, we used CellChat^33^ to predict ligand–receptor interactions driving cell-cell communication in the AML BM (Figure 4A). Looking at major signaling pathways between myeloid progenitor-like cells, macrophages, and T cells, cellular crosstalk was primarily mediated by Galectin-9 (LGALS9), MHC class I/II, PECAM-1, and chemokine axes (Figure 4A). Zooming into specific ligands and receptors, CellChat predicted Galectin-9 signaling from macrophages and myeloid progenitor-like cells to the inhibitory receptors HAVCR2 (TIM-3) on regulatory T cells (Tregs) and CD44 across T-cell subsets (selection in Figure 4B; full results in S4A). These findings were consistent across molecular subtypes (Figure S4B-E) and were orthogonally confirmed by 10x Flex single-cell RNA profiling on 4 FFPE AML BM biopsies (Figures S4F-G). Furthermore, CellChat revealed active recruitment and functional enhancement circuits, characterized by macrophage-derived CXCL12 acting on T cell CXCR4, alongside T cell-derived CCL5 targeting macrophage CCR1/CCR5 (Figure 4B).

**Figure 4:**
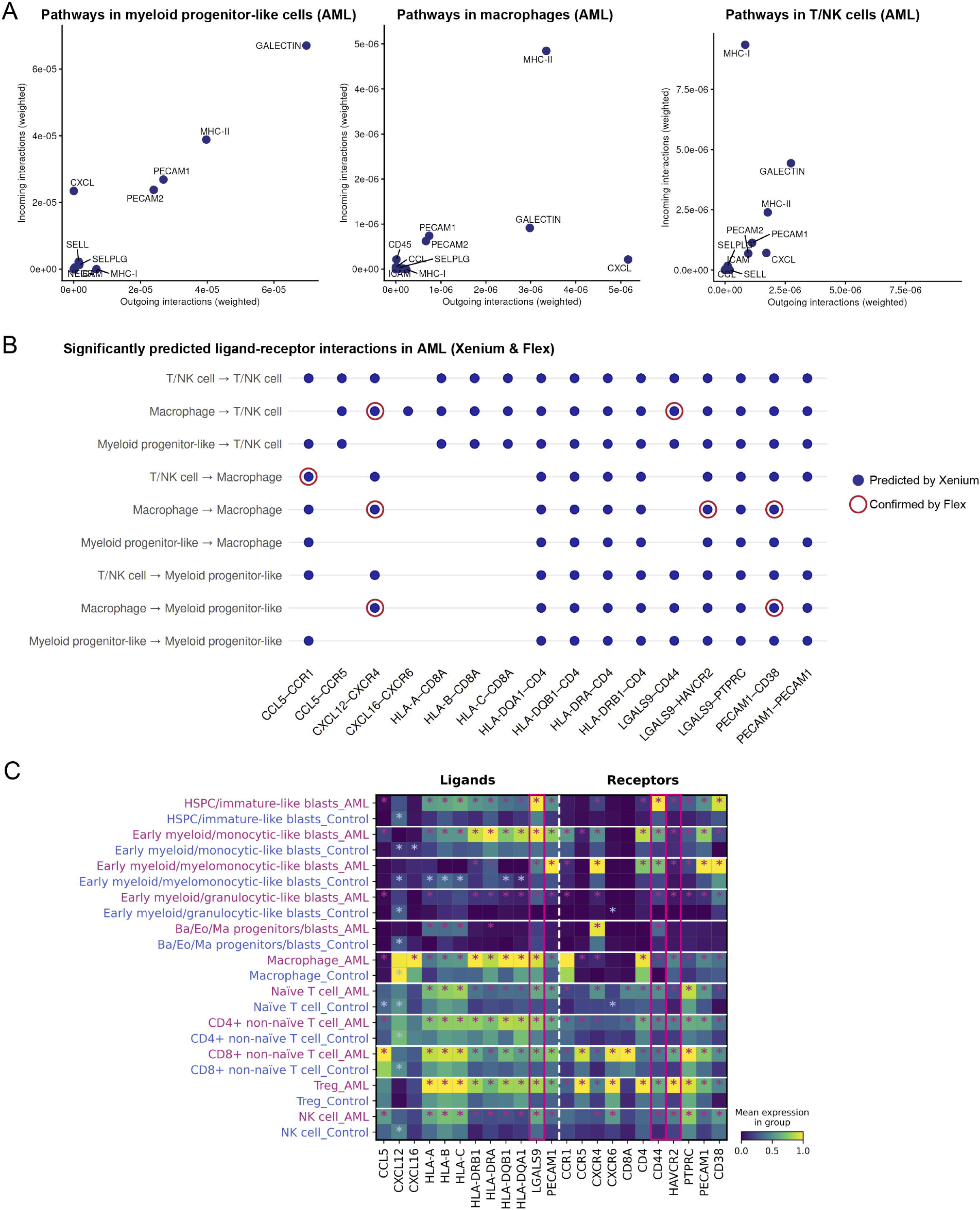
Ligand-receptor interaction analysis reveals immunosuppressive pathways involved in macrophage-T cell interactions. 4A. Scatterplots of the strength of outgoing and incoming signaling pathways for myeloid progenitor-like cells, macrophages, and T/NK cells, on AML patients only (spatially restricted CellChat analysis using level 1 annotation). 4B. Significant (P<0.05) predicted ligand-receptor interactions between T cells, macrophages, and myeloid progenitor-like cells based on the Xenium and Flex data, selection (see Figure S4A for the full results). 4C. Ligand and receptor gene expression (Xenium data) in macrophages, myeloid progenitor-like cell subtypes and T cell subtypes of the relevant ligand-receptor pairs identified by CellChat, for AML and controls separately. Ligands and receptors specifically mentioned in the text (*LGALS9*, *CD44*, *HAVCR2*) are highlighted with pink boxes. Asterisks indicate if a specific gene is expressed significantly higher (adjusted P<0.05) in AML (pink) or controls (blue) according to the differential gene expression analysis for that cell type.

Gene expression of these key ligand-receptor pairs was significantly upregulated in AML compared to non-leukemic BM (Figures 4C, S4H) and was elevated in immune-neutral and immune-enriched patients relative to immune-depleted samples (Figure S4I). Together, this suggests that enhanced signaling through these axes contributes to the immunosuppressive AML BM microenvironment. Indeed, Galectin-9–TIM-3 interactions have previously been shown to contribute to immune evasion in AML by impairing lymphoid cytotoxicity^34^, and to induce an anti-inflammatory phenotype in macrophages in other malignancies^35,36^. Moreover, Galectin-9– CD44 promotes generation of induced Tregs^37^, while CXCL12–CXCR4 and CCL5–CCR5 upregulation suggests enhanced Treg immunosuppressive function and recruitment, respectively^38–40^. Overall, these spatial and expression analyses predict and define an immunosuppressive network in the AML BM microenvironment that is anchored by Galectin-9 – CD44/TIM-3 signaling.

### Fusion-based identification of AML cells highlights enhanced interactions with their microenvironment compared to non-leukemic myeloid cells

Our ligand-receptor analyses between AML and the microenvironment relied on the annotated myeloid progenitor-like cells which may comprise both leukemic and non-leukemic cells. To directly identify malignant cells, we included a probe for the *RUNX1*::*RUNX1T1* fusion transcript in the Xenium panel (Figure S5A). The fusion was specifically detected in all 7 *RUNX1*::*RUNX1T1*-positive cases and correlated with their flow cytometry-based blast percentages (Figure 5A). Fusion-positive cells showed higher expression of proliferation markers (e.g., *MKI67*, *PCNA*) compared to their fusion-negative counterparts (Figure 5B), and were most abundant among HSPC/immature-like, granulocytic-like, and eosinophil progenitor-like blasts (Figure 5C), consistent with known differentiation patterns^27,41^. Together, these observations support the specificity of the fusion detection and the malignant identity of the fusion-positive cells.

**Figure 5:**
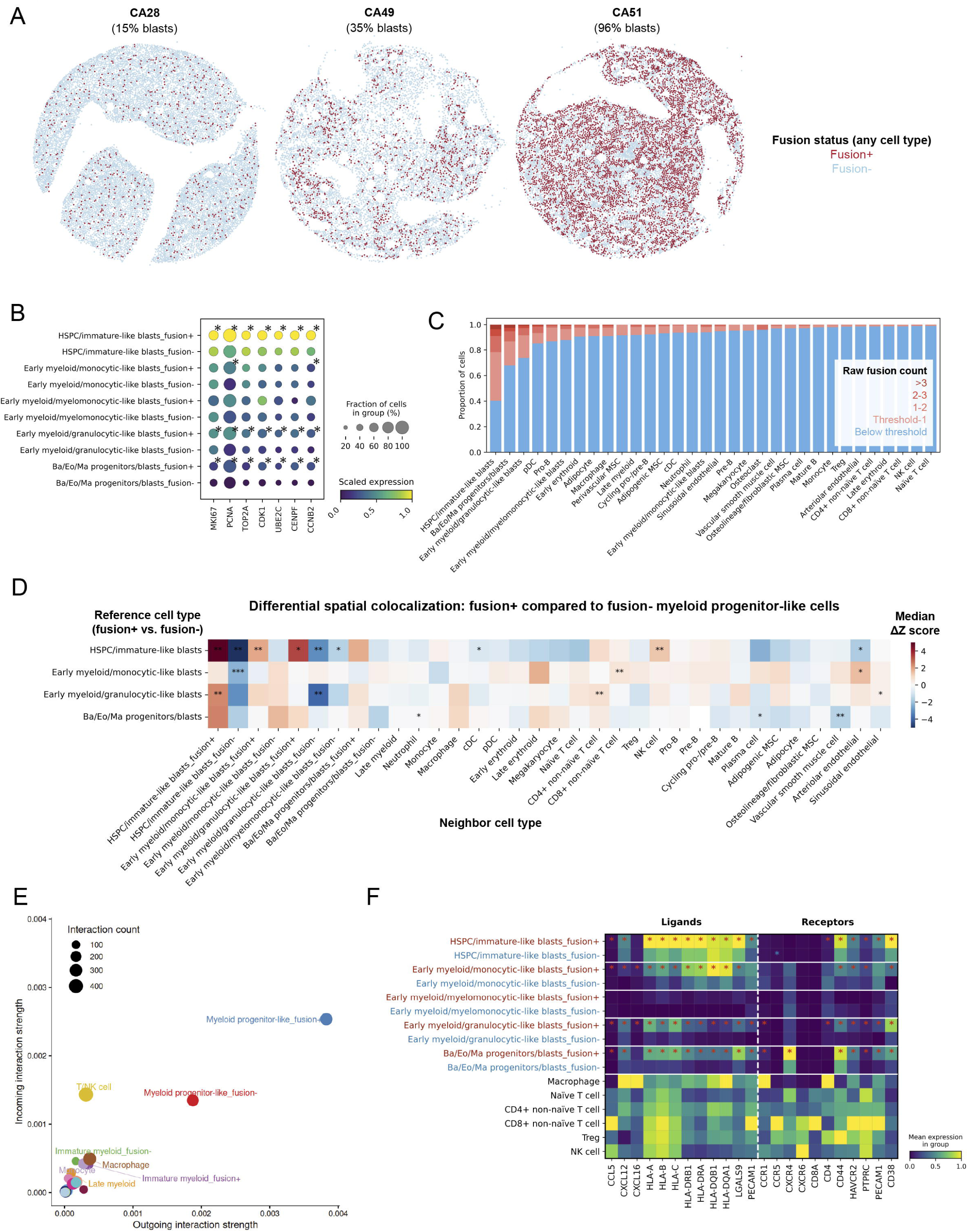
Fusion transcript detection aids identification of malignant cells and confirms ligand-receptor interactions. 5A. Spatial plot of fusion detection in 3 samples with varying reported blast percentages. 5B. Proliferation marker expression for the fusion-positive and fusion-negative versions of the myeloid progenitor-like cell types (for all 7 patients with *RUNX1*::*RUNX1T1* AML). Asterisks indicate if a specific gene is expressed significantly higher (adjusted P<0.05) in fusion-positive or fusion-negative cells, according to the differential gene expression analysis for that cell type. 5C. Bar chart displaying the level fusion detection per cell type, presented as raw counts (which are fractional because of the application of the probability-based segmentation algorithm Proseg). 5D. Differential colocalization of fusion-positive vs. fusion-negative cells: the difference in spatial colocalization between fusion-positive cells (on the y axis) and all other cell types (on the x axis) compared to the colocalization between the fusion-negative counterparts with all other cell types. Performed per sample using CellCharter neighborhood enrichment, presented as median delta Z score. Red means that the fusion-positive cells of the blast-like cell types (on the y axis) have stronger colocalization with the neighboring cell types (on the x axis) than their fusion-negative counterparts; blue means that fusion-positive cells have less colocalization compared to their fusion-negative counterparts. Wilcoxon rank sum test: * unadjusted P <0.05, ** unadjusted P<0.01. After Benjamini-Hochberg false discovery rate correction, none of the P values were significant, due to the small sample size (11 samples from 7 patients). 5E. Interaction strength as predicted by CellChat, per level 1 cell type (taking the population size as present in the data, not correcting for it). 5F. Expression of ligands and receptors of interest across fusion-positive and fusion-negative myeloid progenitor-like cells, T cell subtypes and macrophages, for all patients with *RUNX1*::*RUNX1T1* AML in this cohort together. Asterisks indicate if a specific gene is expressed significantly higher (adjusted P<0.05) in fusion-positive (red) or fusion-negative cells (blue) according to the differential gene expression analysis for that cell type.

To define leukemia-specific interactions of myeloid progenitors with macrophages and T cells, we compared colocalization patterns of fusion-positive (leukemia-associated) vs. fusion-negative (benign) cells of the same myeloid subtype. Although the sample size was too small to reach statistical significance after multiple testing correction, we observed a trend of preferential colocalization between fusion-positive cells with other fusion-positive cells (Figure 5D), especially for the most immature myeloid progenitors, suggesting spatial clustering of malignant cells and their descendants. Spatial proximity analysis further revealed a trend towards increased colocalization between fusion-positive myeloid progenitor-like cells and T/NK cell subsets compared to fusion-negative cells, consistent with the preferential blast-lymphocyte engagement observed in our neighborhood analyses.

Finally, we aimed to validate the ligand-receptor interactions predicted in the entire patient cohort by comparing fusion-positive vs. fusion-negative cells of the same annotated cell type in the subgroup of *RUNX1*::*RUNX1T1*-translocated AML. Overall, fusion-positive cells exhibited stronger predicted interactions with other cells compared to fusion-negative cells (Figure 5E), even after correction for cell number differences (Figure S5B). As reflected by their ligand-receptor expression, fusion-positive myeloid progenitor-like cells were predicted to have stronger interactions with other myeloid progenitor-like cells, macrophages and T cells via the LGALS9, MHC-II and PECAM1 pathways, confirming the importance of these pathways in the AML microenvironment (Figures 5F, S5C-F). In summary, *in situ* detection of the *RUNX1*::*RUNX1T1* fusion aided identification of malignant cells and revealed close spatial interactions between AML cells and the BM microenvironment.

### Macrophages induce differentiation of naïve CD4^+^ T cells into Tregs *in vitro*

The observed colocalization of macrophages and Tregs in AML suggested immunosuppressive crosstalk involving AML cells, macrophages, and T cells. We hypothesized that macrophages induce a Treg phenotype in naïve CD4^+^ T cells and/or attract Tregs from the periphery.

To assess whether macrophages promote Treg differentiation, naïve CD4^+^ T cells were co-cultured with unpolarized (M0) macrophages in the presence or absence of AML cells (Figure S6A). Direct contact between macrophages and naïve T cells, but not transwell conditions, induced robust differentiation into CD4⁺CD25⁺FOXP3⁺ cells (∼40% after 2 days, ∼80% after 4 days; ratio 1:1) (Figures 6A-B). This population is hereafter referred to as induced Tregs (iTregs), while acknowledging that despite their extensive characterization, stability of this phenotype cannot be assumed.

**Figure 6:**
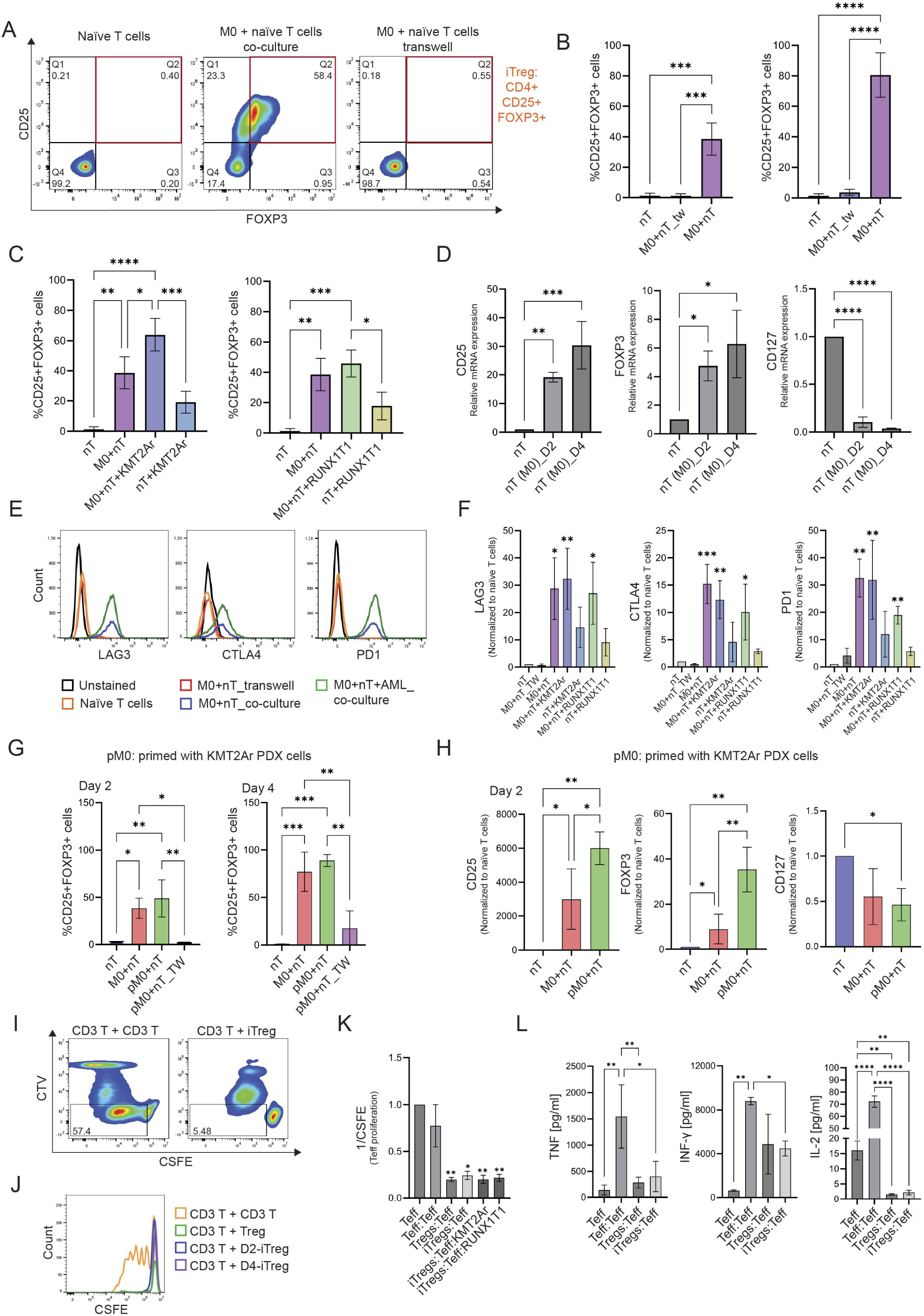
Induction of Tregs through macrophage-naïve T cell crosstalk. 6A. Representative flow cytometry plots showing the induction of induced regulatory T cells (iTregs; CD4^+^CD25^+^FOXP3^+^) from naïve CD4+ T cells (nT) cultured alone (monoculture), co-cultured with macrophages (M0), or cultured in a transwell setting. 6B. Percentage of iTregs after 2 and 4 days of co-culture with macrophages (M0+nT) or in transwell conditions (M0+nT_TW) at a 1:1 ratio of macrophages to naïve CD4^+^ T cells (n=3); 6C. Percentage of CD25^+^FOXP3^+^ cells after 2 days of co-culture with macrophages and KMT2A-rearranged (*KMT2A*r) patient-derived xenograft (PDX) cells (left) or *RUNX1*::*RUNX1T1* (*RUNX1T1*) PDX cells (right) at a 1:1 ratio (n=3). 6D. Relative RNA expression levels of CD25, FOXP3 and CD127 in naïve CD4+ T cells after 2 and 4 days of co-culture with macrophages (n=3). 6E. Representative flow cytometry plots showing immune checkpoints expression on naïve CD4^+^ T cells after 2 days in monoculture, or co-culture with macrophages (M0+nT_co-culture) and/or PDX cells (M0+nT+AML_co-culture), or in a transwell setting (M0+nT_TW). 6F. Quantification of immune checkpoint protein expression intensity on naïve CD4^+^ T cells after 2 days in monoculture, or in co-culture with macrophages alone, or with PDX cells, or in a transwell setting at a 1:1 ratio (n=3). 6G. Percentage of CD25^+^FOXP3^+^ cells after 2 and 4 days of co-culture with pM0 macrophages (macrophages pre-incubated with KMT2Ar PDX cells) at a 1:1 ratio (n=3). 6H. Quantification of CD25, FOXP3, and CD127 expression on iTregs generated from co-cultures between naïve T cells and macrophages primed with *KMT2A*r PDX cells (pM0) compared to co-cultures with (not primed) M0 (n=3). 6I. Representative flow cytometry plots of suppression assay between two populations of CD3^+^ T cells, between healthy CD25^+^ Treg and CD3^+^ T cells, or between macrophage-induced iTregs and CD3^+^ T cells. 6J. Histogram showing proliferation (CSFE signal) of CD3^+^ T cells cultured alone, co-cultured with another CD3^+^ T cell population, co-cultured with healthy donor CD25^+^ Tregs, or co-cultured with macrophage-induced iTregs. 6K. CSFE mean fluorescence intensity of CD3^+^ T cells after 5 days co-culture with healthy CD3^+^ T cells, CD25^+^ Tregs, or macrophage-induced D2-iTregs (n=3). 6L. Cytokine expression measured in supernatants from suppression assay. Statistical analysis: one-way ANOVA. Significance is indicated as *P < 0.05; **P < 0.01; ***P < 0.001; ****P < 0.001.

We next evaluated whether leukemic blasts further augment this macrophage-driven induction. While co-culture with KMT2Ar or *RUNX1*::*RUNX1T1* AML cells alone yielded modest iTreg differentiation (∼20% at day 2), adding KMT2Ar cells to the macrophage–T cell co-culture significantly enhanced iTreg frequencies by day 2 (increasing from ∼40% to ∼65%; Figures 6C, S6B–C). In contrast, adding *RUNX1*::*RUNX1T1* cells provided no additive enhancement (Figures 6C, S6B–C), potentially reflecting the monocytic lineage tendency and distinct immunomodulatory profile of KMT2Ar blasts^42^. By day 4, iTreg frequencies plateaued (Figure S6D). iTregs displayed marked upregulation of *CD25* and *FOXP3* compared to naïve CD4⁺ T cells, together with downregulation of CD127, consistent with a Treg phenotype^43,44^ (Figures 6D, S6E). No induction of these markers was observed when macrophages were co-cultured with healthy B cells, confirming T cell specificity (Figure S6F). Furthermore, iTregs exhibited significantly increased expression of the immune checkpoints CTLA4, PD-1 and LAG3 compared to naïve CD4⁺ T cells (Figures 6E-F, S6G), independent of AML presence or co-culture ratios.

Next, to recreate the AML immunosuppressive microenvironment, macrophages were pre-cultured with AML, before adding naïve CD4^+^ T cells. Pre-conditioning macrophages (pM0) did not alter iTreg frequency compared to naïve macrophages (Figures 6G, S6H). However, pM0-induced iTregs did express higher levels of CD25 and FOXP3 after 2 days (Figures 6H, S6I), suggesting additive effects of AML and macrophages on the Treg phenotype. Functionally, iTregs effectively suppressed CD3⁺ T cell proliferation similarly to CD25^high^ Tregs isolated from healthy donors (Figures 6I–K) and exhibited reduced expression of pro-inflammatory cytokines involved in T cell activation and proliferation, including TNF, IFN-γ and IL2 (Figure 6L).

Together, these data identify macrophages as potent inducers of suppressive FOXP3^+^ Treg-like differentiation *in vitro*, supporting their role in shaping an immunosuppressive environment in AML.

#### Bidirectional macrophage-iTreg interaction is driven by CD44 and TIM-3

Previous studies have shown that Tregs promote the differentiation of monocytes into alternatively activated, anti-inflammatory macrophages, establishing an immunoregulatory feedforward loop^45–47^. In our system, naïve (M0) macrophages co-cultured with naïve CD4^+^ T cells acquired a mixed activation phenotype during Treg induction, expressing both pro-inflammatory (HLA-DR, CD80) and anti-inflammatory markers (CD206, CD163, ARG2, IL-10) (Figures 7A-B), along with increased secretion of CXCL-10, IL-10, TNF, and IFN-γ, but not of IL-1β and IL-6 (Figure S7A). This hybrid phenotype showcases macrophage plasticity^48^ and resembles the phenotype observed in our spatial data (Figure 2D), indicating bidirectional macrophage-iTreg interaction.

**Figure 7:**
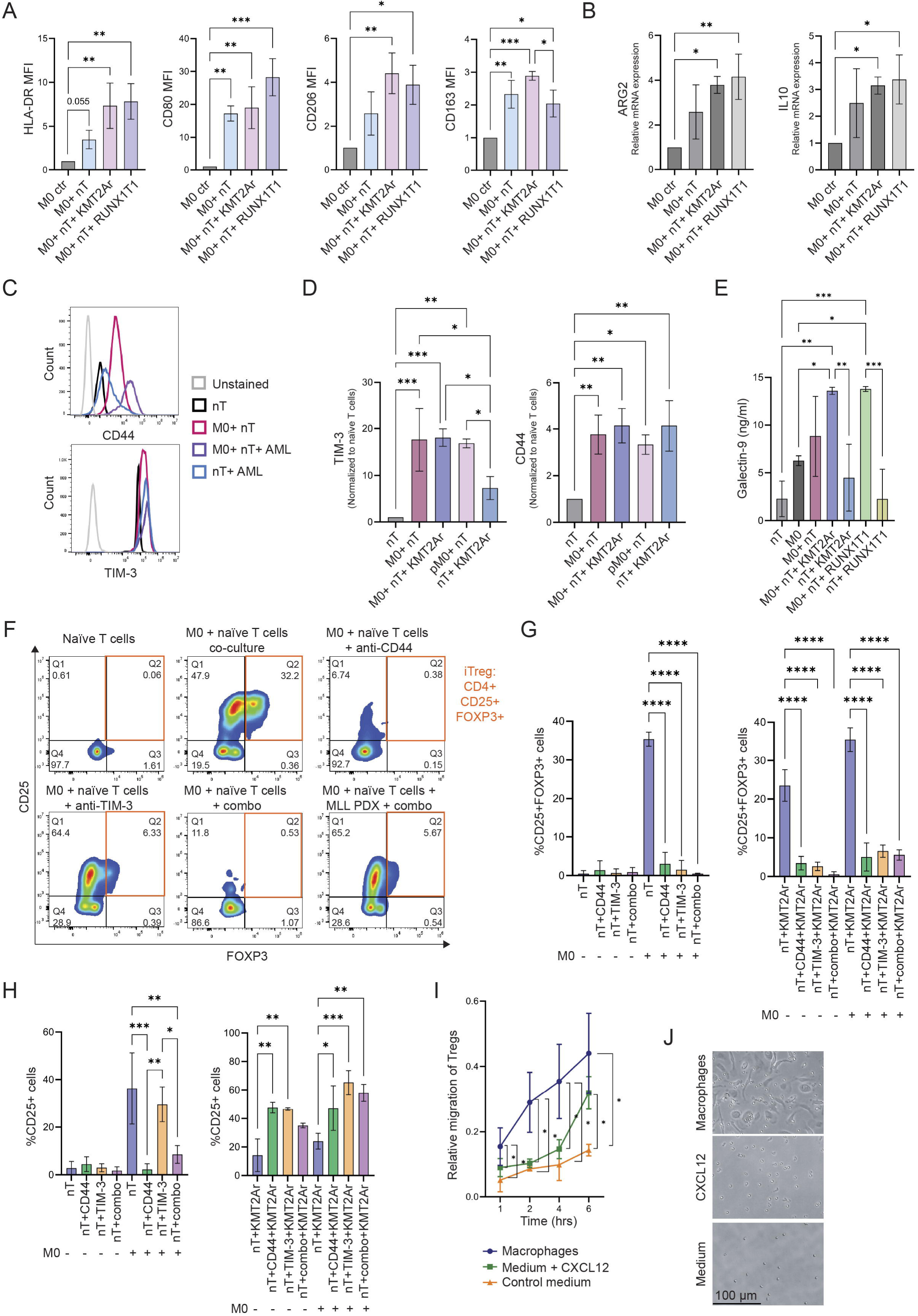
Bidirectional crosstalk between macrophages and iTregs. 7A. Quantification of HLA-DR, CD80, CD206, and CD163 protein expression on macrophages before and after 2 days co-culture with naïve CD4⁺ T cells, in the presence or absence of *KMT2A*r or *RUNX1T1* PDX cells (n=3). 7B. Relative mRNA expression of ARG2 and IL10 in macrophages before and after co-culture with naïve CD4⁺ T cells, in the presence or absence of PDX cells (n=3). 7C. Representative flow cytometry histograms showing TIM-3 and CD44 expression on naïve CD4⁺ T cells and iTregs under the indicated conditions. 7D. Quantification of TIM-3 and CD44 expression on naïve CD4⁺ T cells cultured alone or after 2 days of co-culture with macrophages at a 1:1 ratio, in the presence or absence of *KMT2A*r PDX cells (n=3). 7E. ELISA assay measuring Galectin-9 cytokine level in culture supernatants under the indicated conditions. 7F. Representative flow cytometry plots showing iTreg induction in naïve CD4⁺ T cells co-cultured with macrophages in the presence of anti-CD44 and/or anti-TIM-3 blocking antibodies. 7G. Percentage of CD4⁺CD25⁺FOXP3⁺ iTregs after 2 days of monoculture or co-culture with macrophages at a 1:1 ratio, in the presence of 10 µg/mL anti-CD44 and/or 10 µg/mL anti-TIM-3 blocking antibodies (n = 3). 7H. Percentage of CD4⁺CD25⁺ cells after 2 days of co-culture with KMT2Ar PDX cells and/or macrophages at a 1:1 ratio, in the presence of 10 µg/mL anti-CD44 and/or 10 µg/mL anti-TIM-3 blocking antibodies (n = 3). 7I. Transwell migration assay assessing CD25+ Tregs migratory capacity under the following conditions: macrophages, medium + CXCL12 and culture medium alone. 7J. Representative images from migration assays after 6 hours of incubation. Statistical analysis: one-way ANOVA. Significance is indicated as *P < 0.05; **P < 0.01; ***P < 0.001; ****P < 0.001.

To explore underlying mechanisms, we examined several predicted ligand-receptor pairs (Figures 7C, S7B). TIM-3 was strongly upregulated on iTregs following co-culture with macrophages, whereas CD44 was strongly induced by both macrophages and AML cells (Figures 7C-D, S7C-E). Both receptors bind Galectin-9, which was highly expressed by macrophages. Although macrophage Galectin-9 expression did not differ significantly between co-culture conditions (Figures S7F-G), the level of soluble Galectin-9 in the triple co-culture (macrophages, naïve T cells and AML) was higher than in macrophage or naïve T cell monocultures or in T cell-AML co-cultures, indicating that interaction among the three cell types promotes Galectin-9 secretion (Figures 7E, S7H). Blocking CD44 and/or TIM-3 significantly reduced the induction of CD4^+^CD25^+^FOXP3^+^ iTregs by macrophages, both in the presence or absence of AML (Figures 7F-G, S7I-J). However, TIM-3 blockade alone or blockade of TIM-3 and CD44 together in the presence of AML did not prevent CD25 induction. Since CD25 expression is also a marker of T cell activation, this may indicate that T cell activation is taking place without Treg differentiation^44^ (Figures 7H, S7K).

Finally, in addition to differentiation, we assessed whether macrophages can actively recruit Tregs using transwell migration assays. Indeed, macrophages attracted Tregs more efficiently than CXCL12 or medium alone within 4 hours, while after 6 hours, both macrophages and CXCL12 showed comparable chemotactic activity (Figures 7I-J).

Together, these findings support a bidirectional interaction in which macrophages promote iTreg differentiation and recruitment, and iTregs in turn induce a mixed activation in macrophages, reinforcing an immunosuppressive feedforward loop in AML.

## Discussion

The immunosuppressive BM microenvironment is regarded as an important obstacle to immunotherapy in AML^11^. However, its comprehensive characterization by traditional technologies and scRNA-seq has been hampered by the inability to capture all relevant cell types and the lack of spatial context. Previous spatial profiling studies have demonstrated the feasibility of spatial proteomics and transcriptomics in BM biopsies^25,49–54^, and provided the first important insights into AML spatial niches and immune signaling pathways involved in AML pathogenesis and response to immunotherapy^55–58^. However, these studies were restricted to small adult cohorts or murine BM, and due to technical constraints of the available platforms often lacked the spatial resolution or relevant panels for comprehensive cellular interaction analysis between AML and its microenvironment at the single-cell level^55–60^.

Therefore, we generated a single-cell spatial transcriptomic and proteomic atlas of the BM microenvironment in a large cohort of adult and pediatric AML across various subtypes. This approach enabled detection of typically underrepresented cell types, detailed mapping of the spatial architecture of AML, and identification of key cellular interactions. Notably, we observed colocalization between macrophages and Tregs, and demonstrated that macrophages and AML promote Treg differentiation via CD44 and TIM-3.

Unlike previous BM aspirate-based single-cell profiling studies, the rich representation of macrophages in our atlas enabled in-depth characterization of their localization and interactions. The combination of transcriptomics and proteomics strengthened our findings through cross-validation but also expanded them by leveraging the strengths of both technologies.

Besides the improved cell type detection, the spatial information enabled identification of CNs shared between AML and non-leukemic BM, as well as AML-specific neighborhoods. Remarkably, CN compositions correlated with specific AML subtypes, suggesting that subtype-specific cellular interactions may underlie differences in treatment response and prognosis between AML subtypes.

Furthermore, we identified crosstalk involving macrophages, T cells, and leukemic cells that contributes to the immunosuppressive AML BM microenvironment. While a predominance of anti-inflammatory macrophages and increased Tregs in AML have been described before^61^, we demonstrate that these populations also colocalize more in AML. We show that this may be caused by macrophage- and AML-induced differentiation of naïve T cells to FOXP3^+^ suppressive Treg-like cells. Our spatially informed ligand-receptor analysis predicted several pathways potentially involved in this interaction, including Galectin-9 with CD44 and/or TIM-3. A particular strength of our study is the detection of the *RUNX1*::*RUNX1T1* transcript in the subset of patients with this fusion, which aided identification of malignant cells and thereby supported the predicted ligand-receptor interactions. We subsequently confirmed the role of CD44 and TIM-3 in Treg-like induction by macrophages and AML *in vitro*.

Our findings extend previous observations that anti-inflammatory macrophages can induce Tregs from naïve T cells^62,63^ by demonstrating that AML-primed macrophages have similar capabilities. Moreover, iTregs may reciprocally influence macrophages by promoting a mixed activation phenotype, providing insight into the complex crosstalk underlying immunosuppression in AML. While naïve T cell conversion is one mechanism, some human evidence suggests that Tregs may also arise from (non-naïve) activated CD4^+^ T cells^64^, raising the possibility of multiple pathways contributing to Treg enrichment in AML.

These findings provide a foundation for new therapeutic strategies aimed at modulating the BM microenvironment. Targeting macrophage-T cell interactions, particularly the Galectin-9– CD44/TIM-3 axis, may enhance immunotherapy efficacy. For example, antibodies or small molecules targeting CD44 or TIM-3 could be combined with CAR-T or T cell engager therapy to increase their therapeutic efficacy by rendering the BM microenvironment less immunosuppressive. While CD44-targeted therapies have shown effects on AML cells^65–67^, our results suggest an additional immunomodulatory role to enhance other AML-directed immunotherapies. Interestingly, Galectin-9-targeting antibodies were recently found to reduce AML burden as well as immune suppression in the lung microenvironment of pulmonary extramedullary AML^68^, suggesting that targeting the Galectin-9 axis may be relevant for both medullary and extramedullary AML.

Another approach could involve engineering CAR-T cells to overcome exhaustion and improve persistence, such as by disrupting TIM-3 or CD44 signaling via secreted decoy receptors or deletion of inhibitory receptors ^69–71^. Spatial profiling of BM biopsies during treatment and in a low measurable residual disease setting may further refine the timing and application of such strategies.

Current spatial technologies require trade-offs between spatial resolution and the number of detectable targets^72^. Consequently, the interaction analysis in our study was restricted by the 477-gene panel. Additionally, transcript assignment to individual cells may be imperfect due to technical constraints^73–75^, and because of our probe-based approach we could not fully distinguish malignant from non-malignant myeloid progenitor-like cells, except for patients with a *RUNX1*::*RUNX1T1* translocation for which we included a fusion probe. Finally, the *in vitro* experiments inherently simplify the complexity of the AML BM microenvironment and may not fully recapitulate *in vivo* Treg differentiation pathways.

Taken together, our spatial transcriptomic and proteomic atlas of the AML BM provides detailed insight into the AML microenvironment across molecular subtypes and ages, identifying potentially targetable immunosuppressive interactions. Interfering with macrophage-T cell interactions via CD44 or TIM-3 might represent a promising strategy to enhance immunotherapy in AML.

## Resource availability

All generated data and the corresponding analysis scripts have been deposited in Zenodo at https://doi.org/10.5281/zenodo.19124967.

The publicly available data used in the study are from Bandyopadhyay et al.^57^, Zeng et al.^27^, and Hanemaaijer et al.^25^

## Acknowledgements

We would like to thank the following people for their contributions to this work:

- Rubina Moeniralam for sectioning the tissue.
- Domenico Castigliego for the preparation of the TMA.
- Jeff DeMartino for advice on the Xenium and Flex data analysis.
- Lindy Visser for her advice about CellChat and Flex data analysis.
- Enrico Muzzin for the artwork of the graphical abstract.
- Aleksandra Balwierz for her help with the preparation of the Flex experiment.

## Author contributions

Conceptualization: J.B.K., O.H., L.P., M.v.d.M., B.F.G.

Formal analysis: M.v.d.M., E.P., A.P., E.A., E.K.S., W.J.J.

Funding acquisition: O.H., C.M.Z., H.V., T.M.

Investigation: M.v.d.M., E.P., A.P., W.J.J.

Methodology: M.v.d.M., E.P., J.B.K., L.T.C., A.P., T.M., W.J.J.

Project administration: O.H., C.M.Z., W.J.J.

Resources: L.P., A.L., J.I., M.A.V., M.G., J.M.L.T., T.M.

Software: M.v.d.M., E.P., L.T.C.

Supervision: O.H., C.M.Z., B.F.G., M.G., J.M.L.T., H.V., M.B., D.H., E.v.d.A, S.N., J.V., T.M.

Validation: M.v.d.M., L.T.C., A.P.

Visualization: M.v.d.M., E.P., A.P.

Writing – original draft: M.v.d.M.

Writing – review & editing: all authors.

