## Supplemental figures, legends, methods and tables for "Spatial multi-omics reveals targetable immunosuppressive macrophage T-cell interactions in human AML bone marrow": Spatial AML_Supplemental figures_bioRxiv.pdf

### Figure S1

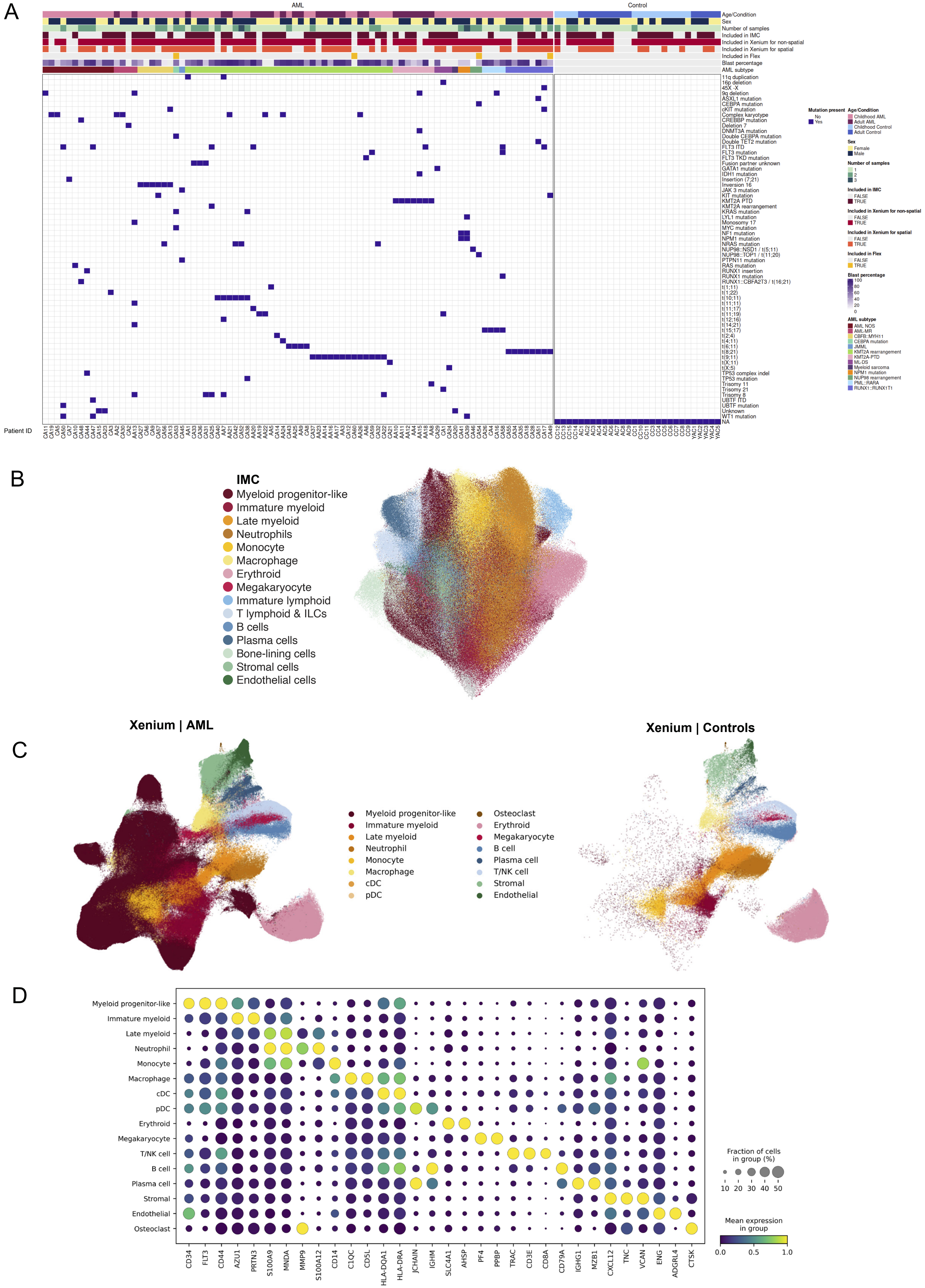

A
AML
Control
Age/Condition
Sex
Number of samples
Included in IMC
Included in Xenium for non-spatial
Included in Xenium for spatial
Included in Flex
Blast percentage
AML subtype
Patient ID
B
IMC
Myeloid progenitor-like
Immature myeloid
Late myeloid
Neutrophils
Monocyte
Macrophage
Erythroid
Megakaryocyte
Immature lymphoid
T lymphoid & ILCs
B cells
Plasma cells
Bone-lining cells
Stromal cells
Endothelial cells
C
Xenium | AML
Xenium | Controls
Myeloid progenitor-like
Immature myeloid
Late myeloid
Neutrophil
Monocyte
Macrophage
cDC
pDC
Osteoclast
Erythroid
Megakaryocyte
B cell
Plasma cell
T/NK cell
Stromal
Endothelial
D
Myeloid progenitor-like
Immature myeloid
Late myeloid
Neutrophil
Monocyte
Macrophage
cDC
pDC
Erythroid
Megakaryocyte
T/NK cell
B cell
Plasma cell
Stromal
Endothelial
Osteoclast
CD34
FLT3
CD44
AZU1
PRTN3
S100A9
MNDA
MMP9
S100A12
CD14
C1QC
CD5L
HLA-DQA1
HLA-DRA
JCHAIN
IGHM
SLC4A1
AHSP
PF4
PPBP
TRAC
CD3E
CD8A
CD79A
IGHG1
MZB1
CXCL12
TNC
VCAN
ENG
ADGRL4
CTSK
Fraction of cells in group (%)
10 20 30 40 50
Mean expression in group
0.0 0.5 1.0

Figure S1 (continued)

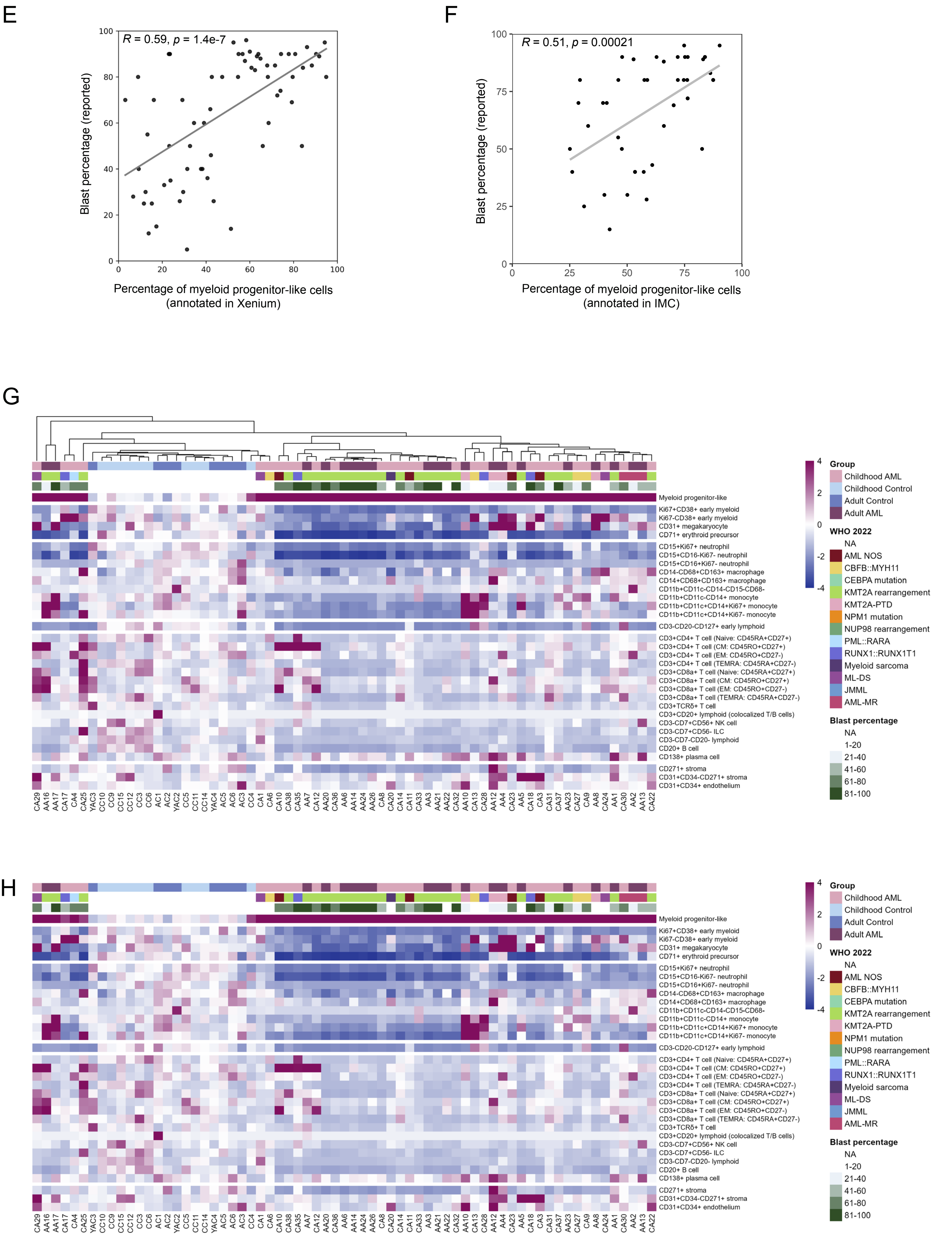

Figure S1 (continued)

I

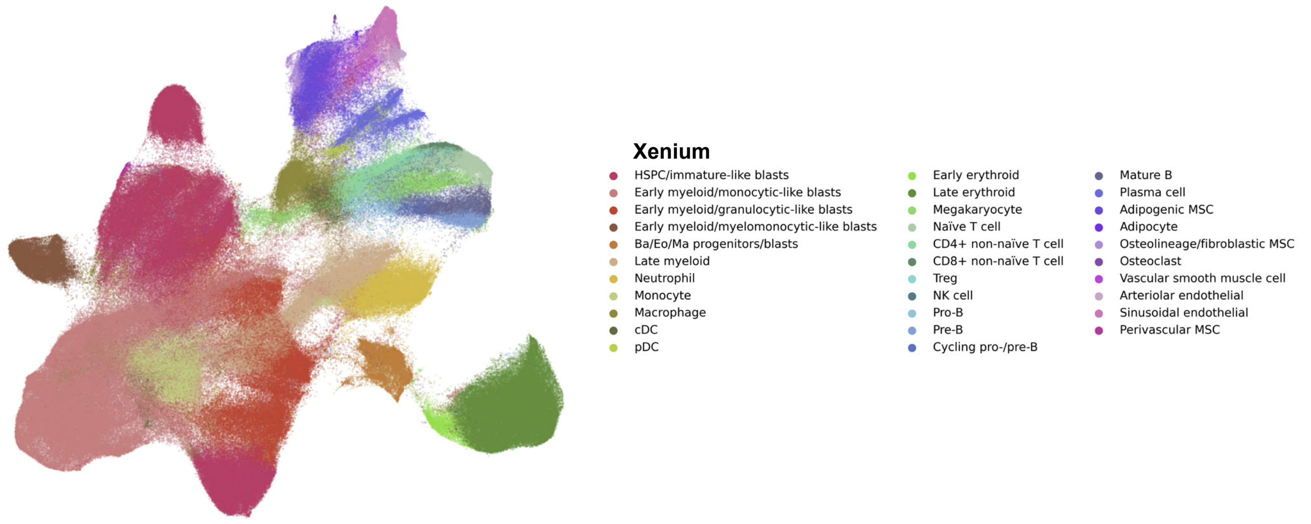


J

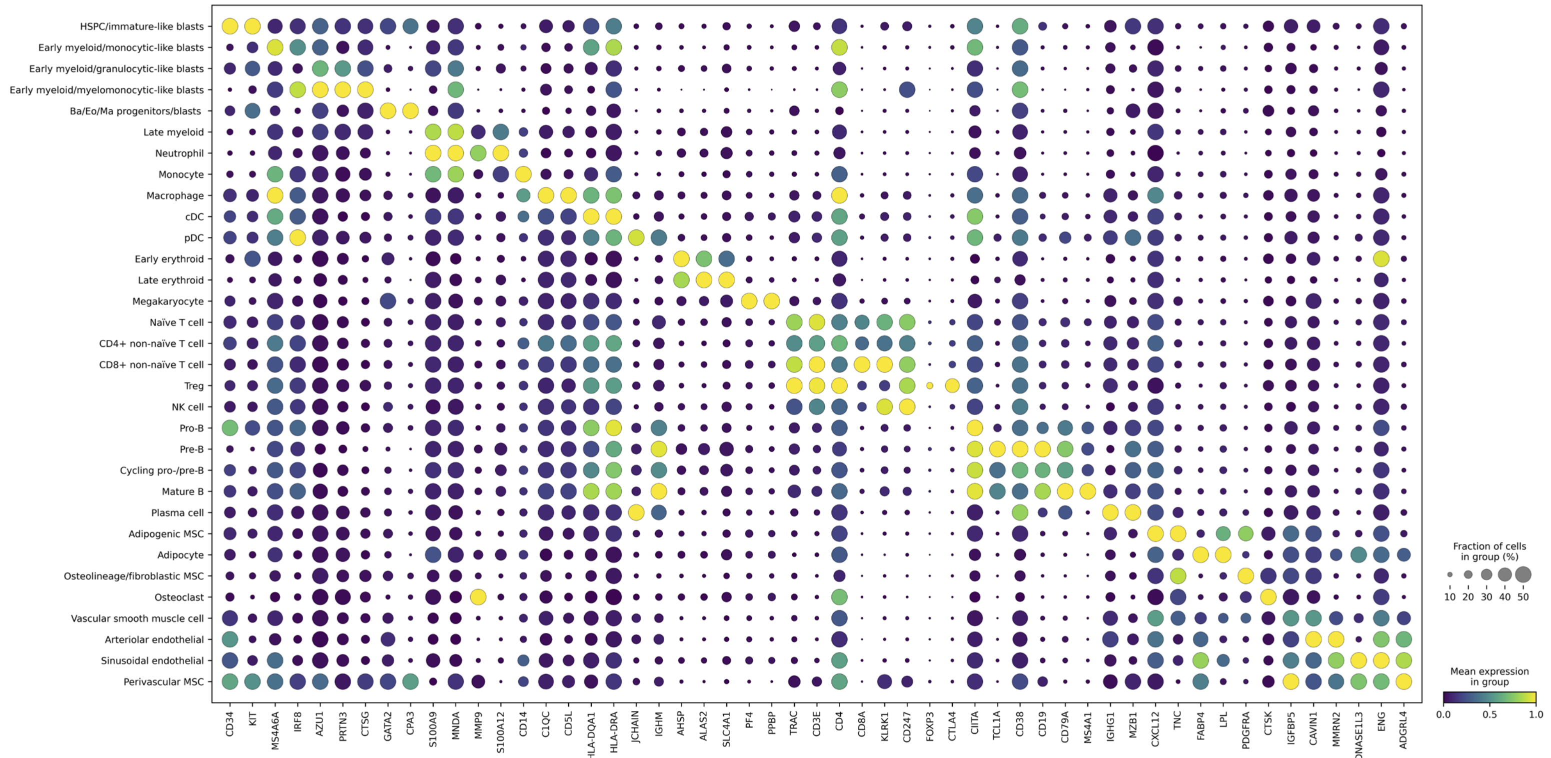


K

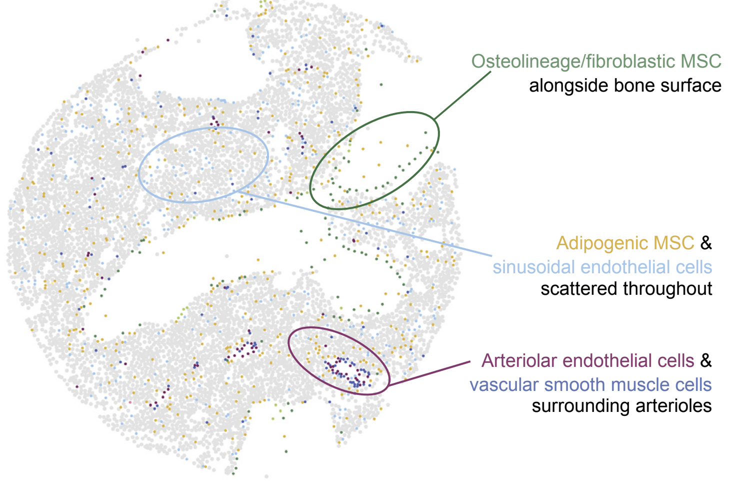


L

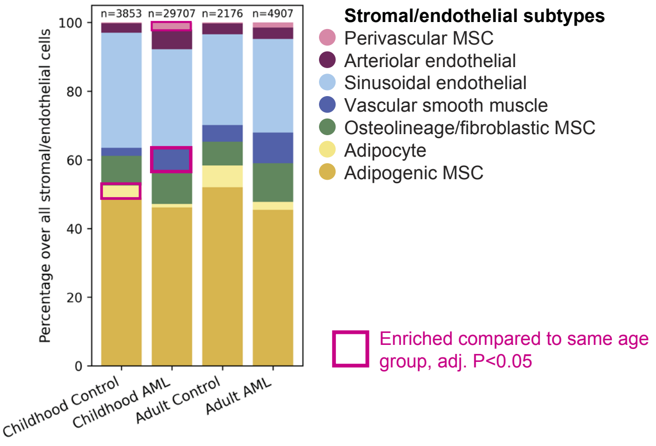

### Figure S1 (continued)

M

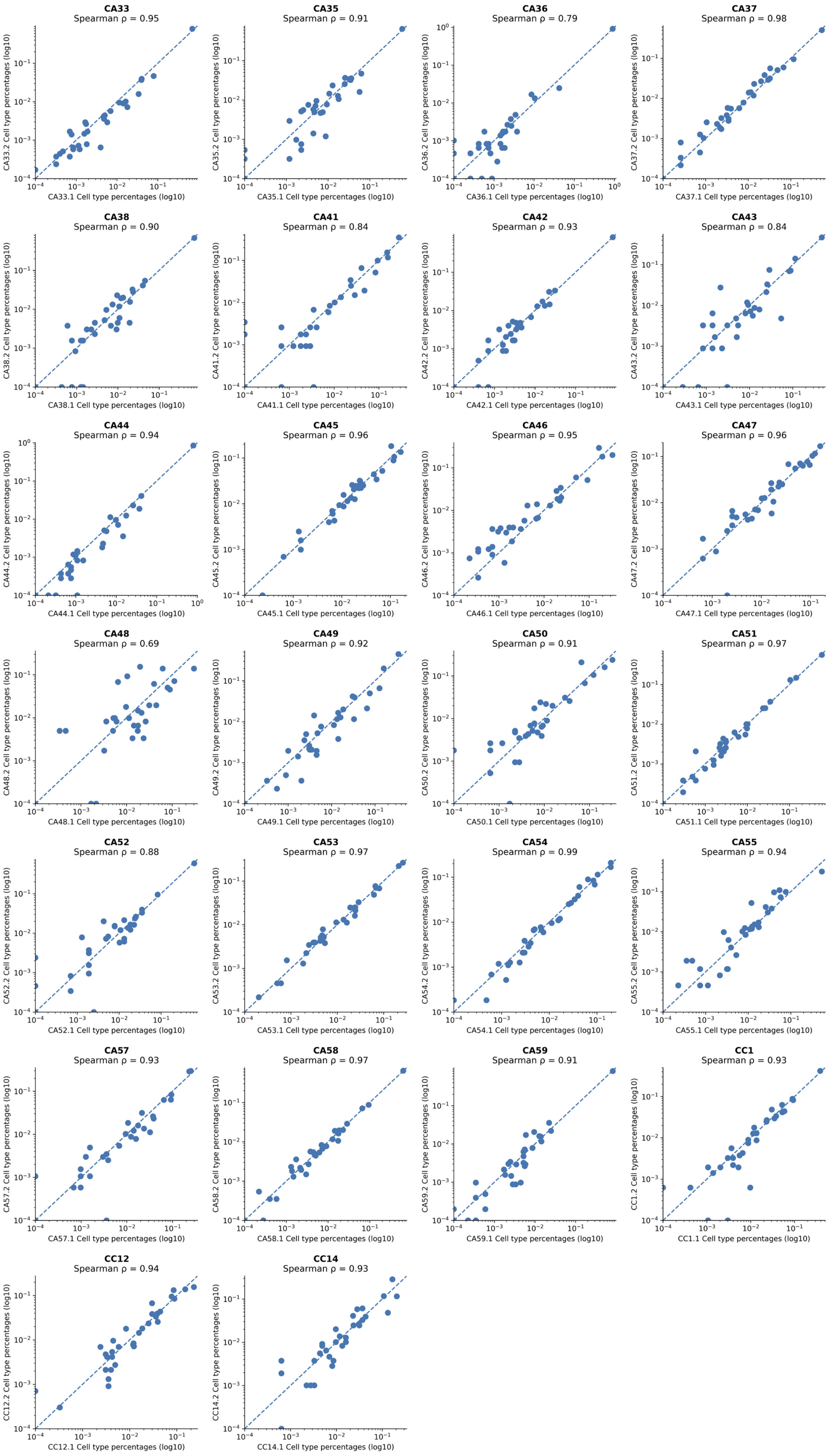

### Figure S1 (continued)

N

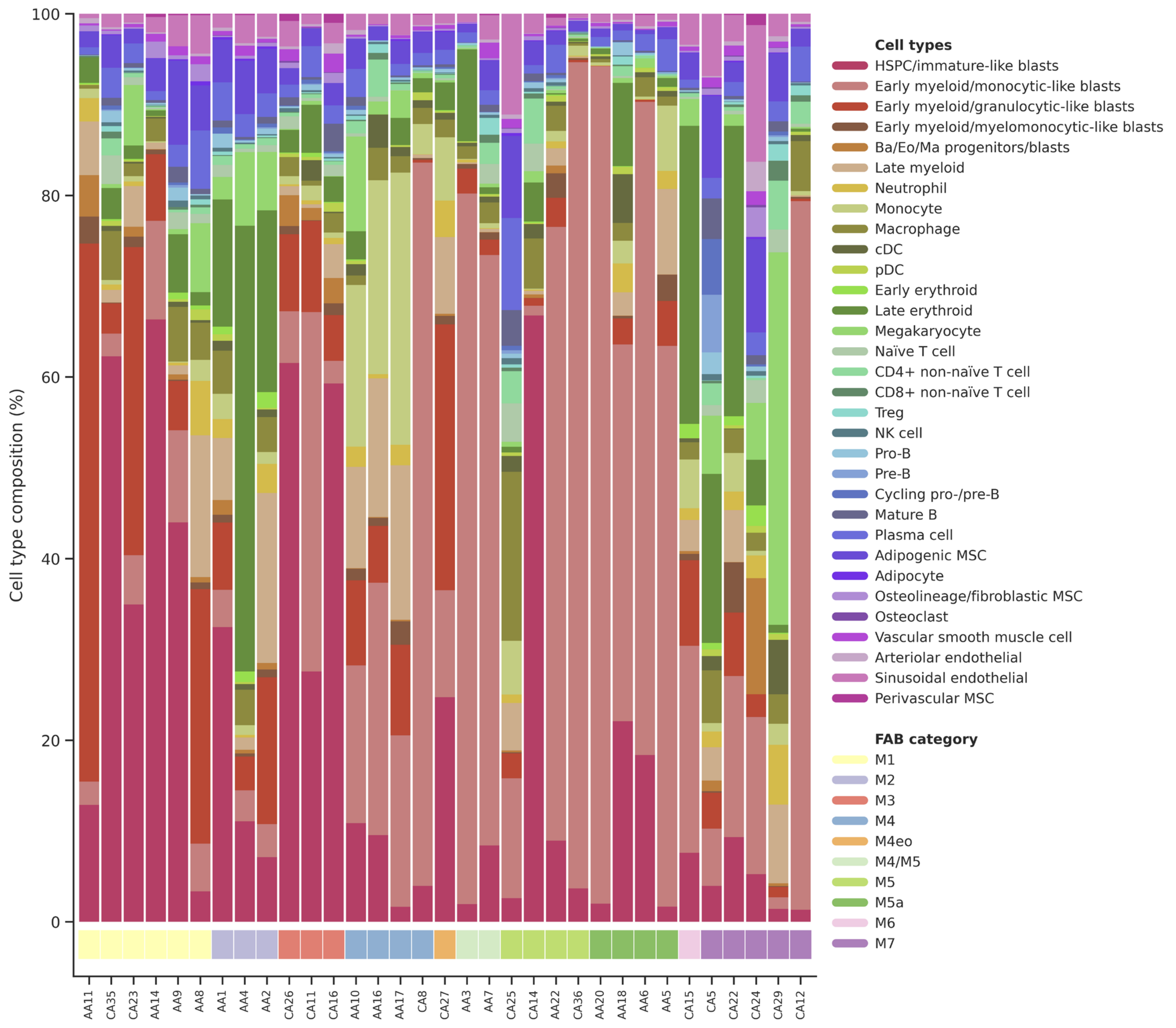

### Figure S2

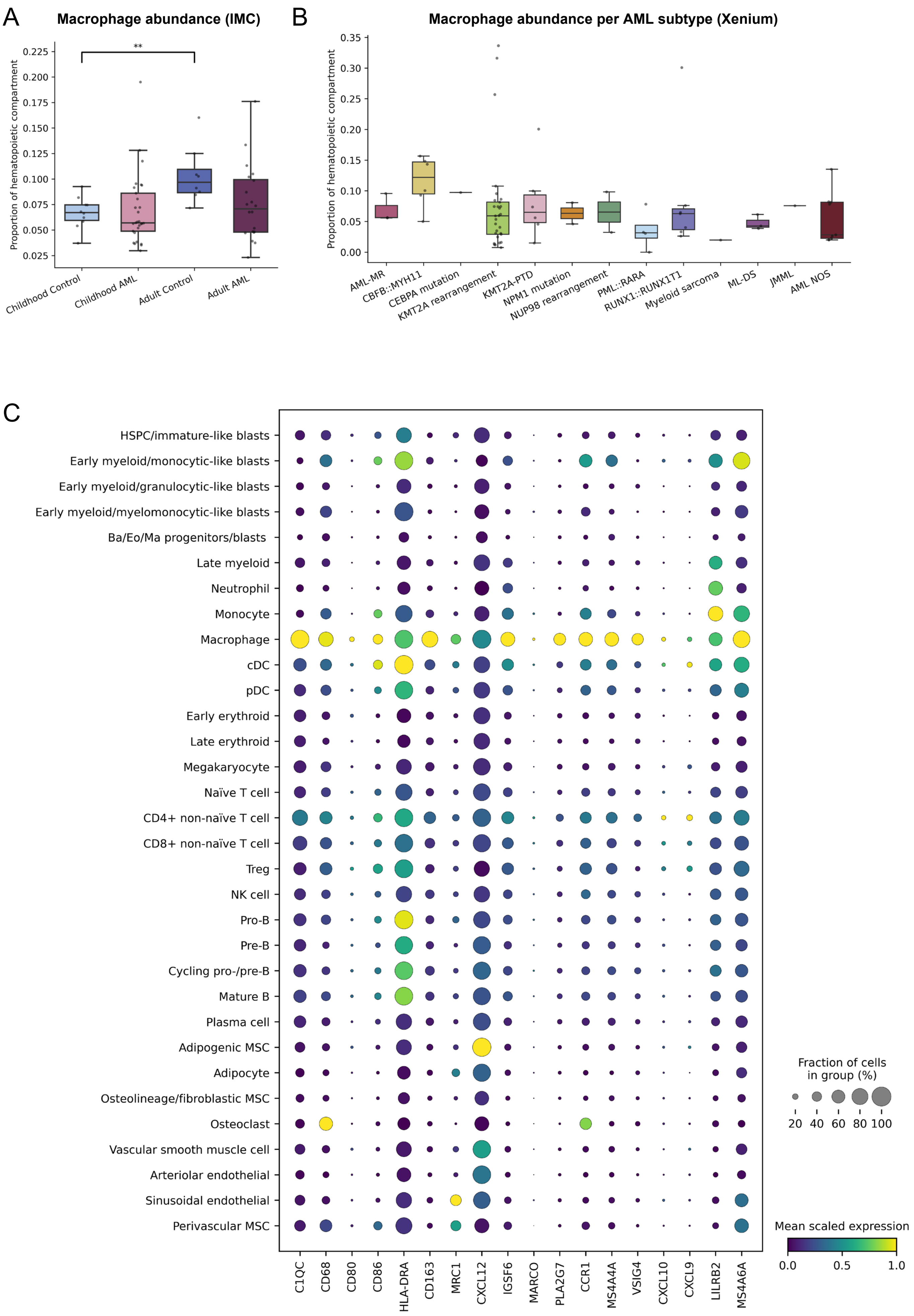

A
Macrophage abundance (IMC)
Proportion of hematopoietic compartment
**
Childhood Control
Childhood AML
Adult Control
Adult AML
B
Macrophage abundance per AML subtype (Xenium)
Proportion of hematopoietic compartment
AML-MR
CBFB::MYH11
CEBPA mutation
KMT2A rearrangement
KMT2A-PTD
NPM1 mutation
NUP98 rearrangement
PML::RARA
RUNX1::RUNX1T1
Myeloid sarcoma
ML-DS
JMML
AML NOS
C
HSPC/immature-like blasts
Early myeloid/monocytic-like blasts
Early myeloid/granulocytic-like blasts
Early myeloid/myelomonocytic-like blasts
Ba/Eo/Ma progenitors/blasts
Late myeloid
Neutrophil
Monocyte
Macrophage
cDC
pDC
Early erythroid
Late erythroid
Megakaryocyte
Naïve T cell
CD4+ non-naïve T cell
CD8+ non-naïve T cell
Treg
NK cell
Pro-B
Pre-B
Cycling pro-/pre-B
Mature B
Plasma cell
Adipogenic MSC
Adipocyte
Osteolineage/fibroblastic MSC
Osteoclast
Vascular smooth muscle cell
Arteriolar endothelial
Sinusoidal endothelial
Perivascular MSC
C1QC
CD68
CD80
CD86
HLA-DRA
CD163
MRC1
CXCL12
IGSF6
MARCO
PLA2G7
CCR1
MS4A4A
VSIG4
CXCL10
CXCL9
LILRB2
MS4A6A
Fraction of cells
in group (%)
20 40 60 80 100
Mean scaled expression
0.0 0.5 1.0

### Figure S2 (continued)

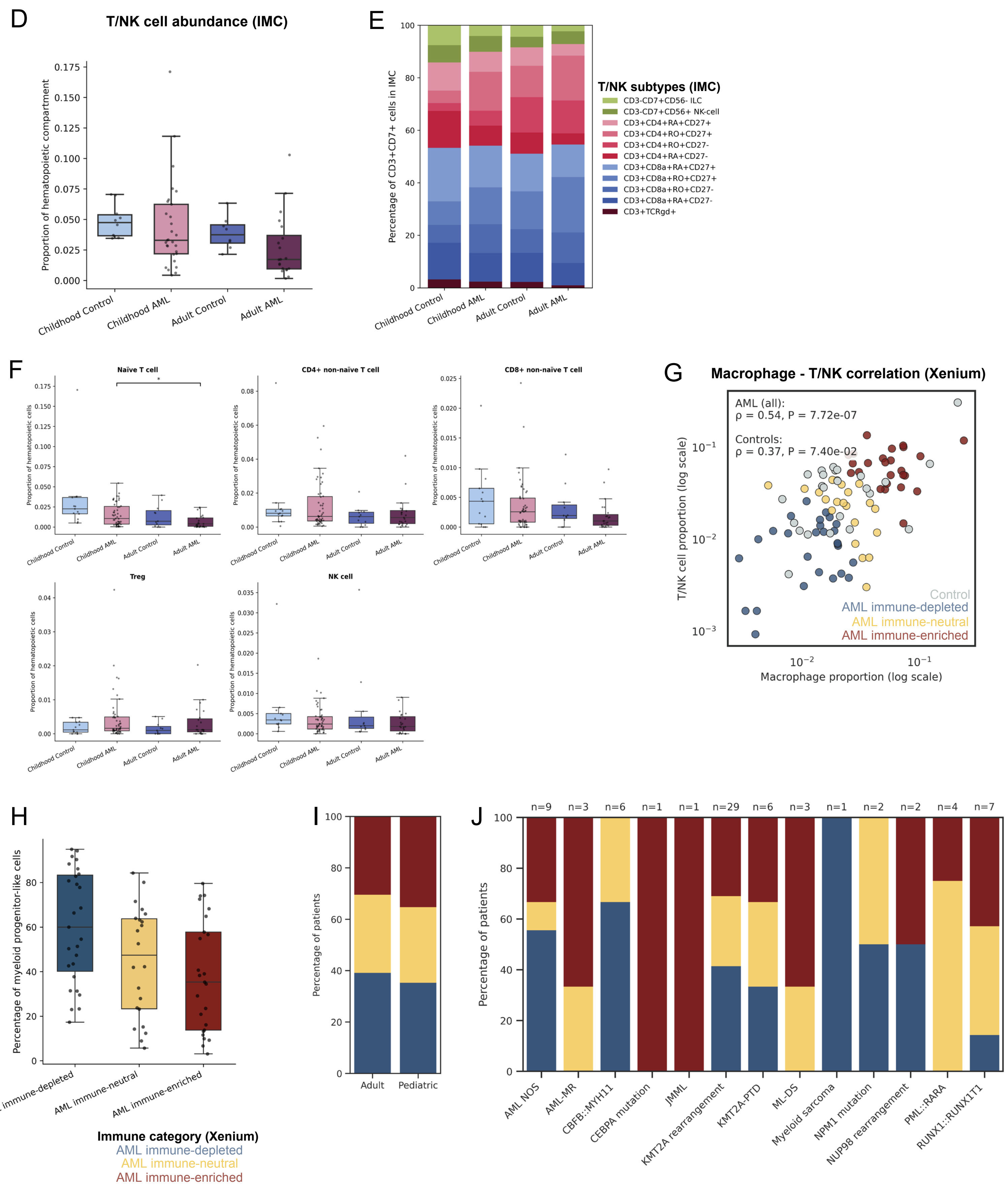

### Figure S2 (continued)

K

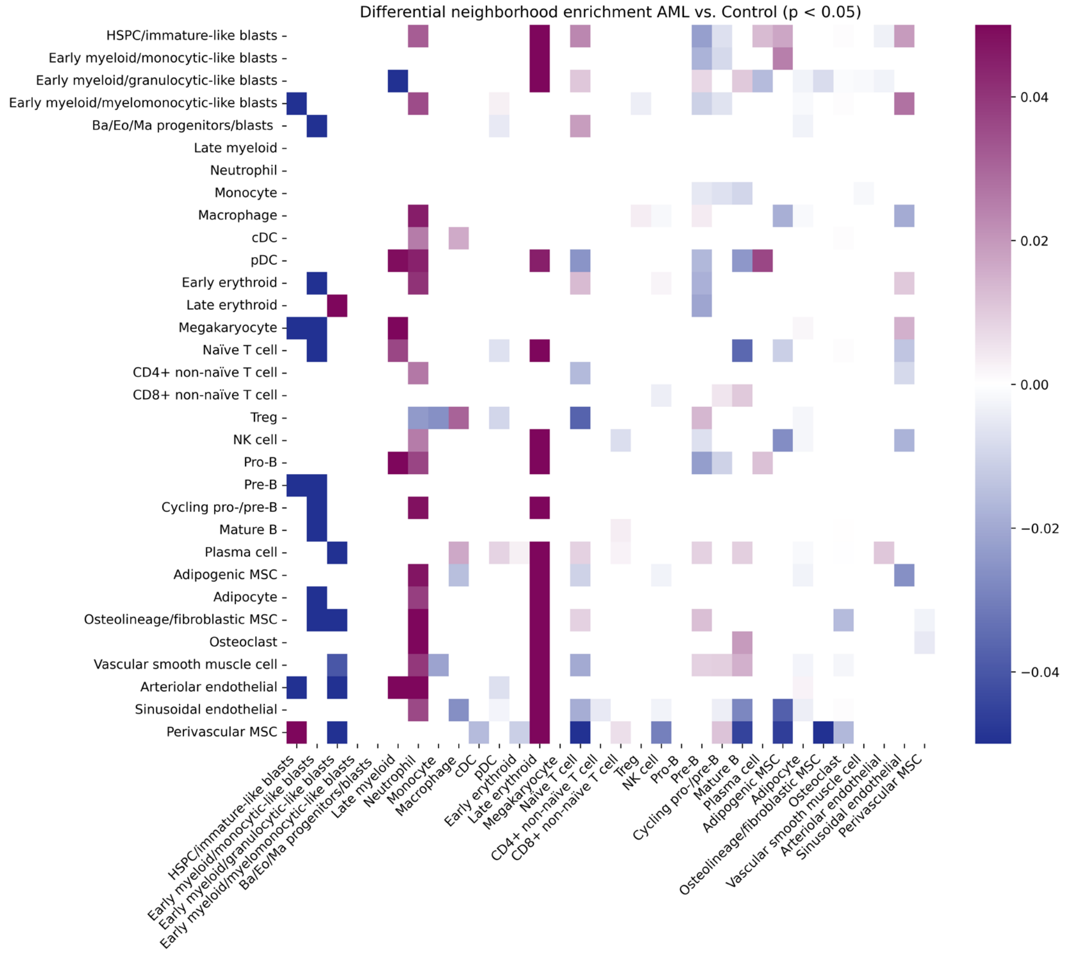

Figure S2 (continued)

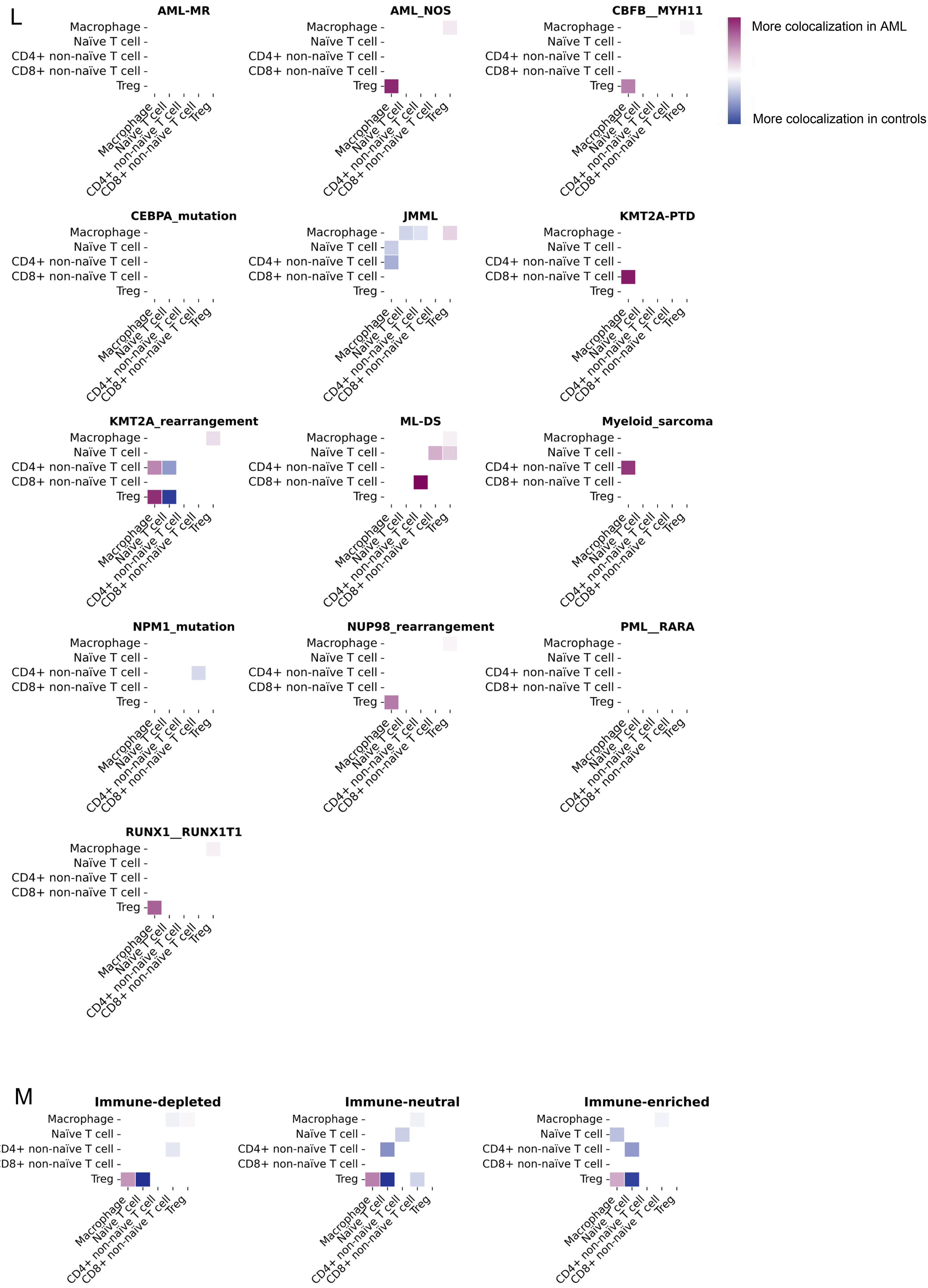

### Figure S3

A

B

C

D

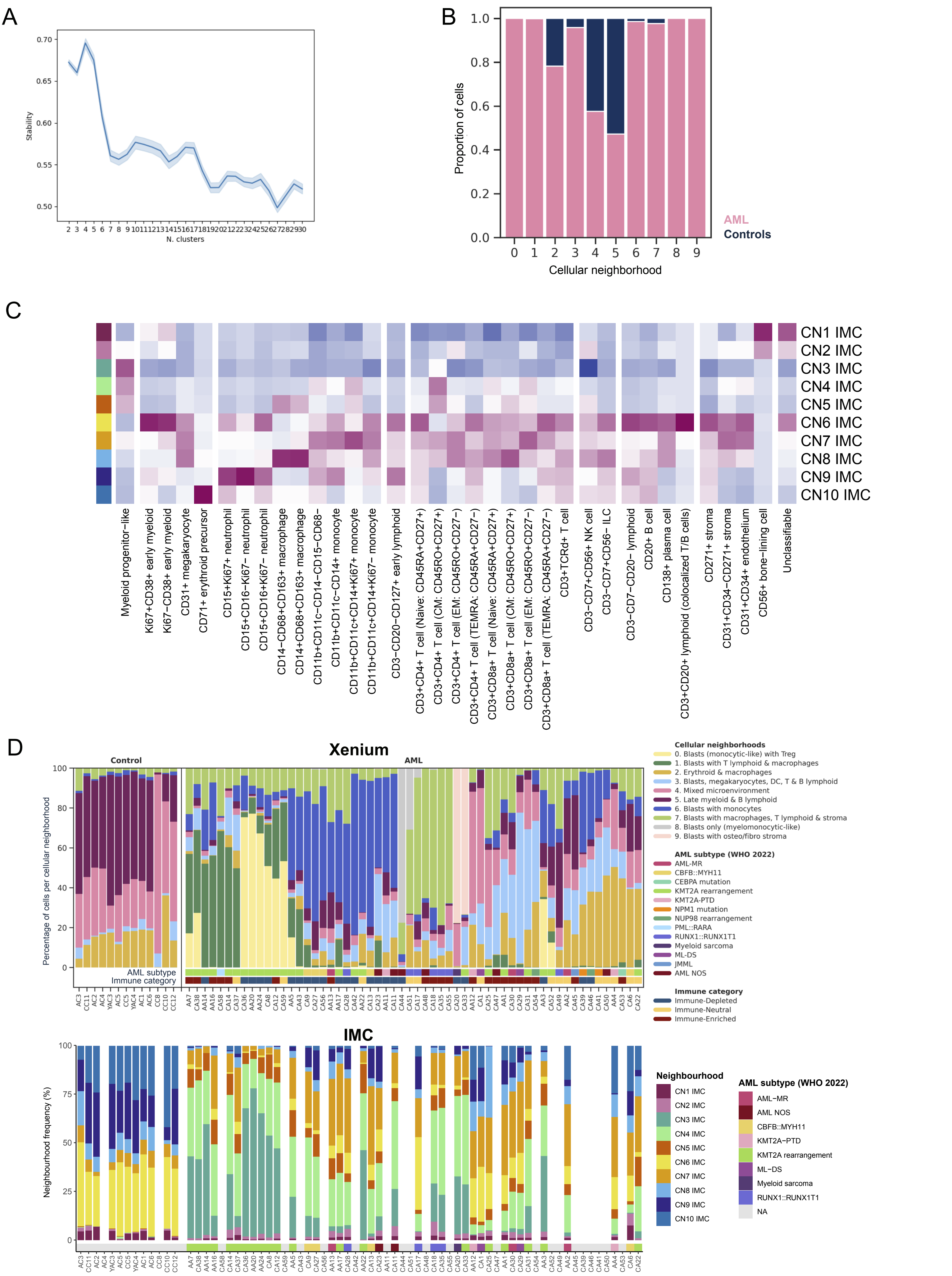

### Figure S3 (continued)

E

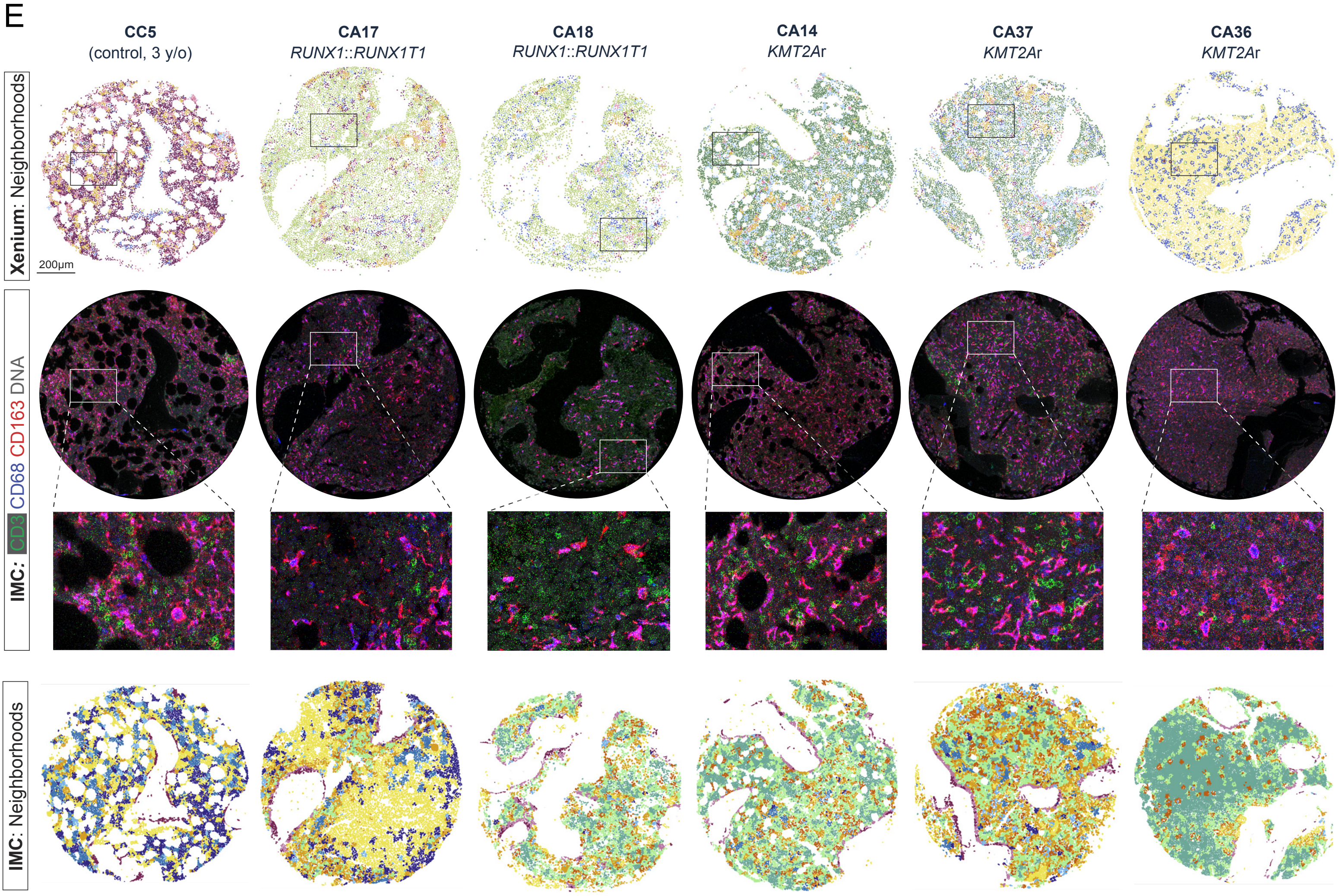


F

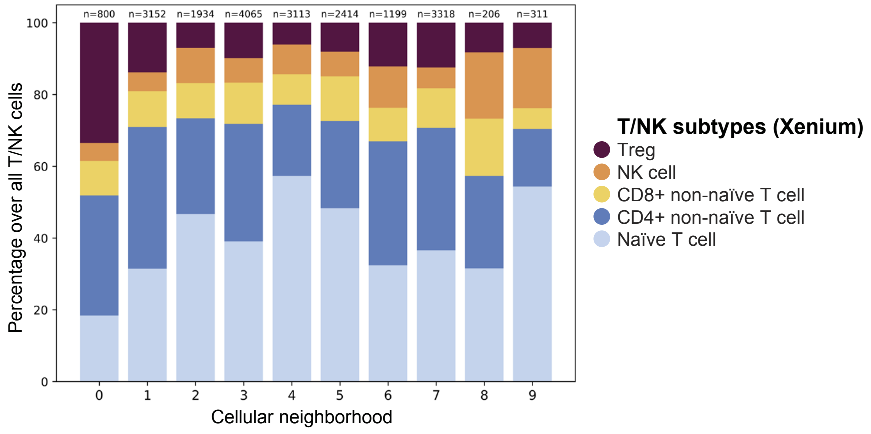

**Xenium | 2-hop neighbors**

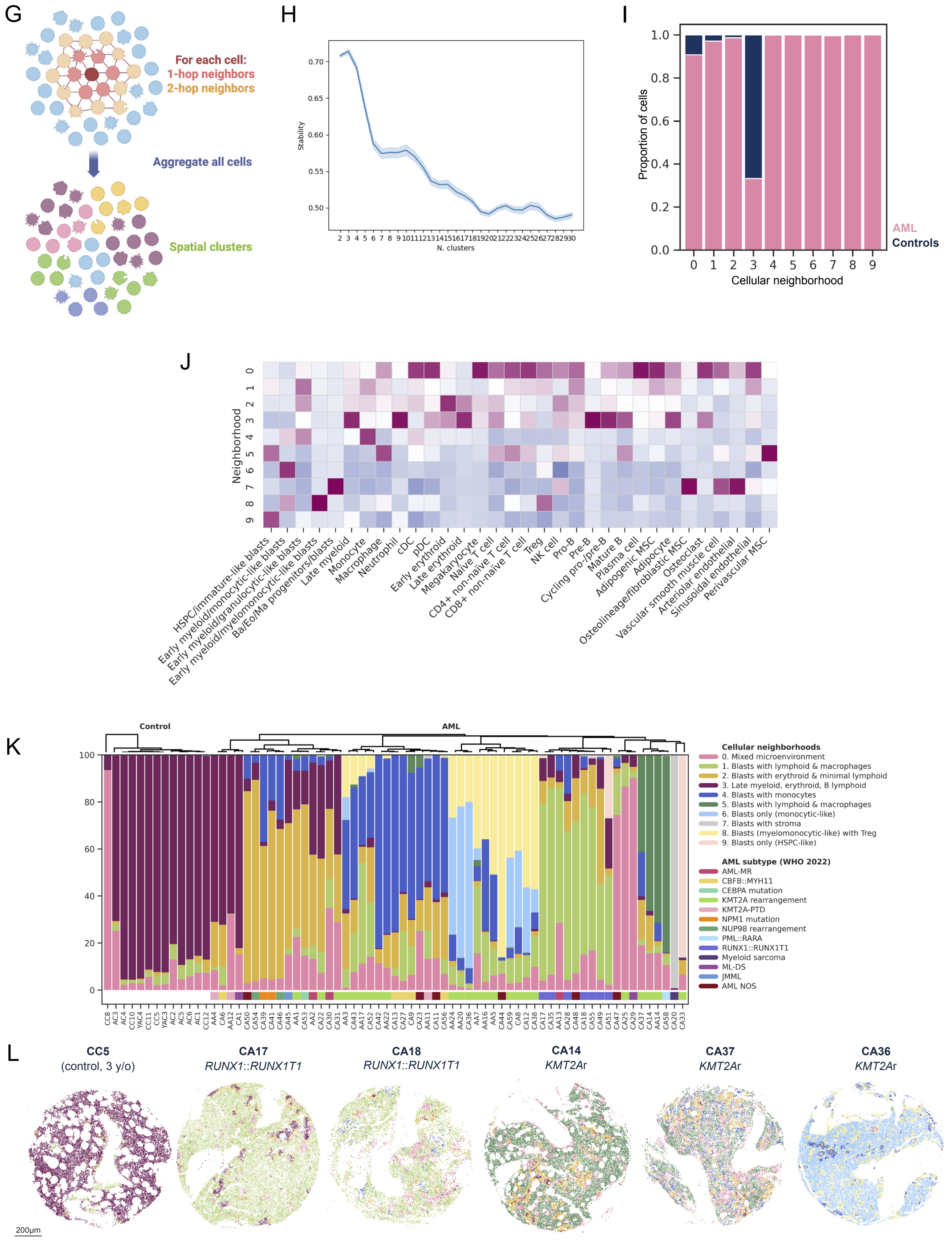

#### Xenium | 3-hop neighbors

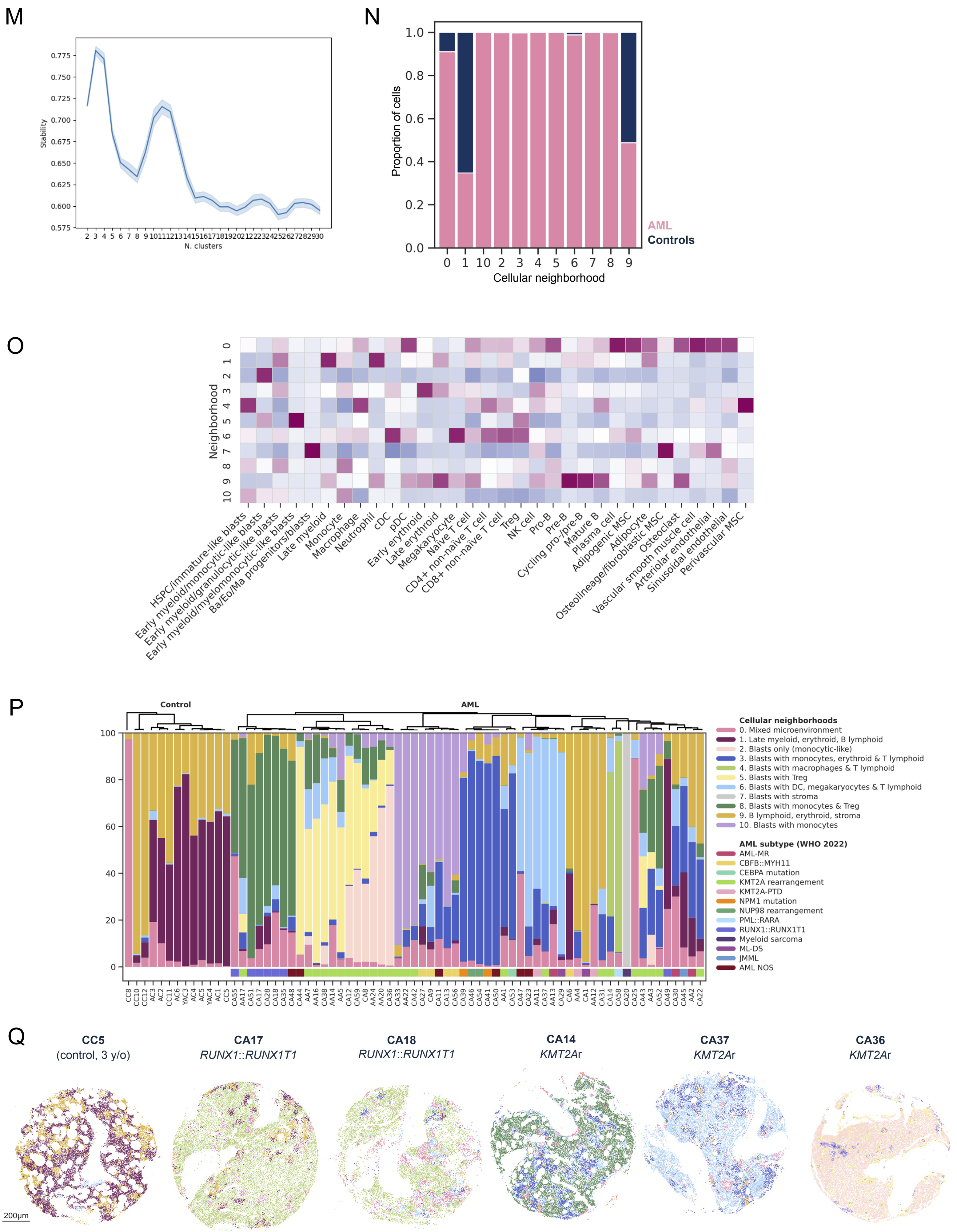

#### Figure S4

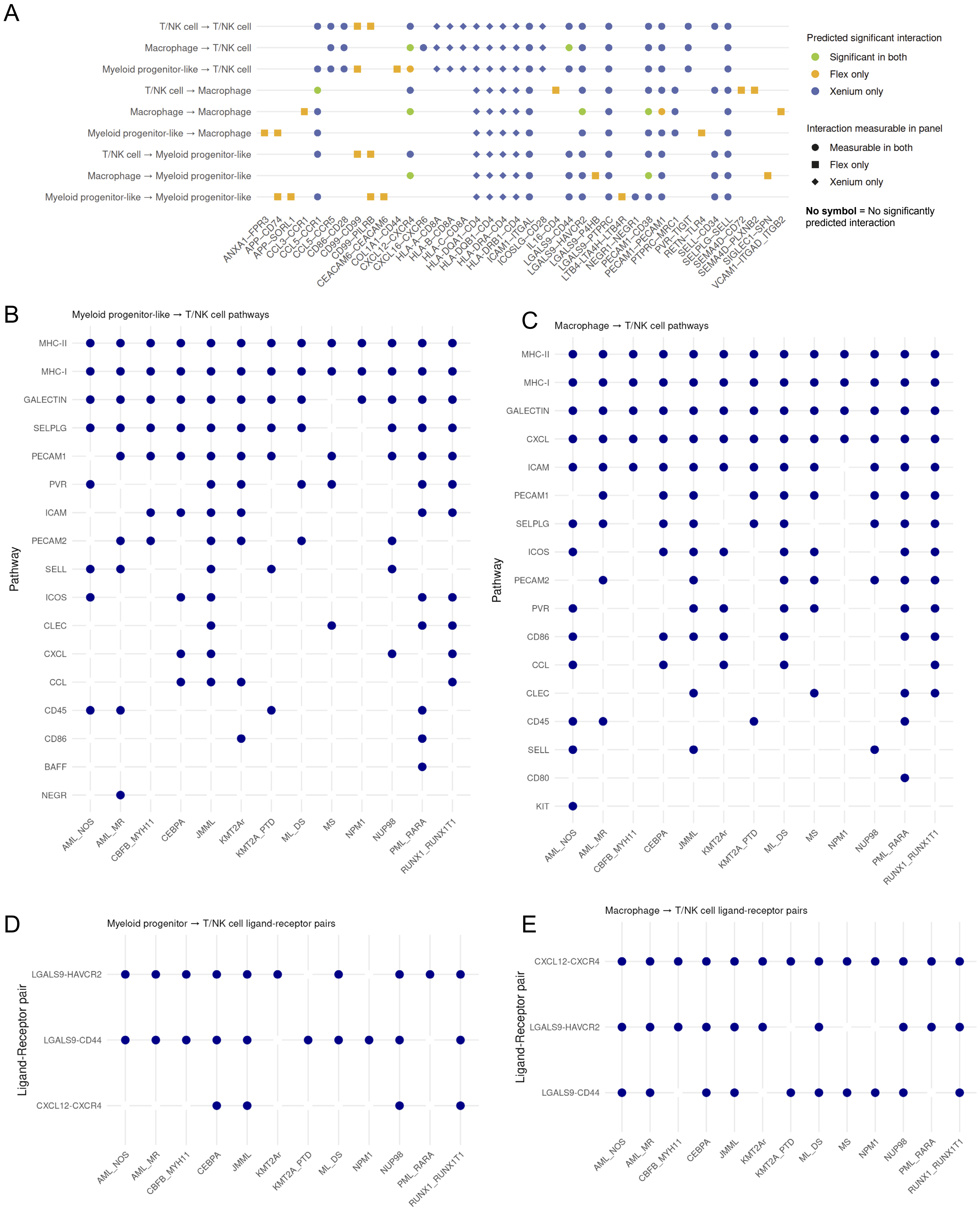
A
T/NK cell → T/NK cell
Macrophage → T/NK cell
Myeloid progenitor-like → T/NK cell
T/NK cell → Macrophage
Macrophage → Macrophage
Myeloid progenitor-like → Macrophage
T/NK cell → Myeloid progenitor-like
Macrophage → Myeloid progenitor-like
Myeloid progenitor-like → Myeloid progenitor-like
ANXA1–FPR3
APP–CD74
APP–SORL1
CCL3–CCR1
CCL5–CCR1
CCL5–CCR5
CD86–CD28
CD99–CD99
CD99–PILRB
CEACAM6–CEACAM6
COL1A1–CD44
CXCL12–CXCR4
CXCL16–CXCR6
HLA-A–CD8A
HLA-B–CD8A
HLA-C–CD8A
HLA-DQA1–CD4
HLA-DQB1–CD4
HLA-DRA–CD4
HLA-DRB1–CD4
ICAM1–ITGAL
ICOSLG–CD28
IL16–CD4
LGALS9–CD44
LGALS9–HAVCR2
LGALS9–P4HB
LGALS9–PTPRC
LTB4–LTA4H–LTB4R
NEGR1–NEGR1
PECAM1–CD38
PECAM1–PECAM1
PTPRC–MRC1
PVR–TIGIT
RETN–TLR4
SELL–CD34
SELPLG–SELL
SEMA4D–CD72
SEMA4D–PLXNB2
SIGLEC1–SPN
VCAM1–ITGAD_ITGB2
Predicted significant interaction
Significant in both
Flex only
Xenium only
Interaction measurable in panel
Measurable in both
Flex only
Xenium only
No symbol = No significantly predicted interaction
B
Myeloid progenitor-like → T/NK cell pathways
Pathway
MHC-II
MHC-I
GALECTIN
SELPLG
PECAM1
PVR
ICAM
PECAM2
SELL
ICOS
CLEC
CXCL
CCL
CD45
CD86
BAFF
NEGR
AML_NOS
AML_MR
CBFB_MYH11
CEBPA
JMML
KMT2Ar
KMT2A_PTD
ML_DS
MS
NPM1
NUP98
PML_RARA
RUNX1_RUNX1T1
C
Macrophage → T/NK cell pathways
Pathway
MHC-II
MHC-I
GALECTIN
CXCL
ICAM
PECAM1
SELPLG
ICOS
PECAM2
PVR
CD86
CCL
CLEC
CD45
SELL
CD80
KIT
AML_NOS
AML_MR
CBFB_MYH11
CEBPA
JMML
KMT2Ar
KMT2A_PTD
ML_DS
MS
NPM1
NUP98
PML_RARA
RUNX1_RUNX1T1
D
Myeloid progenitor → T/NK cell ligand-receptor pairs
Ligand-Receptor pair
LGALS9-HAVCR2
LGALS9-CD44
CXCL12-CXCR4
AML_NOS
AML_MR
CBFB_MYH11
CEBPA
JMML
KMT2Ar
KMT2A_PTD
ML_DS
NPM1
NUP98
PML_RARA
RUNX1_RUNX1T1
E
Macrophage → T/NK cell ligand-receptor pairs
Ligand-Receptor pair
CXCL12-CXCR4
LGALS9-HAVCR2
LGALS9-CD44
AML_NOS
AML_MR
CBFB_MYH11
CEBPA
JMML
KMT2Ar
KMT2A_PTD
ML_DS
MS
NPM1
NUP98
PML_RARA
RUNX1_RUNX1T1

### Figure S4 (continued)

F

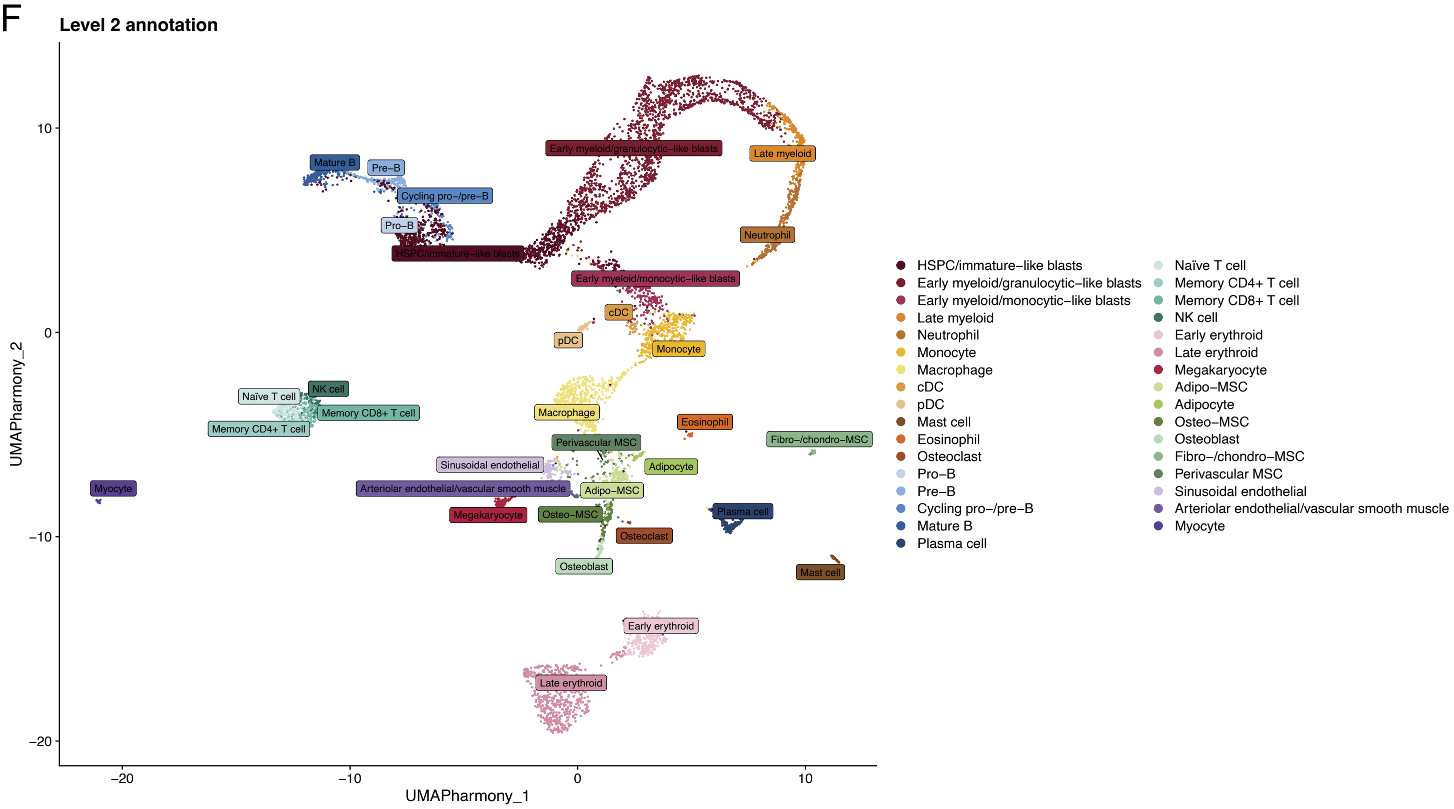


G

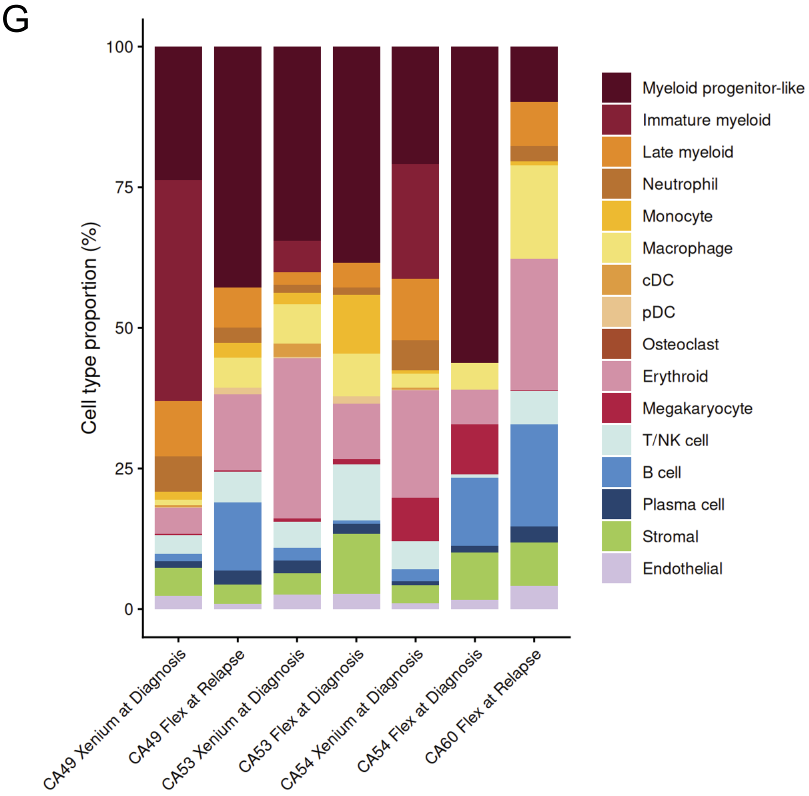

### Figure S4 (continued)

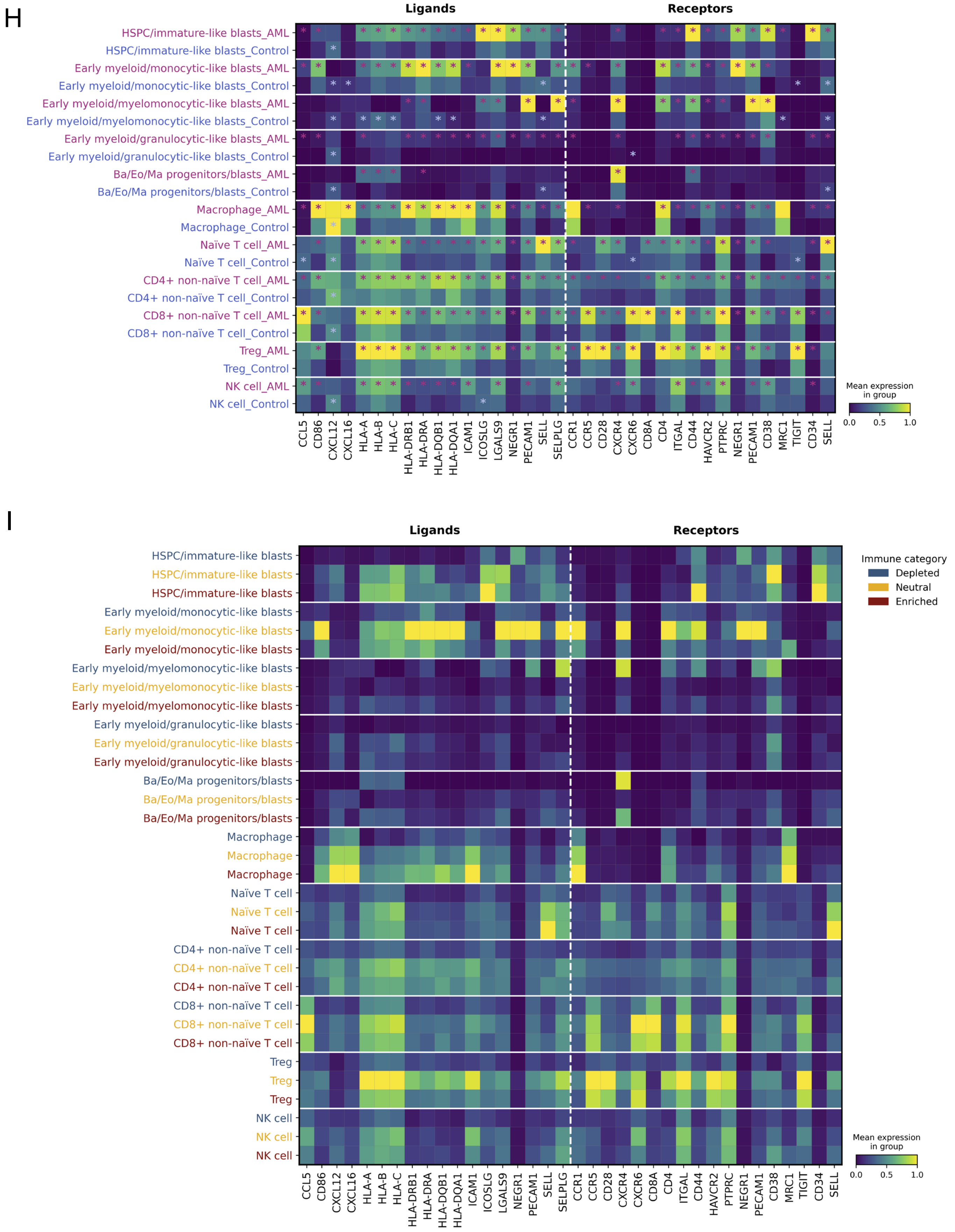

### Figure S5

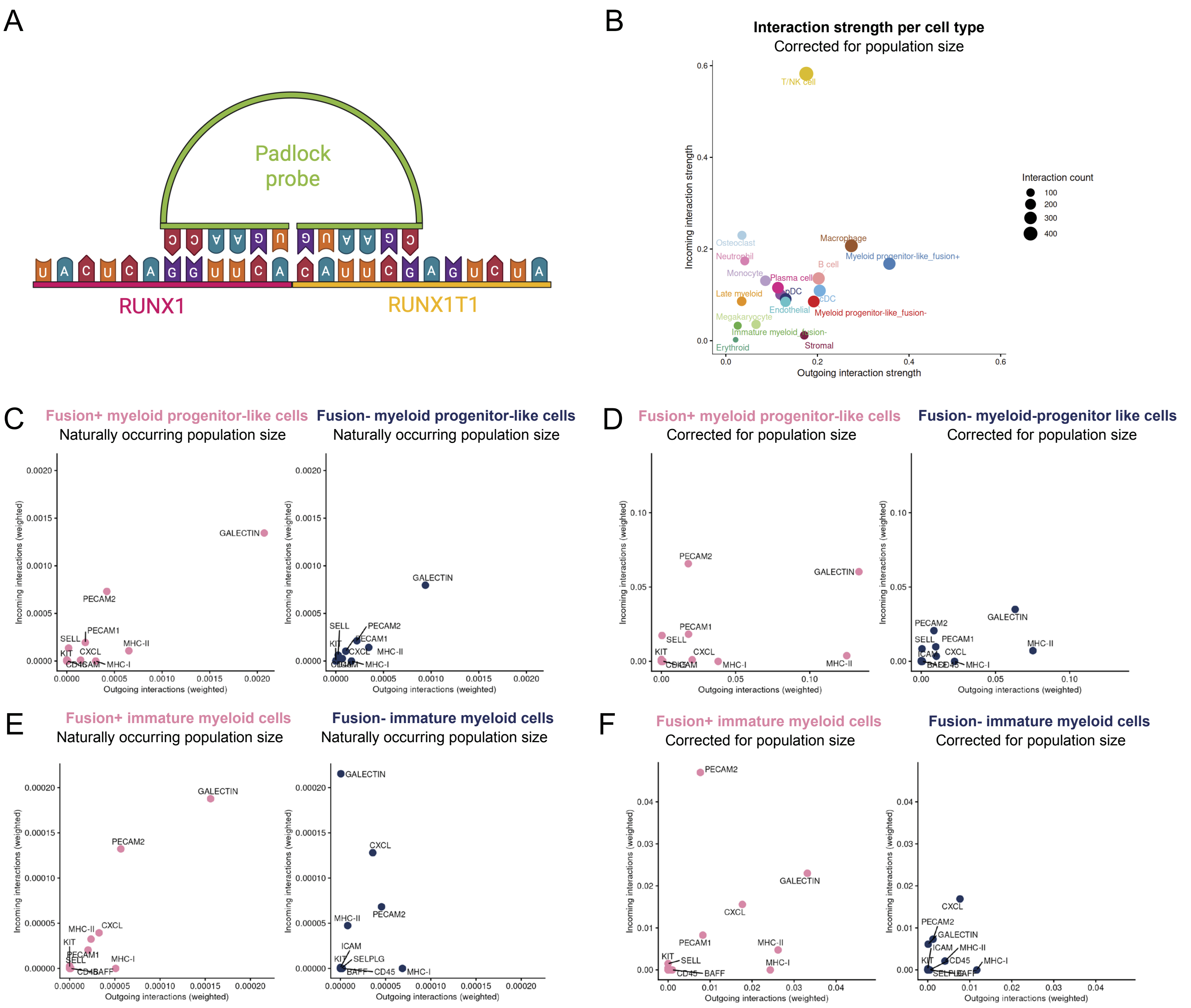

A
Padlock probe
RUNX1
RUNX1T1
B
Interaction strength per cell type
Corrected for population size
Incoming interaction strength
Outgoing interaction strength
Interaction count
T/NK cell
Osteoclast
Neutrophil
Monocyte
Plasma cell
Macrophage
Myeloid progenitor-like_fusion+
B cell
Late myeloid
pDC
cDC
Endothelial
Megakaryocyte
Myeloid progenitor-like_fusion-
Immature myeloid_fusion-
Stromal
Erythroid
C
Fusion+ myeloid progenitor-like cells
Naturally occurring population size
Fusion- myeloid progenitor-like cells
Naturally occurring population size
D
Fusion+ myeloid progenitor-like cells
Corrected for population size
Fusion- myeloid-progenitor like cells
Corrected for population size
E
Fusion+ immature myeloid cells
Naturally occurring population size
Fusion- immature myeloid cells
Naturally occurring population size
F
Fusion+ immature myeloid cells
Corrected for population size
Fusion- immature myeloid cells
Corrected for population size
Incoming interactions (weighted)
Outgoing interactions (weighted)
GALECTIN
PECAM2
PECAM1
SELL
KIT
CXCL
MHC-II
MHC-I
CD45
ICAM
BAFF
SELPLG

### Figure S6

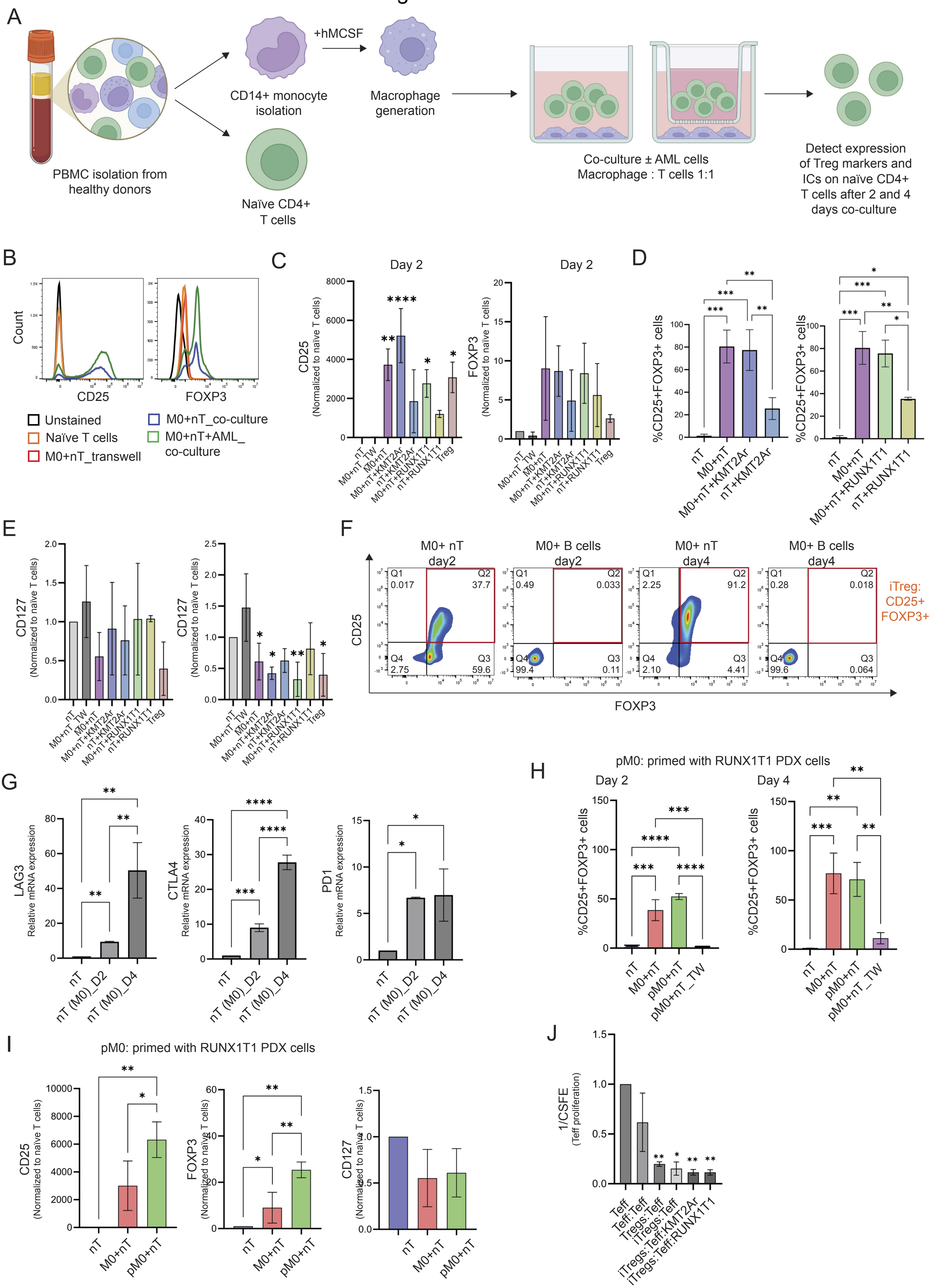

A
PBMC isolation from healthy donors
CD14+ monocyte isolation
+hMCSF
Macrophage generation
Naïve CD4+ T cells
Co-culture ± AML cells
Macrophage : T cells 1:1
Detect expression of Treg markers and ICs on naïve CD4+ T cells after 2 and 4 days co-culture
B
Count
CD25
FOXP3
Unstained
Naïve T cells
M0+nT_transwell
M0+nT_co-culture
M0+nT+AML_co-culture
C
Day 2
CD25 (Normalized to naïve T cells)
FOXP3 (Normalized to naïve T cells)
nT
M0+nT_TW
M0+nT
M0+nT+KMT2Ar
nT+KMT2Ar
M0+nT+RUNX1T1
nT+RUNX1T1
Treg
D
%CD25+FOXP3+ cells
nT
M0+nT
M0+nT+KMT2Ar
nT+KMT2Ar
M0+nT+RUNX1T1
nT+RUNX1T1
E
CD127 (Normalized to naïve T cells)
F
M0+ nT day2
M0+ B cells day2
M0+ nT day4
M0+ B cells day4
Q1 0.017
Q2 37.7
Q4 2.75
Q3 59.6
Q1 0.49
Q2 0.033
Q4 99.4
Q3 0.11
Q1 2.25
Q2 91.2
Q4 2.10
Q3 4.41
Q1 0.28
Q2 0.018
Q4 99.6
Q3 0.064
iTreg: CD25+ FOXP3+
CD25
FOXP3
G
LAG3 Relative mRNA expression
CTLA4 Relative mRNA expression
PD1 Relative mRNA expression
nT
nT (M0)_D2
nT (M0)_D4
H
pM0: primed with RUNX1T1 PDX cells
Day 2
Day 4
%CD25+FOXP3+ cells
nT
M0+nT
pM0+nT
pM0+nT_TW
I
pM0: primed with RUNX1T1 PDX cells
CD25 (Normalized to naïve T cells)
FOXP3 (Normalized to naïve T cells)
CD127 (Normalized to naïve T cells)
nT
M0+nT
pM0+nT
J
1/CSFE (Teff proliferation)
Teff
Teff:Teff
Tregs:Teff
iTregs:Teff
iTregs:Teff:KMT2Ar
iTregs:Teff:RUNX1T1

Figure S7

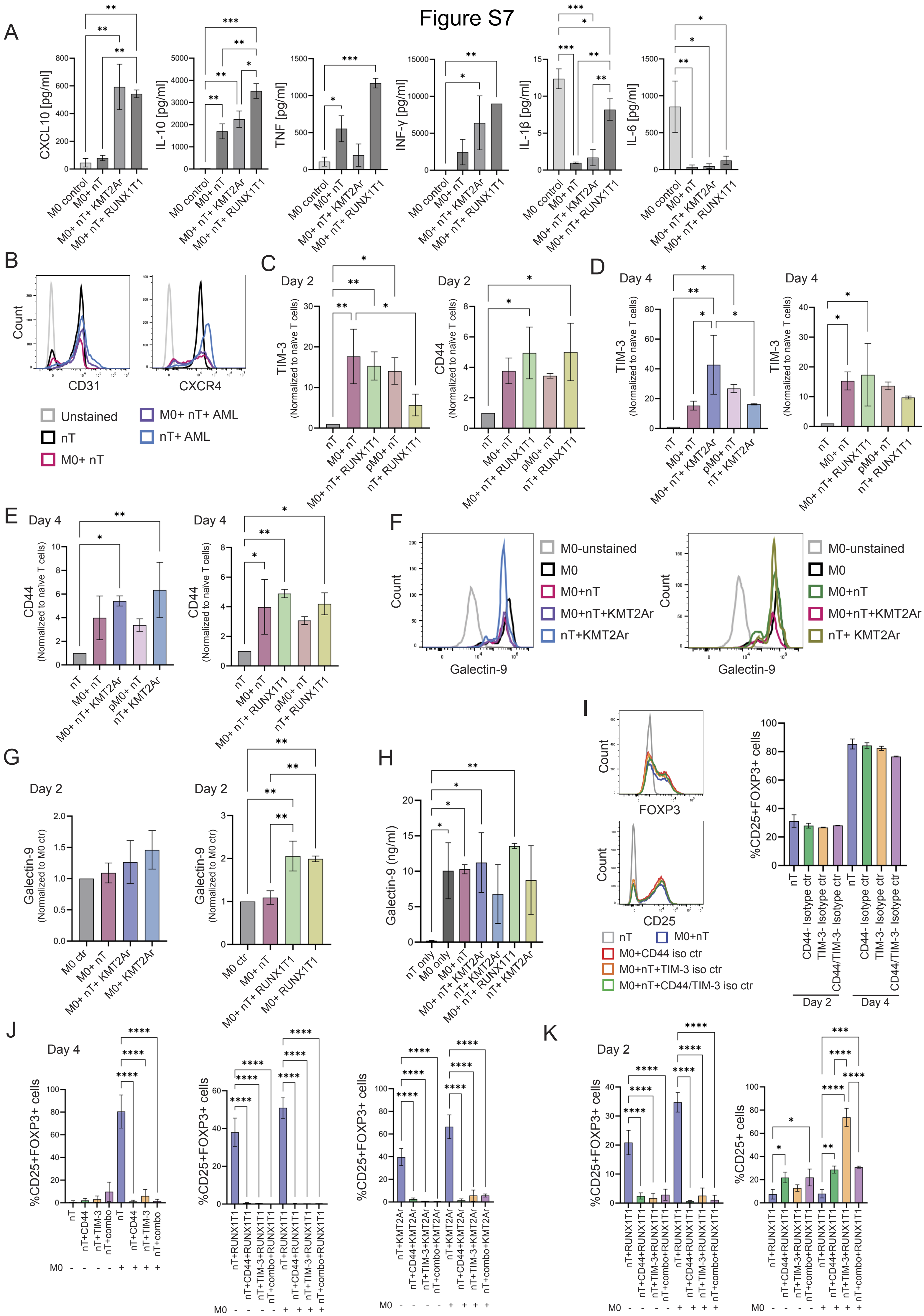
A
CXCL10 [pg/ml]
IL-10 [pg/ml]
TNF [pg/ml]
INF-γ [pg/ml]
IL-1β [pg/ml]
IL-6 [pg/ml]
M0 control
M0+ nT
M0+ nT+ KMT2Ar
M0+ nT+ RUNX1T1
B
Count
CD31
CXCR4
Unstained
nT
M0+ nT
M0+ nT+ AML
nT+ AML
C
Day 2
TIM-3 (Normalized to naïve T cells)
CD44 (Normalized to naïve T cells)
D
Day 4
E
F
Galectin-9
M0-unstained
M0
M0+nT
M0+nT+KMT2Ar
nT+KMT2Ar
nT+ KMT2Ar
G
Galectin-9 (Normalized to M0 ctr)
H
Galectin-9 (ng/ml)
I
FOXP3
CD25
M0+CD44 iso ctr
M0+nT+TIM-3 iso ctr
M0+nT+CD44/TIM-3 iso ctr
%CD25+FOXP3+ cells
J
K
%CD25+ cells

### Figure S8

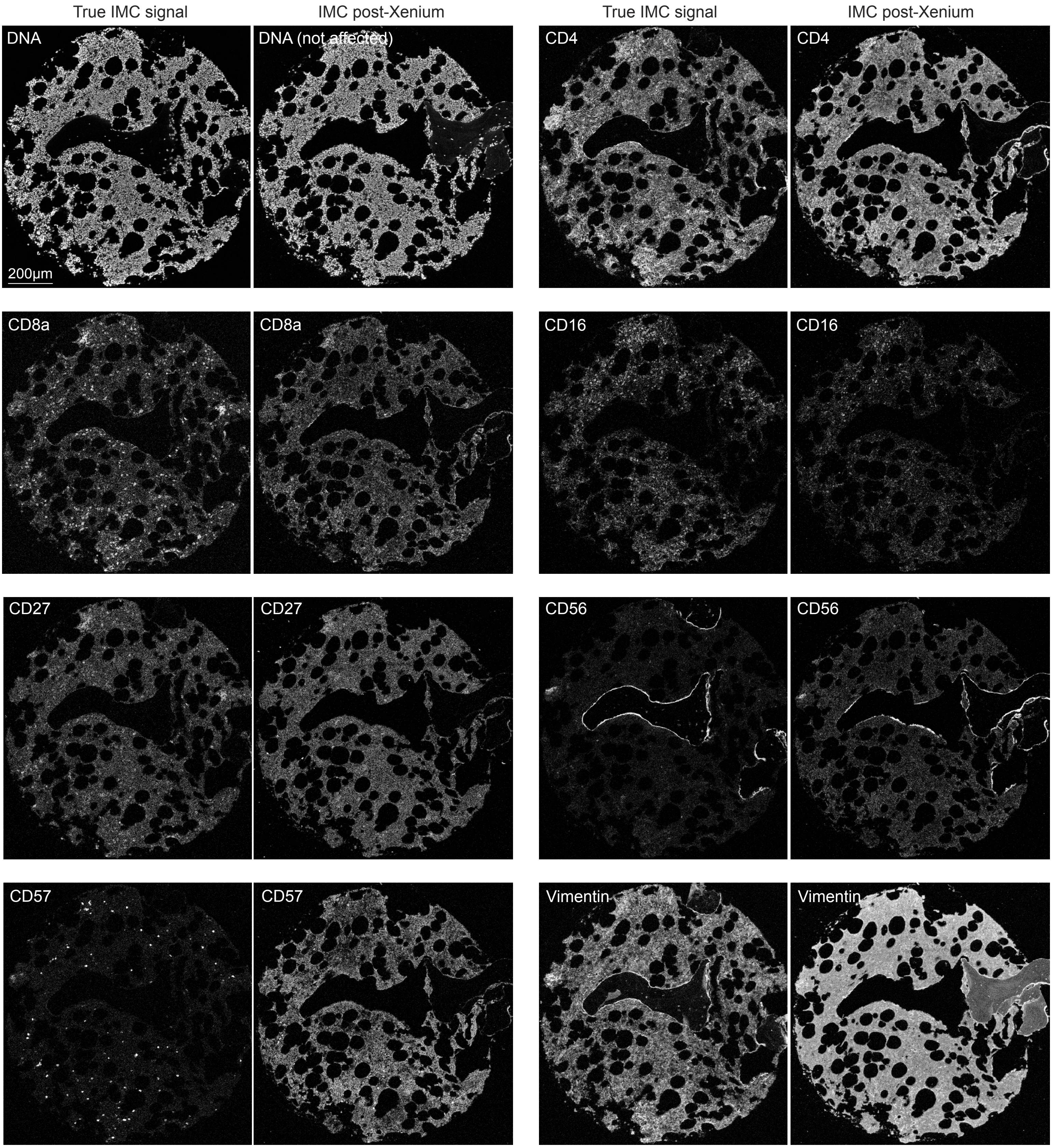
True IMC signal
IMC post-Xenium
True IMC signal
IMC post-Xenium
DNA
DNA (not affected)
CD4
CD4
200µm
CD8a
CD8a
CD16
CD16
CD27
CD27
CD56
CD56
CD57
CD57
Vimentin
Vimentin
