## Supplemental figures, legends, methods and tables for "Spatial multi-omics reveals targetable immunosuppressive macrophage T-cell interactions in human AML bone marrow": Spatial AML_Supplemental files and figure legends_bioRxiv.pdf

**Supplemental data to the manuscript 'Spatial multi-omics uncovers targetable macrophage-T cell interactions in pediatric and adult acute myeloid leukemia'**

### Supplemental methods

*Human BM biopsy selection and tissue preparation for spatial experiments*

BM biopsies used for this study were collected in routine clinical care. Ethical approval for these experiments was obtained from the Institutional Review Board of the Princess Máxima Center (PMCLAB2021.0207), Leiden University Medical Center (RP24.040), and Charité-Universitätsmedizin Berlin (EA2/013/24), in accordance with the Declaration of Helsinki.

Since a BM biopsy is an invasive procedure not often performed on individuals without malignancies or hematological conditions, the samples of the non-leukemic controls were obtained from patients undergoing tumor staging for solid malignancies, where pathological examination confirmed absence of BM involvement. Samples were provided by the biobanks of the Princess Máxima Center (Utrecht, the Netherlands), Leiden University Medical Center (Leiden, the Netherlands) and the Charité-Universitätsmedizin (Berlin, Germany), and by the Dutch Pathology Registry PALGA. All BM biopsies were fixed in formalin, decalcified using EDTA, and embedded in paraffin (FFPE) according to the local protocols: at the Princess Máxima Center using 20% EDTA for 10 hours, at the Charité using 25% EDTA for 8 hours, and at Leiden University Medical Center using 10% EDTA for 16 hours. Of note, the cohort included one BM biopsy of a patient with a myeloid sarcoma with BM involvement (indicated with as myeloid sarcoma in the figures).

Together with a board-certified and specialized haematopathologist, all available samples of AML at diagnosis (before start of therapy), were assessed for tissue quality and 1 to 3 regions of interest of 1 mm diameter each were selected per biopsy. In total, 113 individuals were selected, and 148 biopsy cores were incorporated into 3 tissue microarrays (TMA). From each TMA, consecutive 5-μm and 2-μm sections were prepared according to the Demonstrated Protocols Xenium *In Situ* for FFPE - Tissue Preparation Guide (CG000578, 10x Genomics) and the Xenium *In Situ* for FFPE Tissues – Deparaffinization & Decrosslinking (CG000580, 10x Genomics) protocol. One of the 5-μm sections was used for Xenium (performed on all 3 TMAs), while the consecutive 5-μm section was used for IMC (performed for 2 of the 3 TMAs). The IMC section was mounted on FLEX IHC Microscope slides (Dako, cat K8020) to minimize tissue detachment during the IMC staining procedure. 2-μm sections were used for hematoxylin-eosin (H&E) staining for morphological assessment. Analysis of consecutive sections was preferred over subsequent Xenium and IMC on the same slide, as pilot experiments demonstrated that the combined approach considerably impaired IMC panel performance (Figure S8). Due to the distance between the consecutive sections, some samples show differences in the location of bone trabecula between Xenium and IMC spatial plots.

*Xenium spatial transcriptomics*

Data generation

Spatial transcriptomics data were generated using the 10x Genomics Xenium platform. The Leiden Genome Technology Center at the Leiden University Medical Center performed probe hybridization, ligation and rolling circle amplification according to the manufacturer's protocol (CG000582 Rev E, 10x Genomics), using instrument software version v2.0.1.0 for the first TMA and version v3.2.1.2 for the second and third. The Human Multi-Tissue and Cancer Panel probe set (for 377 genes) was supplemented with probes for 100 custom genes, including four translocations (*RUNX1*::*RUNX1T1, KMT2A*::*MLLT10, KMT2A*::*MLLT3, KMT2A*::*AFDN*) (Table S1), totaling 477 genes. In addition, the multimodal segmentation kit was applied, with four immunofluorescent stainings for the cell nucleus, cell membrane, and cytoplasm. The resolution of Xenium is 0.2125 μm.

Data preprocessing and normalization

The Xenium Onboard Analysis pipeline (10x Genomics, version xenium-2.0.0.10 for the first TMA and xenium-3.2.0.7 for the second and third) was used for decoding of transcripts and multimodal cell segmentation. Due to suspected non-specificity of the custom probes for *FLT3LG*, this feature was removed from the raw data using Xenium Ranger relabeling. Subsequently, the probabilistic transcript-based segmentation method Proseg[1] was applied to the data from each TMA, to refine the cell boundaries generated by the imaging-based multimodal segmentation. The individual biopsy cores were selected using Xenium Explorer (version 3) and their count matrices were imported into Scanpy (version 1.11.1). As recommended for imaging-based spatial transcriptomics[2], counts were normalized to the cell volume determined by Proseg, followed by log1p transformation.

Quality control, integration, and dimensional reduction

Quality control was performed both per biopsy core and per cell. First, visual assessment of the immunofluorescent staining and the H&E was used to classify individual biopsy cores as poor quality (if the core was lost or completely distorted), medium quality (if there was some spatial distortion but (almost) no tissue loss), or good quality. Poor quality cores were excluded from all analyses, and medium quality cores were used for cell type annotation and all non-spatial analyses. Next, from the included cores, individual cells were excluded if they had ≤10 counts and/or ≤3 transcripts with at least a count of 1 (because Proseg generates fractional counts, i.e., decimal counts instead of integers, which represent a probability of the transcript pertaining to a specific cell). Spatial analyses were only performed on the high-quality cores that had at least 30% of the original cell count left after filtering. To integrate the segmented and cleaned datasets from the 3 TMAs and mitigate batch effects, the autoencoder variational model ResolVI (scvi-tools, v1.3.1)[3] was applied, considering each TMA as a batch. This model further corrects for misassigned transcripts and generates a low-dimensional representation of the data. ResolVI hyperparameter tuning was performed using Optuna (optuna, version 4.3.0) and scIB benchmarking (scib-metrics, version 0.5.4) (selected ResolVI parameters: n_hidden=32, n_hidden_encoder=64, n_latent=10, n_layers=2, dropout_rate=0.20).

Of note, because of the difference in quality control metrics available for the different technologies, relatively more cells were excluded from downstream analysis for the transcriptomics data compared to the proteomics data. However, the transcriptomics quality control did not seem to specifically remove one cell type, since cell type proportions were similar between transcriptomics and proteomics (Figure 1C).

Visualization, clustering, and annotation

The latent space generated by ResolVI was used for subsequent visualization by UMAP and clustering using Leiden clustering (resolution 1.0). Cell type annotation was performed using both canonical marker genes and two scRNA-seq reference datasets[4-6] which were mapped onto our data using Tangram[7]. First, the major lineages were assigned as level 1 annotation. Subsequently, each lineage was subset and subclustered at different resolutions, to assign a fine-grained level 2 annotation. In the level 1 annotation, immature myeloid clusters which were patient-specific and/or had an aberrant phenotype based on protein or gene expression were annotated as myeloid progenitor-like cells. We avoided the term ‘aberrant myeloid’ because healthy myeloid progenitor cells may cluster together with the AML cells due to overlap in expression profiles with the limited gene and protein panels used in this study. Immature myeloid cells that were also detected in patients were annotated as 'immature myeloid cells'. Memory T cells could not be annotated due to a lack of specific markers in the panel. The top differentially expressed genes per cell type for the level 2 annotation are shown in Figure S1G.

Statistical and spatial analysis

The relative abundance of cell types was compared between groups of interest using a Mann-Whitney U test followed by Benjamini-Hochberg false discovery rate (BH-FDR) correction for multiple testing. Differential gene expression analysis was performed in Scanpy using a Wilcoxon rank-sum test on the volume-normalized log-transformed expression values followed by BH-FDR multiple-testing correction and selection of significantly upregulated genes (adjusted $p < 0.05$). Ripley's L spatial clustering analysis and pairwise colocalization analysis were performed using the gr.ripley and gr.nhood_enrichment functions provided by Squidpy (version 1.6.5). An adapted function of the Squidpy colocalization analysis as implemented in CellCharter (gr.diff_nhood_enrichment)[8] (version 0.3.4) was used for differential colocalization analysis between AML and controls, using 2000 permutations to calculate significance. Results of this differential colocalization analysis were visually verified by spatially plotting the relevant cell types, both in the Xenium and IMC data. CellCharter was also applied to determine cellular neighborhoods (CN) based on the log-transformed volume-normalized gene expression of the 1-hop (direct) neighbors of each cell. This (low) number of hop neighbors was chosen because the main interest of this study was the direct microenvironment of AML and direct interactions between microenvironmental cell types, instead of high-level overview of the BM organization. The number of CNs was selected based on the results of the tl.ClusterAutoK function, which calculates a measure of CN stability over a range of CN numbers. Based on the spatial transcriptomics data, the highest CN stability was observed for 4 or 10 CNs. We present the more fine-grained solution with 10 CNs because it contains more biologically relevant information. Additional analyses using the 1- and 2- hop neighbors, and the 1 to 3-hop neighbors of each cell were performed to show the robustness of these results.

Ligand-receptor analysis

Spatially informed ligand-receptor interaction analysis was performed on the included AML samples using the spatial mode of CellChat on the level 1 annotation with the following settings: conversion.factor=1, spot.size=10, type="triMean", population.size=TRUE, distance.use=TRUE, scale.distance=1, contact.dependent=FALSE. Since the spatial mode could not be applied to the controls due to insufficient statistical power, the non-spatial mode of CellChat was applied to the AML and control samples separately, followed by merging of the two CellChat objects to enable direct comparison.

Fusion transcript analysis

Of the 4 different fusion transcripts for which probes were included in the panel, only one (*RUNX1*::*RUNX1T1*) showed specific detection in the biopsies of the patients harboring this fusion. The low and more unspecific detection of the other fusion transcripts were suspected to result from challenging probe hybridization for the *KMT2A* fusion partner, since the breakpoint region is very rich in adenosine. We therefore only included data on the *RUNX1*::*RUNX1T1* fusion in downstream analyses. Since the blast percentage may affect the extent of potential leakage of fusion transcripts to surrounding cells, we calculated an individual fusion detection threshold per patient based on the 99th percentile of the fusion levels in the T lymphoid compartment, because these cells were considered least likely to have true fusion expression. Since the identified fusion-positive and fusion-negative cells were part of the same dataset, comparative analyses were performed slightly differently from the comparison between AML and controls. Differential colocalization analysis between fusion-positive and fusion-negative cells was based on the squidpy.gr.nhood_enrichment function, followed by subtraction of the calculated colocalization value for the fusion-positive version of a cell type of interest from its fusion-negative counterpart. Ligand-receptor analysis was performed using the non-spatial mode of CellChat on the subset of patients harboring the fusion, and results were plotted separately for fusion-positive and fusion-negative cells.

*Imaging mass cytometry*

Spatial proteomics was performed by IMC as previously described[9,10]. Briefly, 5 µm sections of two of three TMAs were stained with our published 37-marker panel of metal-tagged antibodies[9], designed to capture both hematopoietic and non-hematopoietic cells in their true morphological shape in their native microenvironment in the BM. The following markers were included in the IMC panel: CD3, CD7, CD4, CD8a, CD11b, CD11c, CD14, CD15, CD16, CD19, CD20, CD27, CD31, CD34, CD38, CD45, CD45RA, CD45RO, CD56, CD57, CD68, CD71, CD127, CD138, CD163, CD271 (NGFR), CCR6, KIT (CD117), Collagen I, CXCL12, FOXP3, Granzyme B, HLA-DR, Ki-67, TCRδ and Vimentin. The staining procedure was initiated by tissue deparaffinization in xylol, followed by rinsing in ethanol, and boiling for ten minutes in 1x IHC Antigen Retrieval Solution High pH (Invitrogen, cat 00495648). Subsequently, slides were cooled to room temperature, blocked for 30 minutes using Superblock Blocking Buffer (ThermoFisher, cat 37515), and stained with the panel of 37 metal-tagged antibodies over the course of three days. On the first day, slides were stained with unconjugated rabbit anti-human CD4 and mouse anti-human TCRγδ antibodies in a volume of 100μL at 4°C. Following an overnight incubation, slides were washed and incubated with anti-rabbit and anti-mouse secondary antibodies conjugated to $^{145}$Nd and $^{148}$Nd, respectively, in a volume of 100 μL at room temperature. After one hour, slides were washed, stained for half of the directly conjugated antibodies of the IMC panel in a volume of 100μL for five hours at room temperature, followed by washing and an overnight incubation with the remaining IMC antibodies in a volume of 100μL at 4°C. On the final day of the staining, slides were washed and stained with DNA intercalator-Ir (Standard BioTools, cat 201192A) in a volume of 100μL for five minutes at room temperature to detect cellular nuclei. Subsequently, slides were washed, dried, and stored at 4°C until measurement. All stainings and washes throughout the staining procedure were performed in PBS supplemented with 1% BSA and 0.05% Tween.

Imaging mass cytometry data acquisition

To acquire high-dimensional IMC data from the two TMAs, stained tissue sections were measured using the Hyperion Imaging System (Standard BioTools). Regions of interest were identified visually and typically covered the full TMA tissue core (approximately 1mm$^2$). For those cores where parts of the BM tissue had detached, only the remaining intact tissue areas were included. Selected regions of interest were subsequently laser-ablated at a frequency of 200 Hz, yielding a dataset of 80 high-dimensional images visualizing the spatial marker expression patterns of all IMC antibodies at 1μm$^2$ resolution. Data inspection in MCD Viewer version 1.0.560.2 (Standard BioTools) showed clear cellular staining in all but four tissue cores divided across both patient and control groups, which were therefore excluded from comparative analyses. The markers CD19, CD59 and Collagen I did not produce the expected staining patterns and were consequently excluded from further analysis.

Segmentation of imaging mass cytometry images

To obtain single-cell information from the measured regions of interest, cell segmentation was applied on all high-dimensional IMC images as previously described, with some adjustments[9]. First, segmentation masks denoting individual cells in each region of interest were generated. To account for differences in cellular shapes and size, three separate segmentation masks were combined to create a three-layer segmentation mask, enabling precise segmentation of hematopoietic cells, spindle-shaped stromal cells, and large megakaryocytes.

To generate the cell segmentation masks, probability maps based on the signal of the DNA stain (cellular nuclei), stromal cell marker CD271 or $CD31^+Vimentin^-$ megakaryocytes were obtained for each region of interest in Ilastik version 1.3.2[11]. Subsequently, exported probability masks were loaded into CellProfiler version 4.2.1[12] to identify cell objects within each respective probability mask. Identified DNA objects were expanded by 1 pixel ($1\mu m^2$) to enable proper capture of marker expression patterns on the cell membrane in downstream analysis. Furthermore, gaps corresponding to megakaryocyte nuclei in the identified $CD31^+Vimentin^-$ megakaryocyte objects were filled using the fill holes function in CellProfiler to ensure that each megakaryocyte was segmented as one cell.

Next, identified nuclei ($DNA^+$), stromal cell ($CD271^+$) and megakaryocyte ($CD31^+Vimentin^-$) objects were compared to the raw IMC images to confirm accurate segmentation of all cell types. For each region of interest, the three sets of identified objects were then combined using the preservation strategy for overlapping objects in CellProfiler and exported as a single cell segmentation mask. Given that this study aimed to characterize the (immune) microenvironment in AML, nuclei were prioritized over stromal cell objects in case of overlap. Furthermore, megakaryocyte objects were placed on top of nuclear objects to segment each megakaryocyte as a single cell, rather than separating its cytoplasm and nucleus.

In parallel with the generation of the cell segmentation masks, PENGUIN[13], a preprocessing tool for multiplexed spatial proteomics data, was applied to optimize signal-to-noise ratios for each marker of the IMC panel by performing scaling, thresholding, normalizing and percentile filtering on the acquired images. To confirm effective and appropriate denoising, results were compared to the raw IMC stainings for each maker of the IMC panel.

Finally, the corresponding three-layer cell segmentation masks and denoised IMC images of each region of interest were aligned in ImaCytE[14]. .fcs files including the marker expression values for each cell object in the cell segmentation masks were exported for downstream analysis.

Analysis of segmented imaging mass cytometry data

.fcs files of all $7 \cdot 10^5$ segmented cell from all regions of interest were loaded into Cytosplore[15] for clustering by a three-level hierarchical stochastic neighbor embedding (HSNE) analysis with default settings (perplexity to 30 and 1000 iterations). CD19, CD59 and Collagen I were not used to generate the HSNE since these markers did not produce the expected staining patterns. At the overview level of the HSNE analysis, a total of $4 \cdot 10^4$ cells were positive or negative for all markers of the IMC panel and were accordingly excluded from further analysis. The remaining cells were selected and visualized in further detail at the second level of the HSNE analysis, which distinguished three groups of cells by Gaussian mean shift clustering. The first group resembled lymphoid cells, based on the expression of CD3, CD7, CD20, CD127 and/or CD138. The remaining cells, including mature myeloid cells, leukemic blasts and stromal cells, were divided into two groups based on their detection across multiple patients and controls. These groups of cells were therefore defined as 'shared' or 'patient-specific', with the patient-specific group most likely containing most of the leukemic blasts.

Subsequently, each group of cells was visualized in detail at the data level of the HSNE analysis. To maintain a maximum embedding of $5 \cdot 10^5$ landmarks (cells), cells within the 'shared' cell group were further divided across two groups based on marker expression. Gaussian mean shift clustering of each of the four final groups of cells at the data level identified a total of 172 phenotypically distinct cell clusters, of which the marker expression profiles were confirmed in the raw IMC images by projecting these phenotypes back onto the cell segmentation masks in ImaCytE. Subsequently, these 172 cell clusters excluding 404 cells located in the bone trabeculae were merged into 15 major cell types, or 32 subpopulations excluding patient-specific cells resembling leukemic blasts. Merging was performed based on marker expression, or cell counts to avoid small cell clusters (<50 cells) in downstream analysis. The 15 identified major cell types were defined as follows: 1) myeloid progenitor like cells displaying an aberrant phenotype or patient-specific expansion; 2) immature myeloid cells expressing CD38 while lacking all mature myeloid markers, present throughout multiple patients and controls; 3) late myeloid cells expressing CD11b while lacking other myeloid markers and an aberrant phenotype; 4) neutrophils expressing CD15 while lacking an aberrant phenotype; 5) monocytes expressing CD14 while lacking an aberrant phenotype; 6) CD68+CD163+ macrophages; 7) CD71+ erythroid cells; 8) CD34-CD31+ megakaryocytes; 9) immature lymphoid cells including CD127+ cells resembling early lymphoid cells, and CD45+CD3-CD7-CD20- cells expressing HLA-DR and CD45RA clustering with mature lymphoid populations (most likely resembling early B cells); 10) CD3+ and/or CD7+ T cells and innate lymphoid cells (ILCs) including CD56+ NK-cells; 11) CD20+ B cells; 12) CD138+ plasma cells; 13) CD56+CD271+ cells lining the bone; 14) CD56-CD271+ stromal cells; and 15) CD31+CD34+ endothelial cells.

IMC-based T-cell profiling focused on discriminating T-cell differentiation stages to complement the Xenium data, since these stages could not be fully distinguished by Xenium but could be assessed at the protein level using CD45RA, CD45RO and CD27 (annotating CD8+ and CD4+ naïve, central memory, effector memory, and terminally differentiated effector memory T cells). More detailed analyses of T-cell subpopulations, including FOXP3 expression, relied on visual inspection to capture these rarer populations within their spatial context.

The main difference in cell type frequencies between the Xenium and IMC was observed in the proportion of erythroid cells, which was higher in the Xenium data. This is likely because the IMC marker panel only included CD71 as distinctive erythroid marker, which does not identify the most immature erythroid cells, while these immature erythroid cells could be annotated in the Xenium data based on a combination of markers. Some immature erythroid cells in the IMC may have been annotated as immature myeloid in the IMC, explaining the slightly higher percentage of immature myeloid cells in the IMC.

To visualize identified cell clusters, merged .fcs files of cell clusters were loaded into OMIQ (Dotmatics) for optSNE analysis. To quantify the abundance of the identified cell clusters across individuals, merged .fcs files were loaded into R version 4.4.1 and analyzed using an adapted Cytofast pipeline script[16]. Spatially resolved cellular neighborhood analyses were performed in R using imcRtools[17] by incorporating the extracted XY coordinates (centroids) of all segmented cells obtained from CellProfiler. Based on the number of cellular neighborhoods identified by CellCharter in the Xenium data, cells were grouped into 10 cellular neighborhoods by unbiased kmeans clustering based on the phenotypes of their 10 nearest neighbors within a 30-pixel distance. Results were projected on cell segmentation masks using Cytomapper. All further analyses and data visualization were performed in R.

*10x Flex single-cell RNA profiling*

Sample selection

To validate the ligand-receptor interactions predicted based on the Xenium data, we selected four FFPE pediatric AML BM biopsies from the biobank of the Princess Máxima Center (Utrecht, the Netherlands) to undergo 10x Flex scRNA-seq. Three samples pertained to patients included in the spatial TMAs (*RUNX1*::*RUNX1T1*, *NUP98*::*TOP1*, and normal karyotype), 2 of which were matched samples at primary diagnosis, while 1 sample was retrieved at relapse. An additional relapse sample (AML with *KMT2A*::*MLLT3)* pertained to a patient whose diagnostic sample had not been included in the TMA. Since our aim was to validate ligand-receptor interactions that occur in the AML BM in general, without the need to match samples within a patient, we included all four samples for downstream analysis.

Processing of FFPE tissue samples for 10x Genomics Gene Expression Flex

The four FFPE BM biopsies were processed using the pestle dissociation protocol: Sample Preparation from FFPE Tissue Sections for Chromium Fixed RNA Profiling (CG000632, 10x Genomics). From each biopsy, 3 to 6 25-mm scrolls were used as input. The isolated cells/nuclei were stained with Acridine Orange/Propidium Iodide (AO/PI) and counted using a CellDrop fluorescent cell counter (DeNovix).

Library preparation and sequencing

Each sample was barcoded using a different probe barcode from the Chromium Human Transcriptome Probe Set v1.0.1 (BC001 – BC004) with a 21-hour probe hybridization step. A single library was generated using the manufacturer's protocol: Chromium Fixed RNA Profiling Reagent Kits (CG000527; 10x Genomics). The 10x Flex library was sequenced on an Illumina NovaSeq 6000 sequencer and subsequently processed with cellranger multi v7.2.0. A total of 14,160 cells were recovered with a mean read depth of 32,628 reads/cell.

Quality control, annotation and ligand-receptor analysis

Downstream analyses were performed in R (version 4.6.0). Cells classified as doublets by scDblFinder [18] in at least 5 out of 10 runs were excluded. DecontX[19] was used to remove ambient RNA contamination. Subsequently, cells with fewer than 500 transcripts, more than 80,000 transcripts, fewer than 500 unique genes, or over 15% of mitochondrial transcripts were discarded. Seurat (version 5.4.0) was used for all subsequent analyses. The gene expression matrix was normalized using SCTransform (v2, using 3000 variable features). From the variable features, the following genes were excluded, using gene lists from the SCutils package (version 2.2): cell cycle-related genes, genes correlating with these cell cycle-related genes in this dataset, ribosomal genes, and stress-related genes. Dimensional reduction was performed using Seurat's RunPCA function. Harmony was used to integrate the individual sample gene expression, using the IntegrateLayers function. The data was clustered using the FindNeighbors and FindClusters functions, and RunUMAP (using 30 principal components) was used for lower dimensional visualization. Cell type annotation was performed using canonical markers, focusing on the markers used for the annotation of the Xenium data to align the annotations. This was verified by mapping scRNA-seq reference datasets[4-6] onto the

Flex data using SingleR[20]. Ligand-receptor interaction analysis was performed using the standard mode of CellChat (parameters: type="triMean", population.size=TRUE).

Of note, discrepancies in cell type proportions between the Xenium and Flex data as shown in Figure S4E may be related to differences in sample processing between the technologies, biological differences between relapse and diagnostic samples of the same patient, small differences between (potentially non-consecutive) sections of the same biopsy used for the different technologies, and potentially differences in annotation due to the difference in gene panels used for both technologies.

*Validation experiments*

Human monocytes and T-cell isolation

Peripheral blood mononuclear cells (PBMCs) were purified from fresh buffy coats (Sanquin, Amsterdam, The Netherlands) of healthy donors by a density gradient centrifugation using Lymphoprep (cat. 2016-02, Stem Cell, Vancouver, BC, Canada). From the same donor, different immune cell populations were isolated by magnetic-activated cell sorting. $CD14^+$ monocytes were first isolated by positive selection using CD14 microbeads, human (cat. 30-097-052, Miltenyi Biotec, Bergisch Gladbach, Germany). Subsequently, pan T cells were isolated by negative selection from the $CD14^+$ monocytes-depleted PBMC fraction using the Pan T cell Isolation Kit, human (cat. 130-096-535, Miltenyi Biotec) according to the manufacturer's instructions. From the pan T cell population, naïve CD4+ T cells were isolated by negative selection with the Naïve CD4+ T cell Isolation Kit II, human (cat. 130-094-131, Miltenyi Biotec). Alternatively, $CD25^{high}$ T regulatory cells were isolated by positive selection with CD25 microbeads II, human (cat. 130-092-083, Miltenyi

Biotec), or CD3$^{+}$ T cells were isolated with CD3 microbeads, human (cat. 130-097-043, Miltenyi Biotec).

Generation of human monocyte-derived macrophages

Macrophages were generated from primary CD14$^{+}$ monocytes previously isolated from heathy PBMCs and cultured for 5 days with RPMI 1640 (cat. 11875093, Gibco, Waltham, MA, USA) supplemented with 10% heat-inactivated human serum (cat. H422, Sigma-Aldrich, Merck, Darmstadt, Germany) in the presence of 20 ng/ml macrophage colony-stimulating factor (M-CSF, cat. 300-25, PeproTech, Cranbury, NJ, USA). Macrophages were seeded at a density of 125,000 cells/cm$^{2}$. Medium with M-CSF was replaced at day 3 of culture.

Patient-derived xenografts

For co-culture with AML cells, two patient-derived xenografts (PDX) were selected based on the molecular diagnoses that showed increased macrophage-Treg colocalization in the Xenium data. One PDX with a *RUNX1*::*RUNX1T1* translocation was generated from an adult patient at relapse and carried additional variants in *KIT* (D816V, variant allele frequency (VAF) 48%) and *TET2* (T1393fs, VAF 94%)[21,22]. The second PDX carried a *KMT2A*::*MLLT1* translocation and was generated from a pediatric patient at diagnosis.

Co-culture of human macrophages with naïve CD4+ T cells for iTreg differentiation

Naïve $CD4^+$ T cells were added to the previously generated macrophages at a macrophage to naïve T cell ratio of 1:1 or 1:3. The cells were stimulated with 0.5 ug/ml of anti-CD3 (cat. 317325, Biolegend, San Diego, CA, USA) and 1 ug/ml of anti-CD28 (cat. 377803, Biolegend) and subsequently incubated for 2 or 4 days in co-culture or in 0.4 µm transwell settings (cat. PTHT24H48, Sigma-Aldrich, Merck, Darmstadt, Germany). In some conditions, PDX cells were added to the co-culture of macrophages and naïve $CD4^+$ T cells from day 0. In other conditions, macrophages were pre-incubated with PDX for 6 days before being co-cultured with naïve $CD4^+$ T cells. In the blocking antibody assay, anti-human CD44 (cat. 103046, Biolegend) and/or anti-human CD366 (TIM-3) (cat. 345002, Biolegend) were added to the co-culture at concentrations stated in the figures. Rat IgG2b, κ (cat. 400643, Biolegend) and/or mouse IgG1, κ (cat. 400101, Biolegend) were used as isotype controls. As a negative control, to establish whether iTreg differentiation was an artefact of the co-culture system, macrophages were co-cultured with B cells, previously isolated from healthy PBMCs by positive selection using B Cell Isolation Kit II, human (cat. 130-091-151, Miltenyi Biotec), instead of T cells.

Flow cytometry

Cell surface antigen staining was performed in the dark with antibody dilution in FACS buffer (1X PBS + 0.025% BSA + 0.02% $NaN_3$ + 1% FBS) for 20 minutes at room temperature. The cells were washed twice with FACS buffer, then analyzed using a flow cytometer or used for subsequent intracellular staining. Intracellular staining was performed with the True-Nuclear Transcription Factor Kit (cat. 424401, Biolegend), according to the manufacturer's protocol. Acquisition was performed on the CytoFLEX LX Flow Cytometer (Beckman Coulter, Brea, CA, USA), and compensation was applied when needed. FACS data were analyzed using FlowJo (LLC, USA).

RNA isolation and RT-qPCR

RNA was extracted using the Nucleospin RNA kit (cat. 740955.50, Macherey-Nagel, Dueren, Germany) according to the manufacturer's instructions. The RevertAid First Strand cDNA Synthesis Kit (cat. K1621, Thermo Fisher, Waltham, MA, USA) was used according to the supplier's protocol for reverse transcription. Real-time quantitative PCRs (RT-qPCR) were performed with the SsoAdvanced Universal SYBR Green Supermix (cat. 1725274, Bio-Rad, Hercules, CA, USA) on a CFX384 qPCR machine (Bio-Rad, Hercules, CA, USA).

T cell suppression assay

After 2 and 4 days of co-culturing macrophages with naïve $CD4^+$ T cells, macrophage-induced T regulatory cells (iTregs) were collected for suppression assays. The iTregs were labeled with the proliferation dye Cell Trace Violet (CTV, cat. C34557, Thermo Fisher) according to the manufacturer's instructions. $CD3^+$ T cells, previously isolated from healthy PBMCs, were labeled with the CFSE Proliferation Kit (cat. C34554, Thermo Fisher) to serve as effector cells. CTV-labeled iTregs and CFSE-labeled $CD3^+$ T cells were co-cultured at a 1:1 ratio in a 96-well plate coated with 1 µg/ml of anti-human CD3, in RPMI 1640 supplemented with 5% human serum and 1 µg/ml of anti-human CD28. After 5 days, cells were harvested, stained with live/dead markers, and analyzed for suppression of proliferation by flow cytometry. Negative controls consisted of $CD3^+$ T cells alone or co-cultured with other $CD3^+$ T cells, while positive controls used $CD3^+$ T cells co-cultured with $CD25^{high}$ Tregs isolated from healthy PBMCs.

Cytokine detection

To analyze cytokine and chemokine levels in cell culture supernatants collected from the co-culture between macrophages and naïve $CD4^{+}$ T cells, the LegendPlex Multiplex Immunoassay (BioLegend, San Diego, CA, USA) was performed. Either after 2 or 4 days of co-culture, samples were prepared according to the manufacturer's guidelines. Samples were centrifuged to remove any particulate matter, and supernatants were collected and stored at -80°C until analysis. The following human cytokines were detected: TNF-α, IFN-γ, IL-1β, IL-2, IL-6, IL-10, CXCL-10, IL-12p70 and active TGF-β1. Blank samples (culture medium only) and known standard concentrations were included in each assay to ensure accuracy and reproducibility. Data were acquired on the CytoFLEX LX Flow Cytometry (Beckman Coulter) and analyzed by the provided LEGENDplex Data Analysis Software Suite (BioLegend, San Diego, CA, USA).

<u>ELISA</u>

The Galectin-9 ELISA Set (cat. DGAL90, BD Biosciences, RD Systems) was used according to the manufacturer's instructions. After the colour reaction developed, absorbance was measured at 450 nm using the microplate reader SpectraMax iD3 (Molecular Devices, LLC).

<u>Migration assay</u>

Transwell inserts with a 5.0 μm pore size (cat. CLS3421, Merk) were placed into wells containing either macrophages, culture medium supplemented with 100 μg/ml of the chemoattractive cytokine CXCL12 (cat. 300-28A, PeproTech), or culture medium alone. After 1, 2, 4, 6 and 24 hours of incubation, cells that migrated through the transwell were collected and stained with antibodies against CD4 and CD25, along with a live/dead viability dye. Data acquisition was performed using the CytoFLEX LX flow cytometer (Beckman Coulter).

Statistics

Each figure legend provides details about statistical comparisons. All computations were performed using GraphPad Prism, and all P-values are 2-sided.

### References to Supplemental methods

**Supplemental Figures titles and legends**

Supplemental Figure S1

S1A. Overview of clinical characteristics per patient. NA, not applicable.

S1B. UMAP of IMC data (including the non-classifiable cells).

S1C. UMAPs of Xenium data, broad (level 1) annotation, for AML and controls separately.

S1D. Marker gene expression per cell type (broad level 1 annotation) in Xenium data.

S1E. Scatter plot showing the correlation (Pearson R) between reported blast percentage and percentage of myeloid progenitor-like annotated in the Xenium data.

S1F. Scatter plot showing the correlation (Pearson R) between reported blast percentage and percentage of myeloid progenitor-like annotated in the IMC data.

S1G. Heatmap of cell composition per individual based on IMC data, showing Z scores of the percentage of the subpopulations.

S1H. Heatmap of cell composition per individual based on IMC data, showing Z scores of the absolute counts of the subpopulations, with patients in same order as frequency heatmap, demonstrating that analysis of cell frequencies or absolute cell counts display a highly similar trend.

S1I. UMAP of Xenium data, detailed (level 2) annotation.

S1J. Marker gene expression per cell type (detailed level 2 annotation) in Xenium data.

S1K. Spatial plot of a representative AML bone marrow sample (CA55), showing the spatial localization of the different stromal and endothelial cell types (Xenium).

S1L. Proportions of stromal and endothelial cell subtypes per age and condition group (Xenium). The pink box indicates significant enrichment compared to the same age group in the other condition (e.g. childhood control vs. childhood AML) according to the Mann-Whitney U test with BH-FDR correction.

S1M. Scatter plots showing the correlations (Spearman rho) between annotated cell type proportions (detailed level 2 annotation, Xenium) between different samples from the same individual, for those individuals with 2 samples available (n=26). Presented on a log10 scale, with the dotted lines representing the diagonals (perfect correlation).

S1N. Bar chart showing cell type proportions (detailed level 2 annotation, Xenium) per patient for whom a morphological (FAB) blast classification was available, sorted according to FAB class.

Supplemental Figure S2

S2A. Macrophage abundance between age and condition groups (IMC).

S2B. Macrophage abundance according to AML subtype (Xenium), no significant differences after Mann-Whitney U test with BH-FDR correction.

S2C. Gene expression (Xenium) of markers used for macrophage annotation, for all detailed (level 2) cell types.

S2D. T cell abundance as percentage of hematopoietic cells (IMC), no significant differences between age and condition groups after Mann-Whitney U test with BH-FDR correction.

S2E. T cell subtype abundance over the T/NK cell compartment per age and condition group (IMC).

S2F. T cell subtype abundance over all annotated cells, per age and condition group; * $P < 0.05$ after Mann-Whitney U test with BH-FDR correction.

S2G. Scatter plot showing the correlation between macrophage and T/NK cell proportions in the Xenium data per individual, colored by their immune category.

S2H. Box plots showing the percentage of annotated myeloid progenitor-like cells per immune category.

S2I. Bar plots showing the distribution of the immune categories over adult and pediatric AML patients.

S2J. Bar plots showing the distribution of the immune categories over the different AML subtypes, with the number of patients per subtype indicated above the bars.

S2K. Heatmap of the differential colocalization analysis for all detailed (level 2) cell types (Xenium); only those pairs with a significant stronger colocalization ($P<0.05$ according to CellCharter analysis) in either AML or controls are shown.

S2L. Heatmap of the differential colocalization analysis between AML subtypes and controls (Xenium); only those pairs with a significant stronger colocalization ($P<0.05$ according to CellCharter analysis) in either AML or controls are shown.

S2M. Heatmap of the differential colocalization analysis between the three AML immune categories and controls (Xenium); only those pairs with a significant stronger colocalization ($P<0.05$ according to CellCharter analysis) in either AML or controls are shown.

Supplemental Figure S3

S3A. Measure of stability of the identified cellular neighborhoods by CellCharter, for different numbers of cellular neighborhoods (Xenium), for the analysis using 1-hop neighbors.

S3B. Proportions of AML and controls per cellular neighborhood (Xenium), for the analysis using 1-hop neighbors.

S3C. Cellular composition of IMC-based cellular neighborhoods, for the analysis using 1-hop neighbors.

S3D. Comparison of Xenium-based cellular neighborhood proportions (with indication of AML subtype and immune category; same hierarchical ordering as Figure 3C; for the analysis using 1-hop neighbors) with IMC-based cellular neighborhood proportions (in the same order as the Xenium samples). Empty spaces indicate the samples that were not profiled using IMC.

S3E. Examples of Xenium-based neighborhoods, IMC images showing macrophage and T cell markers, and IMC-based neighborhoods for the same biopsies. Differences in location of bone trabecula between modalities are due to the fact that the biopsy sections used for the IMC and Xenium were either consecutive or a few micrometers apart.

S3F. T/NK cell subtypes per cellular neighborhood (Xenium), for the analysis using 1-hop neighbors.

S3G. Schematic depiction of the method of neighborhood detection by CellCharter for 1- and 2-hop neighbors: for each cell, the transcriptomes of the 1- and 2-hop neighbors were considered their neighborhood; the transcriptomes of these neighbors were aggregated across all cells and all biopsies and clustered into spatial clusters (cellular neighborhoods).

S3H. Measure of stability of the identified cellular neighborhoods by CellCharter, for different numbers of cellular neighborhoods (Xenium), for the analysis using 1- and 2-hop neighbors.

S3I. Proportions of AML and controls per cellular neighborhood (Xenium), for the analysis using 1- and 2-hop neighbors.

S3J. Cell type composition for the 10 cellular neighborhoods (CNs) identified by CellCharter (Xenium), for the analysis using 1- and 2-hop neighbors.

S3K. Proportions of CellCharter CNs per individual in relation to condition (AML vs. control) and to AML subtypes, with hierarchical clustering based on similarity of CN proportions, for the analysis using 1- and 2-hop neighbors.

S3L. Spatial plots showing examples of CN architecture in a control sample and 5 AML samples, for the analysis using 1- and 2-hop neighbors.

S3H. Measure of stability of the identified cellular neighborhoods by CellCharter, for different numbers of cellular neighborhoods (Xenium), for the analysis using 1-, 2- and 3-hop neighbors.

S3I. Proportions of AML and controls per cellular neighborhood (Xenium), for the analysis using 1-, 2- and 3-hop neighbors.

S3J. Cell type composition for the 10 cellular neighborhoods (CNs) identified by CellCharter (Xenium), for the analysis using 1-, 2- and 3-hop neighbors.

S3K. Proportions of CellCharter CNs per individual in relation to condition (AML vs. control) and to AML subtypes, with hierarchical clustering based on similarity of CN proportions, for the analysis using 1-, 2- and 3-hop neighbors.

S3L. Spatial plots showing examples of CN architecture in a control sample and 5 AML samples, for the analysis using 1-, 2- and 3-hop neighbors.

<u>Supplemental Figure S4</u>

S4A. All significant (P<0.05) predicted ligand-receptor interactions between macrophages, T/NK cells, and myeloid progenitor-like cells predicted by CellChat based on the Xenium and Flex data.

S4B. All significant (P<0.05) predicted pathways from myeloid progenitor-like cells (sender) to T/NK cells (receiver) per molecular AML subtype (Xenium); a blue dot indicates that the pathway was predicted for that subtype.

S4C. All significant (P<0.05) predicted pathways from macrophages (sender) to T/NK cells (receiver) per molecular AML subtype (Xenium); a blue dot indicates that the pathway was predicted for that subtype.

S4D. Significant (P<0.05) predicted specific ligand-receptor interactions of a selection of pathways from myeloid progenitor-like cells (sender) to T/NK cells (receiver) per molecular AML subtype (Xenium); a blue dot indicates that the interaction was predicted for that subtype.

S4E. Significant (P<0.05) predicted specific ligand-receptor interactions of a selection of pathways from macrophages (sender) to T/NK cells (receiver) per molecular AML subtype (Xenium); a blue dot indicates that the interaction was predicted for that subtype.

S4F. UMAP of the Flex data, with detailed level 2 annotation.

S4G. Bar plot showing the percentage of the level 1 annotation of both Flex and Xenium data, side by side for those samples with both modalities available.

S4H. Ligand and receptor gene expression in macrophages, myeloid progenitor-like cell subtypes and T cell subtypes of all ligand-receptor pairs identified by CellChat, for AML and controls separately. Asterisks indicate if a specific gene is expressed significantly higher (adjusted P<0.05) in AML (pink) or controls (blue) according to the differential gene expression analysis for that cell type.

S4I. Ligand and receptor gene expression in macrophages, myeloid progenitor-like cell subtypes and T cell subtypes of all ligand-receptor pairs identified by CellChat, for the three immune categories separately.

Supplemental Figure S5

S5A. Schematic overview of Xenium fusion probe design, showing that the padlock probe spans the fusion breakpoint.

S5B. CellChat predicted interaction strength per cell type (corrected for the population size: as if each cell type has the same cell number).

S5C. CellChat interaction strength per pathway for fusion+ and fusion- myeloid progenitor-like cells, with naturally occurring population size.

S5D. CellChat interaction strength per signaling pathway for fusion+ and fusion- myeloid progenitor-like cells, corrected for population size.

S5E. CellChat interaction strength per signaling pathway for fusion+ and fusion- immature myeloid cells, with naturally occurring population size.

S5F. CellChat interaction strength per signaling pathway for fusion+ and fusion- immature myeloid cells, corrected for population size.

Supplemental Figure S6

S6A. Schematic representation of the co-culture system between macrophages and naïve $CD4^+$ T cells.

S6B. Representative flow cytometry histograms showing CD25 and FOXP3 expression on naïve $CD4^+$ T cells cultured alone, in co-culture, or in a transwell condition with macrophages, in the presence or absence of PDX cells, for 2 days.

S6C. Quantification of CD25 and FOXP3 expression levels on naïve $CD4^+$ T cells cultured alone, in co-culture, or in transwell conditions with macrophages, in the presence or absence of PDX cells, for 2 days (n=3).

S6D. Percentage of CD25+FOXP3+ cells after 4 days of co-culture with macrophages and *KMT2A*-rearranged (*KMT2A*r) patient-derived xenograft (PDX) cells (left) or *RUNX1*::*RUNX1T1* PDX cells (right) at a 1:1 ratio (n=3).

S6E. Quantification of CD127 expression levels after 2 and 4 days of co-culture (n = 3).

S6F. Representative flow cytometry plots used as internal controls of the co-culture system, showing the induction of CD25 and FOXP3 on naïve $CD4^+$ T cells or on B cells after 2 and 4 days of co-culture with macrophages.

S6G. Relative mRNA expression levels of *LAG3*, *CTLA4* and *PD1* in naïve $CD4^+$ T cells after 2 and 4 days of co-culture with macrophages (n=3).

S6H. Percentage of iTregs after 2 and 4 days of co-culture with macrophages pre-incubated with *RUNX1*::*RUNX1T1* PDX cells (pM0) at a 1:1 ratio (n=3).

S6I. Quantification of CD25, FOXP3, and CD127 expression in iTregs generated from macrophages pre-incubated with *RUNX1T1* PDX cells (pM0), compared with M0 macrophage co-cultures, after 2 days at a 1:1 ratio (n=3).

S6J. CSFE mean fluorescence intensity of CD3+ effector T cells after 5 days co-culture with healthy CD3+ T cells, CD25+ Tregs, or macrophage-induced D4-iTregs (n=3).

Statistical analysis: one-way ANOVA. Significance is indicated as *P < 0.05; **P < 0.01; ***P < 0.001; ****P < 0.001.

<u>Supplemental Figure S7</u>

S7A. Cytokines expression levels after 2 days co-culture between macrophages and naïve T cells (M0+nT) or between macrophages, naïve T cells and *KMT2A*r PDX cells (M0+nT+KMT2Ar) or *RUNX1*::*RUNX1T1* PDX cells (M0+nT+RUNX1T1) (n=3).

S7B. Representative flow cytometry histograms showing CD31 and CXCR4 expression on naïve $CD4^+$ T cells and iTregs under the indicated conditions.

S7C. Quantification of TIM-3 and CD44 expression on naïve $CD4^+$ T cells cultured alone or after 2 days of co-culture with macrophages at a 1:1 ratio, in the presence or absence of *RUNX1*::*RUNX1T1* PDX cells (n = 3).

S7D. Quantification of TIM-3 expression on naïve $CD4^+$ T cells cultured alone or after 4 days of co-culture with macrophages at a 1:1 ratio, in the presence or absence of *KMT2A*r or *RUNX1T1* PDX cells (n = 3).

S7E. Quantification of CD44 expression on naïve $CD4^+$ T cells cultured alone or after 4 days of co-culture with macrophages at a 1:1 ratio, in the presence or absence of *KMT2A*r or *RUNX1*::*RUNX1T1* PDX cells (n = 3).

S7F. Representative flow cytometry histograms showing Galactin-9 expression on macrophages under the indicated conditions.

S7G. Quantification of Galectin-9 expression on macrophages after 2 and 4 days of monoculture or co-culture with naïve $CD4^+$ T cells, in the presence or absence of *KMT2A*r or *RUNX1*::*RUNX1T1* PDX cells (n=3).

S7H. ELISA assay measuring Galectin-9 cytokine level in culture supernatants under the indicated conditions (n=3).

S7I. Representative flow cytometry histogram showing CD25 and FOXP3 expression on naïve $CD4^+$ T cells culture alone or in co-culture with macrophages, in presence or absence of isotype controls (left). Percentage of $CD4^+CD25^+FOXP3^+$ cells under stated conditions (n=3).

S7J. Percentage of $CD4^+CD25^+FOXP3^+$ iTregs after 4 days of monoculture or co-culture with macrophages at a 1:1 ratio, in the presence of 10 µg/mL anti-CD44 and/or 10 µg/mL anti-TIM-3 blocking antibodies (n = 3).

S7K. Percentage of $CD4^+CD25^+$ cells after 2 days of co-culture with RUNX1::RUNX1T1 PDX cells and/or macrophages at a 1:1 ratio, in the presence of 10 µg/mL anti-CD44 and/or 10 µg/mL anti-TIM-3 blocking antibodies (n = 3).

Statistical analysis: one-way ANOVA. Significance is indicated as *P < 0.05; **P < 0.01; ***P < 0.001; ****P < 0.001.

<u>Supplemental Figure S8</u>

Example of several IMC stainings for sample CC5, on the slide that underwent Xenium (IMC post-Xenium) and the consecutive slide (True IMC signal), showing that the staining patterns for some antibodies were affected post-Xenium.
